# High performance single-cell gene regulatory network inference at scale: The Inferelator 3.0

**DOI:** 10.1101/2021.05.03.442499

**Authors:** Claudia Skok Gibbs, Christopher A Jackson, Giuseppe-Antonio Saldi, Andreas Tjärnberg, Aashna Shah, Aaron Watters, Nicholas De Veaux, Konstantine Tchourine, Ren Yi, Tymor Hamamsy, Dayanne M Castro, Nicholas Carriero, Bram L Gorissen, David Gresham, Emily R Miraldi, Richard Bonneau

## Abstract

**Motivation:** Gene regulatory networks define regulatory relationships between transcription factors and target genes within a biological system, and reconstructing them is essential for understanding cellular growth and function. Methods for inferring and reconstructing networks from genomics data have evolved rapidly over the last decade in response to advances in sequencing technology and machine learning. The scale of data collection has increased dramatically; the largest genome-wide gene expression datasets have grown from thousands of measurements to millions of single cells, and new technologies are on the horizon to increase to tens of millions of cells and above.

**Results:** In this work, we present the Inferelator 3.0, which has been significantly updated to integrate data from distinct cell types to learn context-specific regulatory networks and aggregate them into a shared regulatory network, while retaining the functionality of the previous versions. The Inferelator is able to integrate the largest single-cell datasets and learn cell-type specific gene regulatory networks. Compared to other network inference methods, the Inferelator learns new and informative *Saccharomyces cerevisiae* networks from single-cell gene expression data, measured by recovery of a known gold standard. We demonstrate its scaling capabilities by learning networks for multiple distinct neuronal and glial cell types in the developing *Mus musculus* brain at E18 from a large (1.3 million) single-cell gene expression dataset with paired single-cell chromatin accessibility data.

**Availability:** The inferelator software is available on GitHub (https://github.com/flatironinstitute/inferelator) under the MIT license and has been released as python packages with associated documentation (https://inferelator.readthedocs.io/).

## 1. Background

Gene expression is tightly regulated at multiple levels in order to control cell growth, development, and response to environmental conditions (Figure 1A). Transcriptional regulation is principally controlled by Transcription Factors (TFs) that bind to DNA and effect chromatin remodeling (Zaret, 2020) or directly modulate the output of RNA polymerases (Kadonaga, 2004). Three percent of *Saccharomyces cerevisiae* genes are TFs (Hahn and Young, 2011), and more than six percent of human genes are believed to be TFs or cofactors (Lambert *et al*., 2018). Connections between TFs and genes combine to form a transcriptional Gene Regulatory Network (GRN) that can be represented as a directed graph (Figure 1B). Learning the true regulatory network that connects regulatory TFs to target genes is a key problem in biology (Thompson *et al*., 2015; Chasman *et al*., 2016). Determining the valid GRN is necessary to explain how mutations that cause gene dysregulation lead to complex disease states (Hu *et al*., 2016), how variation at the genetic level leads to phenotypic variation (Mehta *et al*., 2021; Peter and Davidson, 2011), and how to re-engineer organisms to efficiently produce industrial chemicals and enzymes (Huang *et al*., 2017).

**Figure 1:**
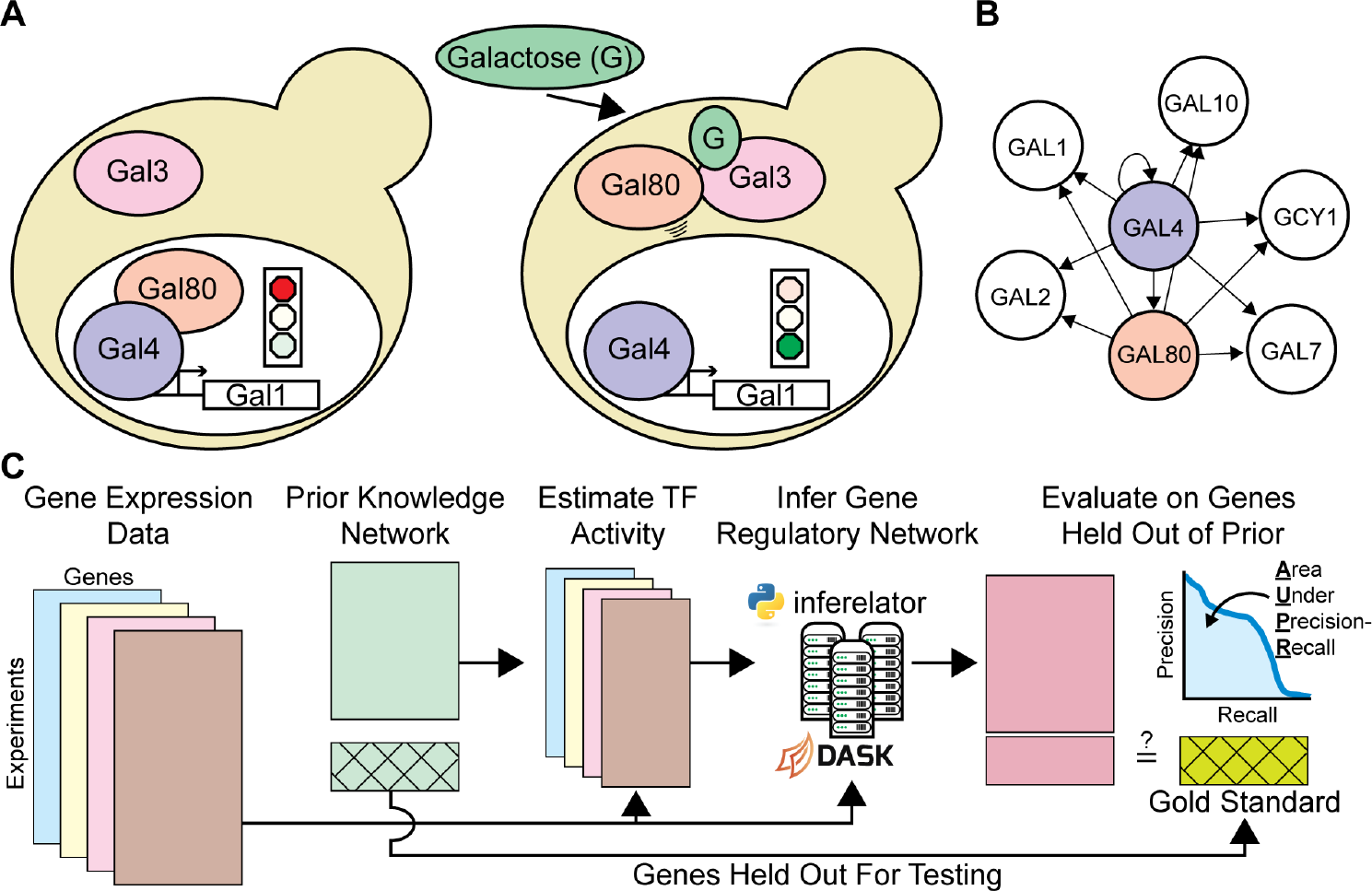
Learning Gene Regulatory Networks with the Inferelator (**A**) The response to the sugar galactose in *Saccharomyces cerevisiae* is mediated by the Gal4 and Gal80 TFs, a prototypical mechanism for altering cellular gene expression in response to stimuli. (**B**) Gal4 and Gal80 regulation represented as an unsigned directed graph connecting regulatory TFs to target genes. (**C**) Genome-wide Gene Regulatory Networks (GRNs) are inferred from gene expression data and prior knowledge about network connections using the Inferelator, and the resulting networks are scored by comparison with a gold standard of known interactions. A subset of genes are held out of the prior knowledge and used for evaluating performance.

Learning genome-scale networks relies on genome-wide expression measurements, initially captured with microarray technology (DeRisi *et al*., 1997), but today typically measured by RNA-sequencing (RNA-seq) (Nagalakshmi *et al*., 2008). A major difficulty is that biological systems have large numbers of both regulators and targets, and many regulators are redundant or interdependent. Many plausible networks can explain observed expression data and the regulation of gene expression (Szederkényi *et al*., 2011), which makes identifying the correct network challenging. Designing experiments to produce data that increases network identifiability is possible (Ud-Dean and Gunawan, 2016), but most data is collected for specific projects and repurposed for network inference as a consequence of the cost of data collection. Large-scale experiments in which a perturbation is made and dynamic data is collected over time is exceptionally useful for learning GRNs but systematic studies that collect this data are rare (Hackett *et al*., 2020).

Measuring the expression of single cells using single-cell RNA-sequencing (scRNAseq) is an emerging and highly scalable technology. Microfluidic-based single-cell techniques (Macosko *et al*., 2015; Zilionis *et al*., 2017; Zheng *et al*., 2017) allow for thousands of measurements in a single experiment. Split-pool barcoding techniques (Rosenberg *et al*., 2018) are poised to increase single-cell throughput by an order of magnitude. These techniques have been successfully applied to generate multiplexed gene expression data from pools of barcoded cell lines with loss-of-function TF mutants (Dixit *et al*., 2016; Jackson *et al*., 2020), enhancer perturbations (Schraivogel *et al*., 2020), and disease-causing oncogene variants (Ursu *et al*., 2020). Individual cell measurements are sparser and noisier than measurements generated using traditional RNA-seq, although in aggregate the gene expression profiles of single-cell data match RNA-seq data well (Svensson, 2020), and techniques to denoise single-cell data have been developed (Arisdakessian *et al*., 2019; Tjärnberg *et al*., 2021).

The seurat (Stuart *et al*., 2019) and scanpy (Wolf *et al*., 2018) bioinformatics toolkits are established tools for single-cell data analysis, but pipelines for inferring GRNs from single-cell data are still nascent, although many are under development (Zappia and Theis, 2021). Recent work has begun to systematically benchmarking network inference tools, and the BEELINE (Pratapa *et al*., 2020) and other (Nguyen *et al*., 2021; Chen and Mar, 2018) benchmarks have identified promising methods. Testing on real-world data has proved difficult, as reliable gold standard networks for higher eukaryotes do not exist. scRNAseq data for microbes which have some known ground truth networks (like *Saccharomyces cerevisiae* and *Bacillus subtilis*) was not collected until recently. As a consequence, most computational method benchmarking has been done using simulated data. Finally, GRN inference is computationally challenging, and the most scalable currently-published GRN pipeline has learned GRNs from 50,000 cells of gene expression data (Van de Sande *et al*., 2020).

Here we describe the Inferelator 3.0 pipeline for single-cell GRN inference, based on regularized regression (Bonneau *et al*., 2006). This pipeline calculates TF activity (Ma and Brent, 2021) using a prior knowledge network and regresses scRNAseq expression data against that activity estimate to learn new regulatory edges. We compare it directly to two other network inference methods that also utilize prior network information and scRNAseq data, benchmarking using real-world *Saccharomyces cerevisiae* scRNAseq data and comparing to a high-quality gold standard network. The first comparable method, SCENIC (Van de Sande *et al*., 2020), is GRN inference pipeline that estimates the importance of TFs in explaining gene expression profiles and then constrains this correlative measure with prior network information to identify regulons. The second comparable method, CellOracle (Kamimoto *et al*., 2020), has been recently proposed as a pipeline to integrate single-cell ATAC and expression data using a motif-based search for potential regulators, followed by bagging Bayesian ridge regression to enforce sparsity in the output GRN.

Older versions of the Inferelator (Madar *et al*., 2009) have performed well inferring networks for *Bacillus subtilis* (Arrieta-Ortiz *et al*., 2015), human Th17 cells (Ciofani *et al*., 2012; Miraldi *et al*., 2019), mouse lymphocytes (Pokrovskii *et al*., 2019), *Saccharomyces cerevisiae* (Tchourine *et al*., 2018), and *Oryza sativa* (Wilkins *et al*., 2016). We have implemented the Inferelator 3.0 with new functionality in python to learn GRNs from scRNAseq data. Three different model selection methods have been implemented: a Bayesian best-subset regression method (Greenfield *et al*., 2013), a StARS-LASSO (Miraldi *et al*., 2019) regression method in which the regularization parameter is set by stability selection (Liu *et al*., 2010), and a multitask-learning regression method (Castro *et al*., 2019). This new package provides scalability, allowing millions of cells to be analyzed together, as well as integrated support for multi-task GRN inference, while retaining the ability to utilize bulk gene expression data. We show that the Inferelator 3.0 is a state-of-the-art method by testing against SCENIC and CellOracle on model organisms with reliable ground truth networks, and show that the Inferelator 3.0 can generate a mouse neuronal GRN from a publicly available dataset containing 1.3 million cells.

## 2. Results

### 2.1. The Inferelator 3.0

In the 12 years since the last major release of the Inferelator (Madar *et al*., 2009), the scale of data collection in biology has accelerated enormously. We have therefore rewritten the Inferelator as a python package to take advantage of the concurrent advances in data processing. For inference from small scale gene expression datasets (*<* 10^4^ observations), the Inferelator 3.0 uses native python multiprocessing to run on individual computers. For inference from extremely large scale gene expression datasets (*>* 10^4^ observations) that are increasingly available from scRNAseq experiments, the Inferelator 3.0 takes advantage of the Dask analytic engine (Rocklin, 2015) for deployment to high-performance clusters (Figure 1C), or for deployment as a kubernetes image to the Google cloud computing infrastructure.

### 2.2. Network Inference using Bulk RNA-Seq Expression Data

We incorporated several network inference model selection methods into the Inferelator 3.0 (Figure 2A) and evaluate their performance on the prokaryotic model *Bacillus subtilis* and the eukaryotic model *Saccharomyces cerevisiae*. Both *B. subtilis* (Arrieta-Ortiz *et al*., 2015; Nicolas *et al*., 2012) and *S. cerevisiae* (Tchourine *et al*., 2018; Hackett *et al*., 2020) have large bulk RNA-seq and microarray gene expression datasets, in addition to a relatively large number of experimentally determined TF-target gene interactions that can be used as a gold standard for assessing network inference. Using two independent datasets for each organism, we find that the model selection methods Bayesian Best Subset Regression (BBSR) (Greenfield *et al*., 2010) and Stability Approach to Regularization Selection for Least Absolute Shrinkage and Selection Operator (StARS-LASSO) (Miraldi *et al*., 2019) perform equivalently (Figure 2B). The Inferelator performs substantially better than a network inference method (GRNBOOST2) that does not use prior network information (Figure 2B; dashed blue lines).

**Figure 2:**
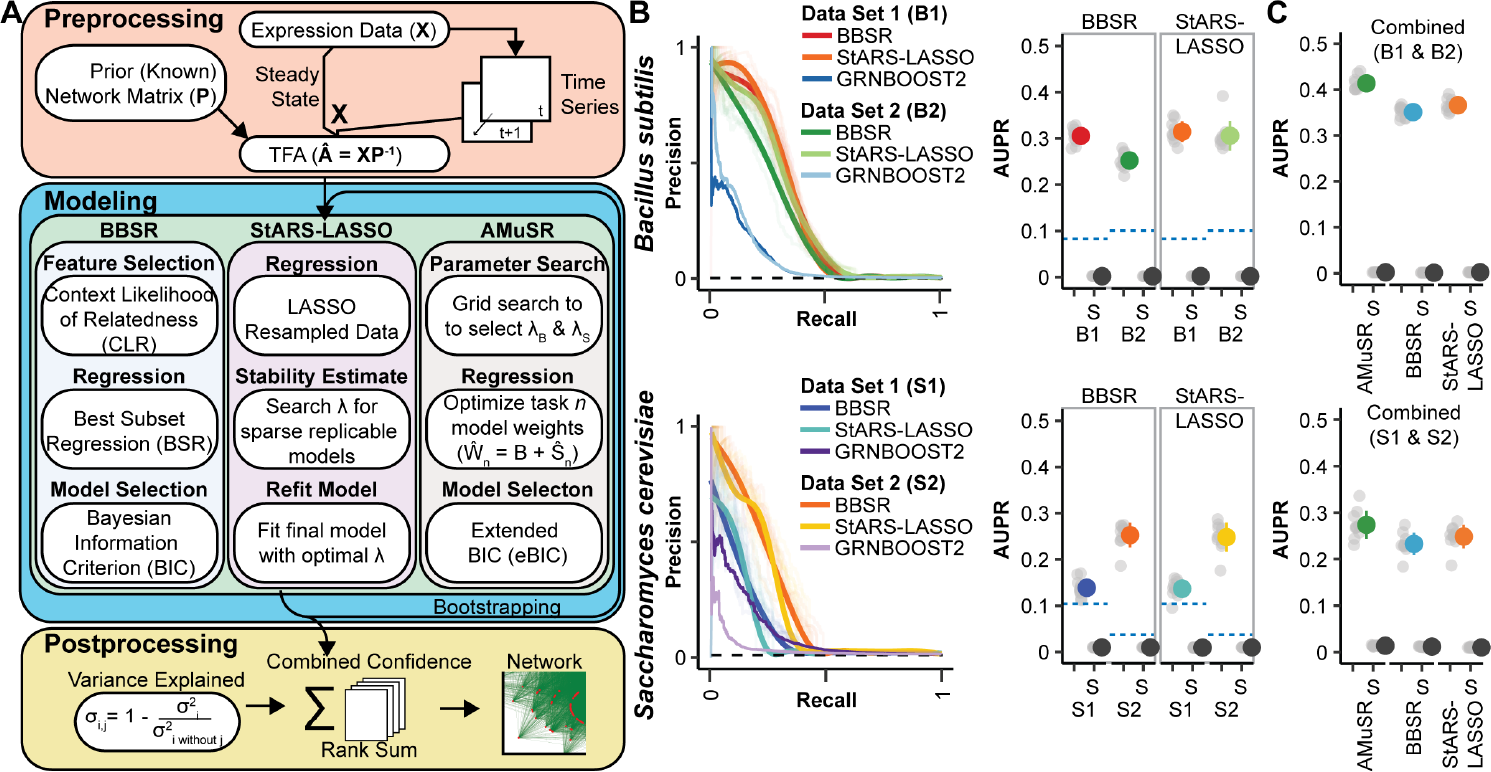
Network Inference Performance on Multiple Model Organism Datasets (**A**) Schematic of Inferelator workflow and a brief summary of the differences between GRN model selection methods (**B**) Results from 10 replicates of GRN inference for each modeling method on (**i**) *Bacillus subtilis* GSE67023 (B1), GSE27219 (B2) and (**ii**) *Saccharomyces cerevisiae* GSE142864 (S1), and Tchourine *et al*. (2018) (S2). Precision-recall curves are shown for replicates where 20% of genes are held out of the prior and used for evaluation, with a smoothed consensus curve. The black dashed line on the precision-recall curve is the expected random performance based on random sampling from the gold standard. AUPR is plotted for each cross-validation result in gray, with mean *±* standard deviation in color. Experiments labeled with (S) are shuffled controls, where the labels on the prior adjacency matrix have been randomly shuffled. 10 shuffled replicates are shown as gray dots, with mean *±* standard deviation in black. The blue dashed line is the the performance of the GRNBOOST2 network inference algorithm, which does not use prior network information, scored against the entire gold standard network. (**C**) Results from 10 replicates of GRN inference using two datasets as two network inference tasks on *Bacillus subtilis* and (ii) *Saccharomyces cerevisiae*. AMuSR is a multi-task learning method; BBSR and StARS-LASSO are run on each task separately and then combined into a unified GRN. AUPR is plotted as in **B**.

The two independent data sets show clear batch effects (Supplemental Figure 1A), and combining them for network inference is difficult; conceptually, each dataset is in a separate space, and must be mapped into a shared space. We take a different approach to addressing the batch effects between datasets by treating them as separate learning tasks (Castro *et al*., 2019) and then combining network information into a unified GRN. This results in a considerable improvement in network inference performance over either dataset individually (Figure 2C). The best performance is obtained with Adaptive Multiple Sparse Regression (AMuSR) (Castro *et al*., 2019), a multi-task learning method that shares information between tasks during regression. The GRN learned with AMuSR explains the variance in the expression data better than learning networks from each dataset individually with BBSR or StARS-LASSO and then combining them (Supplemental Figure 1B), and retains a common network core across different tasks (Supplemental Figure 1C).

### 2.3. Generating Prior Networks from Chromatin Data and Transcription Factor Motifs

The Inferelator 3.0 produces an inferred network from a combination of gene expression data and a prior knowledge GRN constructed from existing knowledge about known gene regulation. Curated databases of regulator-gene interactions culled from domain-specific literature are an excellent source for prior networks. While some model systems have excellent databases of known interactions, these resources are unavailable for most organisms or cell types. In these cases, using chromatin accessibility determined by a standard Assay for Transposase-Accessible Chromatin (ATAC) in combination with the known DNA-binding preferences for TFs to identify putative target genes is a viable alternative (Miraldi *et al*., 2019).

To generate these prior networks we have developed the inferelator-prior accessory package that uses TF motif position-weight matrices to score TF binding within gene regulatory regions and build sparse prior networks (Figure 3A). These gene regulatory regions can be identified by ATAC, by existing knowledge from TF Chromatin Immunoprecipitation (ChIP) experiments, or from known databases (e.g. ENCODE (ENCODE Project Consortium *et al*., 2020)). Here, we compare the inferelator-prior tool to the CellOracle package (Kamimoto *et al*., 2020) that also constructs motif-based networks that can be constrained to regulatory regions, in *Saccharomyces cerevisiae* by using sequences 200bp upstream and 50bp downstream of each gene TSS as the gene regulatory region. The inferelator-prior and CellOracle methods produce networks that are similar when measured by Jaccard index but are dissimilar to the YEASTRACT literature-derived network (Figure 3B). These motif-derived prior networks from both the inferelator-prior and CellOracle methods perform well as prior knowledge for GRN inference using the Inferelator 3.0 pipeline (Figure 3C). The source of the motif library has a significant effect on network output, as can be seen with the well-characterized TF GAL4. GAL4 has a canonical *CGGN*_11_*CGG* binding site; different motif libraries have different annotated binding sites (Supplemental Figure 2A) and yield different motif-derived networks with the inferelatorprior pipeline (Supplemental Figure 2B-C).

**Figure 3:**
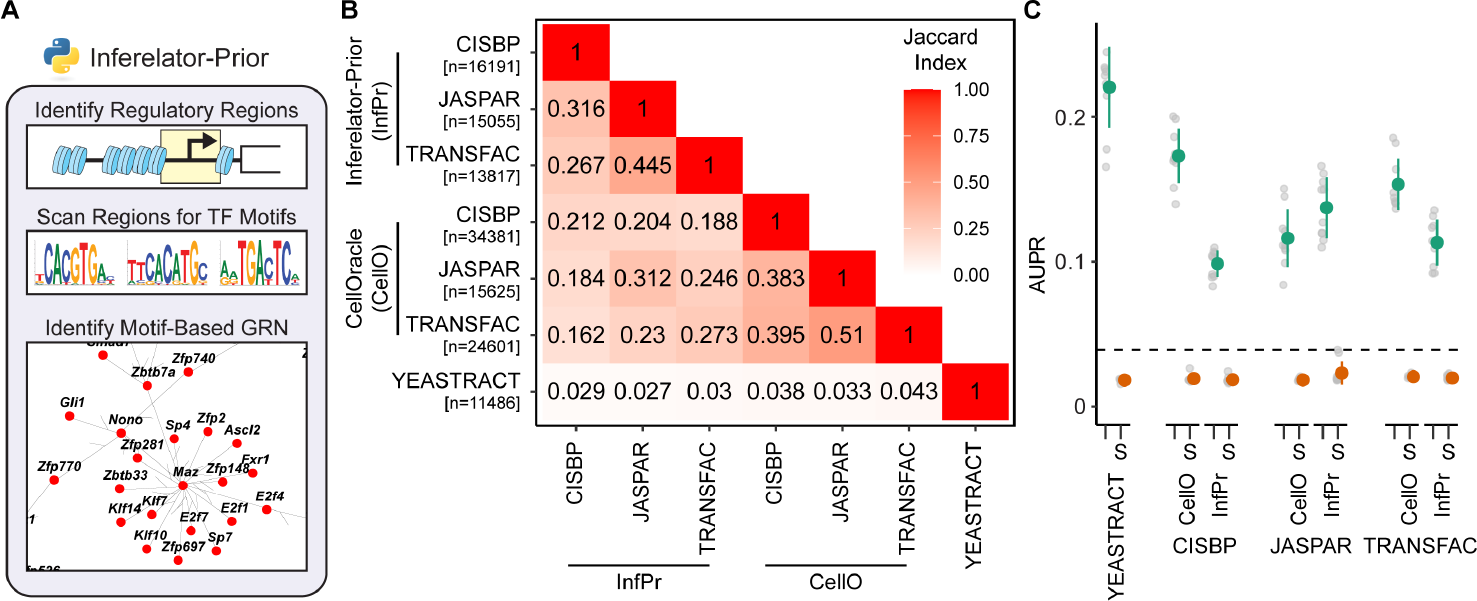
Construction and Performance of Network Connectivity Priors Using TF Motif Scanning (**A**) Schematic of inferelator-prior workflow, scanning identified regulatory regions (e.g. by ATAC) for TF motifs to construct adjacency matrices (**B**) Jaccard similarity index between *Saccharomyces cerevisiae* prior adjacency matrices generated by the inferelator-prior package, by the CellOracle package, and obtained from the YEASTRACT database. Prior matrices were generated using TF motifs from the CIS-BP, JASPAR, and TRANSFAC databases with each pipeline (n is the number of edges in each prior adjacency matrix). (**C**) The performance of Inferelator network inference using each motif-derived prior. Performance is evaluated by AUPR, scoring against genes held out of the prior adjacency matrix, based on inference using 2577 genome-wide microarray experiments. Experiments labeled with (S) are shuffled controls, where the labels on the prior adjacency matrix have been randomly shuffled. The black dashed line is the performance of the GRNBOOST2 algorithm, which does not incorporate prior knowledge, scored against the entire gold standard network.

### 2.4. Network Inference using Single-Cell Expression Data

Single-cell data is undersampled and noisy, but large numbers of observations are collected in parallel. As network inference is a population-level analysis which must already be robust against noise, we reason that data preprocessing that improves per-cell analyses (like imputation) is unnecessary. We test this by quantitatively evaluating networks learned from *Saccharomyces cerevisiae* scRNAseq data (Jackson *et al*., 2020; Jariani *et al*., 2020) with a previously-defined yeast gold standard (Tchourine *et al*., 2018). This expression data is split into 15 separate tasks, based on labels that correspond to experimental conditions from the original works (Figure 4A). A network is learned for each task separately using the YEASTRACT literature-derived prior network, from which a subset of genes are withheld, and aggregated into a final network for scoring on held-out genes from the gold standard. We test a combination of several preprocessing options with three network inference model selection methods (Figure 4B-D).

**Figure 4:**
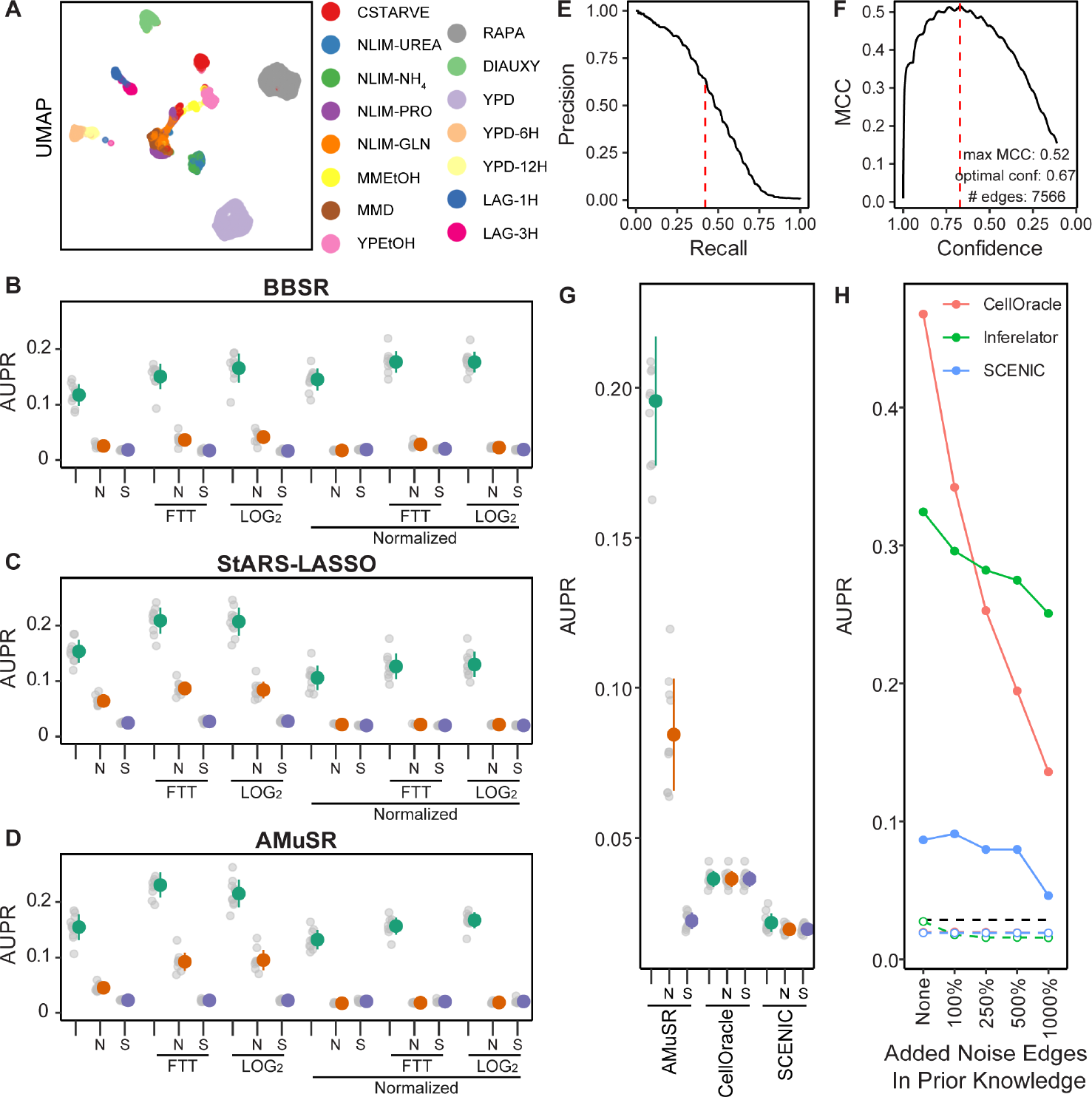
Network Inference Performance Using *Saccharomyces cerevisiae* Single-Cell Data (**A**) Uniform Manifold Approximation and Projection (UMAP) plot of yeast scRNAseq data, colored by the experimental grouping of individual cells (tasks). (**B**) The effect of preprocessing methods on network inference using BBSR model selection on 14 task-specific expression datasets, as measured by AUPR. Colored dots represent mean *±* standard deviation of all replicates. Data is either untransformed (raw counts), transformed by Freeman-Tukey Transform (FTT), or transformed by *log*2(*x*1) pseudocount. Non-normalized data is compared to data normalized so that all cells have identical count depth. Network inference performance is compared to two baseline controls; data which has been replaced by Gaussian noise (N) and network inference using shuffled labels in the prior network (S). (**C**) Performance evaluated as in **B** on StARS-LASSO model selection. (**D**) Performance evaluated as in **B** on AMuSR model selection. (**E**) Precision-recall of a network constructed using FTT-transformed, non-normalized AMuSR model selection, as determined by the recovery of the prior network. Dashed red line is the retention threshold identified by Matthews Correlation Coefficient. (**F**) Matthews Correlation Coefficient (MCC) of the same network as in **E**. Dashed red line is the confidence score of the maximum MCC. (**G**) Performance evaluated as in **B** comparing the Inferelator (FTTtransformed, non-normalized AMuSR) against the SCENIC and CellOracle network inference pipelines. (**H**) Performance of the Inferelator (FTT-transformed, non-normalized AMuSR) compared to SCENIC and CellOracle without holding genes out of the prior knowledge network. Additional edges are added randomly to the prior knowledge network as a percentage of the true edges in the prior. Colored dashed lines represent controls for each method where the labels on the prior knowledge network are randomly shuffled. The black dashed line represents performance of the GRNBOOST2 algorithm, which identifies gene adjacencies as the first part of the SCENIC pipeline without using prior knowledge.

We find that network inference is generally sensitive to the preprocessing options chosen, and that this effect outweighs the differences between different model selection methods (Figure 4B-D). A standard Freeman-Tukey or log_2_ pseudocount transformation on raw count data yields the best performance, with notable decreases in recovery of the gold standard when count data is count depth-normalized (such that each cell has the same total transcript counts). The performance of the randomly generated Noise control (N) is higher than the performance of the shuffled (S) control when counts per cell are not normalized, suggesting that total counts per cell provides additional information during inference.

Different model performance metrics, like AUPR, Matthews Correlation Coefficient (MCC), and F1 score correlate very well and identify the same optimal hyperparameters (Supplemental Figure 4). We apply AMuSR to data that has been Freeman-Tukey transformed to generate a final network without holding out genes for cross-validation (Figure 4E). While we use AUPR as a metric for evaluating model performance, selecting a threshold for including edges in a GRN by precision or recall requires a target precision or recall to be chosen arbitrarily. Choosing the Inferelator confidence score threshold to include the edges in a final network that maximize MCC is a simple heuristic to select the size of a learned network that maximizes overlap with another network (e.g. a prior knowledge GRN or gold standard GRN) while minimizing links not in that network (Figure 4F). Maximum F1 score gives a less conservative GRN as true negatives are not considered and will not diminish the score. Both metrics balance similarity to the test network with overall network size, and therefore represent straightforward heuristics that do not rely on arbitrary thresholds.

In order to determine how the Inferelator 3.0 compares to similar network inference tools, we apply both CellOracle and SCENIC to the same network inference problem, where a set of genes are held out of the prior knowledge GRN and used for scoring. We see that the Inferelator 3.0 can make predictions on genes for which no prior information is known, but CellOracle and SCENIC cannot (Figure 4G). When provided with a complete prior knowledge GRN, testing on genes which are not held out, CellOracle outperforms the Inferelator, although the Inferelator is more robust to noise in the prior knowledge GRN (Figure 4H). This is a key advantage, as motif-generated prior knowledge GRNs are expected to be noisy.

### 2.5. Large-scale Single-Cell Mouse Neuron Network Inference

The Inferelator 3.0 is able to distribute work across multiple computational nodes, allowing networks to be rapidly learned from *>* 10^5^ cells (Supplemental Figure 5A). We show this by applying the Inferelator to a large (1.3 million cells of scRNAseq data), publicly available dataset of mouse brain cells (10x genomics) that is accompanied by 15,000 single-cell ATAC (scATAC) measurements. We separate the expression and scATAC data into broad categories; Excitatory neurons, Interneurons, Glial cells and Vascular cells (Figure 5A-E). After initial quality control, filtering, and cell type assignment, 766,402 scRNAseq and 7,751 scATAC observations remain (Figure 5F, Supplemental Figure 5B-D).

**Figure 5:**
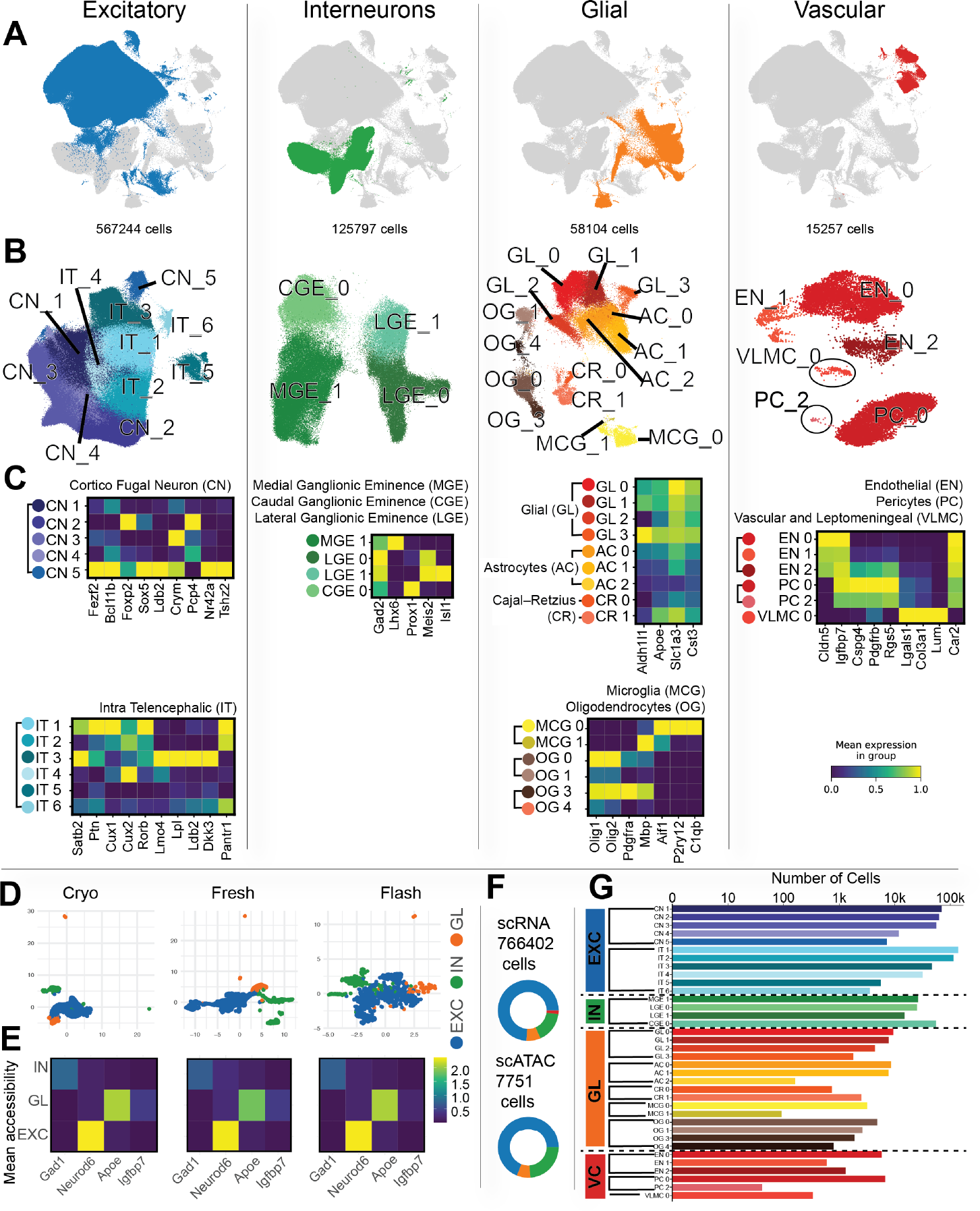
Processing Large Single-Cell Mouse Brain Data for Network Inference (**A**) UMAP plot of all mouse brain scRNAseq data with Excitatory neurons, Interneurons, Glial cells and Vascular cells colored. (**B**) UMAP plot of cells from each broad category colored by louvain clusters and labeled by cell type. (**C**) Heatmap of normalized gene expression for marker genes that distinguish cluster cell types within broad categories. (**D**) UMAP plot of mouse brain scATAC data with Excitatory neurons, Interneurons, and Glial cells colored. (**E**) Heatmap of normalized mean gene accessibility for marker genes that distinguish broad categories of cells. (**F**) The number of scRNA-seq and scATAC cells in each of the broad categories. (**G**) The number of scRNA-seq cells in each cell type specific cluster.

scRNAseq data is further clustered within broad categories into clusters (Figure 5B) that are assigned to specific cell types based on marker expression (Figure 5C, Supplemental Figure 6). scATAC data is aggregated into chromatin accessibility profiles for Excitatory neurons, Interneurons, and Glial cells (Figure 5D) based on accessibility profiles (Figure 5E), which are then used with the TRANSFAC mouse motif position-weight matrices to construct prior knowledge GRNs with the inferelator-prior pipeline. Most scRNAseq cell type clusters have thousands of cells, however rare cell type clusters are smaller (Figure 5G) After processing scRNAseq into 36 cell type clusters and scATAC data into 3 broad (Excitatory neurons, Interneurons, and Glial) prior GRNs, we used the Inferelator 3.0 to learn an aggregate mouse brain GRN. Each of the 36 clusters was assigned the most appropriate of the three prior GRNs and learned as a separate task using the AMuSR model selection framework. The resulting aggregate network contains 20,991 TF gene regulatory edges, selected from the highest confidence predictions to maximize MCC (Figure 6A-B). A common regulatory core of 1,909 network edges is present in every task-specific network (Figure 6C). Task-specific networks from similar cell types tend to be highly similar, as measured by Jaccard index (Figure 6D). We learn very similar GRNs from each excitatory neuron task, and very similar GRNs from each interneuron task, although each of these broad categories yields different regulatory networks. There are also notable examples where glial and vascular tasks produce GRNs that are distinctively different from other glial and vascular GRNs.

**Figure 6:**
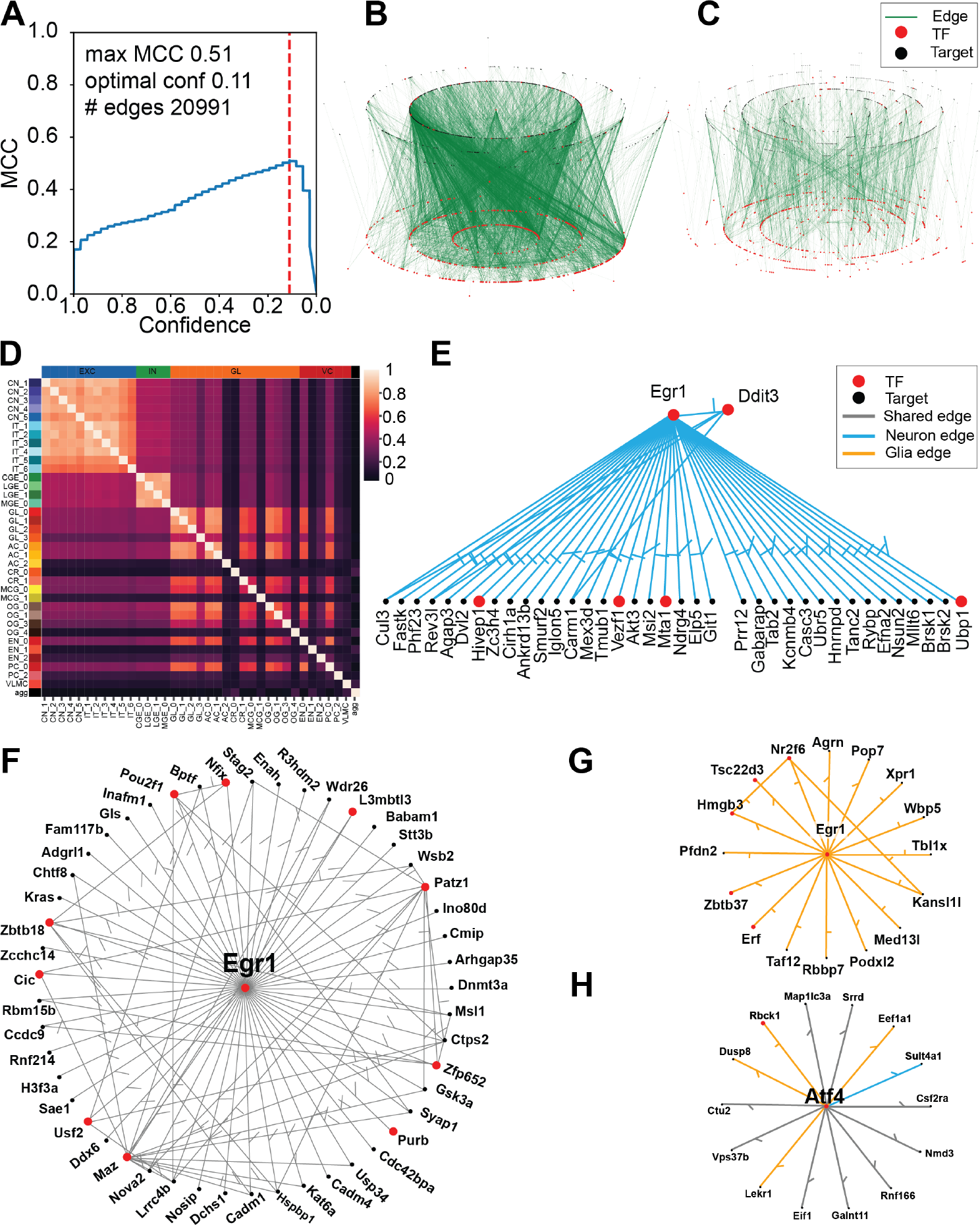
Learned GRN For The Mouse Brain (**A**) MCC for the aggregate network based on Inferelator prediction confidence. The dashed line shows the confidence score which maximizes MCC. Network edges at and above this line are retained in the final network. (**B**) Aggregate GRN learned. (**C**) Network edges which are present in every individual task. (**D**) Jaccard similarity index between each task network (**E**) Network targets of the *EGR1* TF in neurons. (**F**) Network targets of the *EGR1* TF in both neurons and glial cells. (**G**) Network targets of the *EGR1* TF in glial cells. (**H**) Network of the *ATF4* TF where blue edges are neuron specific, orange edges are glial specific, and black edges are present in both categories.

Finally, we can examine specific TFs and compare networks between cell type categories (Supplemental Figure 7). The TFs Egr1 and Atf4 are expressed in all cell types and Egr1 is known to have an active role at embryonic day 18 (E18) (Sun *et al*., 2019). In our learned network, Egr1 targets 103 genes, of which 20 are other TFs (Figure 6E-G). Half of these targets (49) are common to both neurons and glial cells, while 38 target genes are specific to neuronal GRNs and 16 target genes are specific to glial GRNs. We identify 14 targets for Atf4 (Figure 6H), the majority of which (8) are common to both neurons and glial cells, with only 1 target gene specific only to neuronal GRNs and 5 targets specific only to glial GRNs.

## 3. Discussion

We have developed the Inferelator 3.0 software package to scale to match the size of any network inference problem, with no organism-specific requirements that preclude easy application to non-mammalian organisms. Model baselines can be easily established by shuffling labels or generating noised data sets, and cross-validation and scoring on holdout genes is built directly into the pipeline. We believe this is particularly important as evaluation of single-cell network inference tools on real-world problems has lagged behind the development of inference methods themselves. Single-cell data collection has focused on complex higher eukaryotes and left the single-cell network inference field bereft of reliable standards to test against. Recent collection of scRNAseq data from traditional model organisms provides an opportunity to identify successful and unsuccessful strategies for network inference. For example, we find that performance differences between our methods of model selection may be smaller than differences caused by data cleaning and preprocessing. Benchmarking using model organism data should be incorporated in all single-cell method development, as it mitigates cherry-picking from complex network results and can prevent use of flawed performance metrics which are the only option when no reliable gold standard exists. In organisms without a reliable gold standard, network inference predictions should not be assumed correct and must be validated experimentally (Allaway *et al*., 2021).

Unlike traditional RNA-seq that effectively measures the average gene expression of large number of cells, scRNAseq can yield individual measurements for many different cell types that are implementing distinct regulatory programs. Learning GRNs from each of these cell types as a separate learning task in a multi-task framework allows cell type differences to be retained, while still taking advantage of the common regulatory programs. We demonstrate the use of this multi-task approach to simultaneously learn regulatory GRNs for a variety of mouse neuronal cell types from a very large (10^6^) single-cell data set. This includes learning GRNs for rare cell types; by sharing information between cell types during regression, we are able to learn a core regulatory network while also retaining cell type specific interactions. As the GRNs that have been learned for each cell type are sparse and consist of the highest-confidence regulatory edges, they are very amenable to exploration and experimental validation.

A number of limitations remain that impact our ability to accurately predict gene expression and cell states. Most important is a disconnect between the linear modeling that we use to learn GRNs and the non-linear biophysical models that incorporate both transcription and RNA decay. Modeling strategies that more accurately reflect the underlying biology will improve GRN inference directly, and will also allow prediction of useful latent parameters (e.g. RNA half-life) that are experimentally difficult to access. It is also difficult to determine if regulators are activating or repressing specific genes (Kamimoto *et al*., 2020), complicated further by biological complexity that allows TFs to switch between activation and repression (Papatsenko and Levine, 2008). Improving prediction of the directionality of network edges, and if directionality is stable in different contexts would also be a major advance. Many TFs bind cooperatively as protein complexes, or antagonistically via competitive binding, and explicit modeling of these TF-TF interactions would also improve GRN inference and make novel biological predictions. The modular Inferelator 3.0 framework will allow us to further explore these open problems in regulatory network inference without having to repeatedly reinvent and reimplement existing work. We expect this to be a valuable tool to build biologically-relevant GRNs for experimental followup, as well as a baseline for further development of computational methods in the network inference field.

## 4. Methods

Additional methods available in Supplemental Methods

### 4.1. Network Inference in Bacillus subtilis

Microarray expression data for *Bacillus subtilis* was obtained from NCBI GEO; GSE67023 (Arrieta-Ortiz *et al*., 2015) (n=268) and GSE27219 (Nicolas *et al*., 2012) (n=266). GRNs were learned using each expression dataset separately in conjunction with a known prior network (Arrieta-Ortiz *et al*., 2015) (Supplemental Data 1). Performance was evaluated by AUPR on ten replicates by holding 20% of the genes in the known prior network out, learning the GRN, and then scoring based on the held-out genes. Baseline shuffled controls were performed by randomly shuffling the labels on the known prior network.

Multi-task network inference uses the same *B. subtilis* prior for both tasks, with 20% of genes held out for scoring. Individual task networks are learned and rank-combined into an aggregate network. Performance was evaluated by AUPR on the held-out genes.

### 4.2. Network Inference in Saccharomyces cerevisiae

A large microarray dataset was obtained from NCBI GEO and normalized for a previous publication (Tchourine *et al*., 2018) (n=2,577; 10.5281/zenodo.3247754). In short, this data was preprocessed with limma (Ritchie *et al*., 2015) and quantile normalized. A second microarray dataset consisting of a large dynamic perturbation screen (Hackett *et al*., 2020) was obtained from NCBI GEO accession GSE142864 (n=1,693). This dataset is the median of three replicate *log*_2_ fold changes of an experimental channel over a control channel (which is the same for all observations). The *log*_2_ fold change is further corrected for each time course by subtracting the *log*_2_ fold change observed at time 0. GRNs were learned using each expression dataset separately in conjunction with a known YEASTRACT prior network (Teixeira *et al*., 2018; Monteiro *et al*., 2020) (Supplemental Data 1). Performance was evaluated by AUPR on ten replicates by holding 20% of the genes in the known prior network out, learning the GRN, and then scoring based on the held-out genes in a separate gold standard (Tchourine *et al*., 2018). Baseline shuffled controls were performed by randomly shuffling the labels on the known prior network.

Multi-task network inference uses the same YEASTRACT prior for both tasks, with 20% of genes held out for scoring. Individual task networks are learned and rank-combined into an aggregate network, which is then evaluated by AUPR on the held-out genes in the separate gold standard.

### 4.3. Single-Cell Network Inference in Saccharomyces cerevisiae

Single-cell expression data for *Saccharomyces cerevisiae* was obtained from NCBI GEO (GSE125162 (Jackson *et al*., 2020) and GSE144820 (Jariani *et al*., 2020)). Individual cells (n=44,343) are organized into one of 14 groups based on experimental metadata and used as separate tasks in network inference. Genes were filtered such that any gene with fewer than than 2217 total counts in all cells (1 count per 20 cells) was removed. Data was used as raw, unmodified counts, was Freeman-Tukey transformed (*^√^x* + 1 + *^√^x −* 1), or was log_2_ pseudocount transformed (log_2_(*x* + 1)). Data was either not normalized, or depth normalized by scaling so that the sum of all counts for each cell is equal to the median of the sum of counts of all cells. For each set of parameters, network inference is run 10 times, using the YEASTRACT network as prior knowledge with 20% of genes held out for scoring. For noise-only controls, gene expression counts are simulated randomly such that for each gene *i*, *x_i_ ∼ N* (*µ_xi_ , σ_xi_* ) and the sum for each cell is equal to the sum in the observed data. For shuffled controls, the gene labels on the prior knowledge network are randomly shuffled.

### 4.4. Single-Cell Network Inference in Mus musculus neurons

GRNs were learned using AMuSR on log_2_ pseudocount transformed count data for each of 36 cell type specific clusters as separate tasks with the appropriate prior knowledge network. An aggregate network was created by rank-summing each cell type GRN. MCC was calculated for this aggregate network based on a comparison to the union of the three prior knowledge networks, and the confidence score which maximized MCC was selected as a threshold to determine the size of the final network. Neuron specific edges were identified by aggregating filtered individual task networks with their respective confidence score to maximize MCC. Each edge that was shared with a glial or vascular network was excluded. The remaining neuron specific edges are interneuron specific, excitatory specific or shared.

## Supporting information

Supplemental Data 1 (.tar.gz)

Supplemental Data 2 (.tar.gz)

## Acknowledgements Funding

This work was supported by the NSF [MCB-1818234, IOS-1546218], the NIH [R35GM122515, R01HD096770, R01NS116350, R01NS118183, R01GM107466, R01GM134066, R01AI140766, R01AI130945, U01AI150748, R01AI153442, RM1HG011014, U54AG076040], and the Simons Foundation.

## Acknowledgements

We thank past and present members of the Gresham, Miraldi, and Bonneau labs for discussions and valuable feedback on this manuscript. We also thank the staff of the Flatiron Institute Scientific Computing Core for their tireless efforts to build and maintain the High Performance Computing resources which we rely on. This work was supported in part through the NYU IT High Performance Computing resources, services, and staff expertise.

## 1. Supplemental Methods

### 1.1. BEELINE Benchmarks

Test data and networks for the BEELINE panel were obtained from Zenodo (DOI: 10.5281/zenodo.3378975). For tests without any prior network information, the Inferelator was provided with expression data and scored against the entire gold-standard network. For tests with prior information, the Inferelator was provided with expression data and half the genes from the gold-standard network as a prior knowledge network. Scoring was performed on genes which were not provided in the prior knowledge network. Network inference was performed on each of expression data sets 10 times, with different random seeds each time. The median AUPR of the 10 network inference runs is reported as the performance for that specific expression data set. AUPR ratios are calculated using the baseline AUPR as defined in the BEELINE benchmarks. Scores for other methods are taken from supplemental data of the previously published BEELINE benchmark.

### 1.2. Benchmarking CellOracle & Scenic

CellOracle (v 0.7.5) was obtained from GitHub (https://github.com/ morris-lab/CellOracle commit: cda023a) and installed into a new Anaconda environment. pySCENIC (v0.11.2) was obtained from the python package manager pypi and installed into a new Anaconda environment. A benchmarking module was written for the Inferelator to run CellOracle and pySCENIC from the inferelator workflow. Data loading, crossvalidation, simulation, and scoring functions are identical between all methods. CellOracle was provided the prior knowledge network as a binary dataframe. pySCENIC was provided the prior knowledge network as a ranked-interaction feather database and TF lookup table, in accordance with the pySCENIC pipeline for generating prior knowledge databases for new organisms. Expression data for pySCENIC was log pseudocount transformed and scaled. Expression data for CellOracle was provided as raw counts, which was then log pseudocount transformed and scaled during CellOracle run.

### 1.3. Inferelator 3.0 Single-Cell Computational Speed Profiling

144,682 mouse cells from the mouse neuronal subcluster EXC IT 1 were used with the mouse excitatory neuron prior knowledge network to determine Inferelator 3.0 runtime. To benchmark the python-based multiprocessing engine, the Inferelator was deployed to a single 28-core (Intel*O*R Xeon*O*R E5-2690) node. The Dask implementations of the Inferelator and pySCENIC were deployed to 5 28-core (Intel*O*R Xeon*O*R E5-2690) nodes for a total of 140 cpu cores. Either all 144,682 mouse cells were used, or a subset was randomly selected for each run, and used to learn a single GRN. Runtime was determined by the length of workflow execution, which includes loading data, running all regressions, and producing output files. We were unable to run the full 144k cell data set with pySCENIC due to runtime limitations (with GENIE3) or cryptic memory-related errors (with GRNBOOST2).

### 1.4. Preprocessing Mus musculus single-cell data

Single-cell expression data from *Mus musculus* brain samples taken at E18 was obtained from 10x genomics (10x Genomics, 2017). SCANPY was used to preprocess and cluster the scRNAseq dataset. Genes present in fewer than 2% of cells were removed. Cells were filtered out when fewer than 1000 genes were detected, the cell had more than 20,000 total gene counts, or the cell had more than 7% of gene counts assigned to mitochondrial transcripts. Transcript counts were then log transformed and normalized and scaled. Cells were assigned to mitotic or post mitotic phase based on cell cycle marker genes using score genes cell cycle (Satija *et al*., 2015). In order to focus on neuronal cells, all 374,369 mitotic cells were removed. Remaining cells were clustered by Leiden clustering (Resolution = 0.5) using the first 300 principal components of the 2000 most highly variable genes. Broad cell types were assigned to each cluster based on the expression of marker genes Neurod6 for Excitatory neurons, Gad1 for Interneurons, and Apoe for glial cells. Cells from each broad cell type were then re-clustered into clusters based on the 2000 most highly variable genes within the cluster. Specific cell types were assigned to each subcluster based on the expression of marker genes(Di Bella *et al*., 2020). Ambiguous clusters were discarded, removing 151,765 cells, leaving resulting in 36 specific cell type clusters that consist of 766,402 total cells.

Single-cell ATAC data from *Mus musculus* brain samples taken at E18 was obtained from 10x genomics; datasets are from samples prepared fresh (10x Genomics, 2019c), samples dissociated and cryopreserved (10x Genomics, 2019a), and samples flash-frozen (10x Genomics, 2019b). ChromA (Gabitto *et al*., 2020) and SnapATAC (Fang *et al*., 2021) were used to process the scATACseq datasets. Consensus peaks were called on the 3 datasets using ChromA. Each dataset was then run through the SnapATAC pipeline using the consensus peaks. Cells were clustered and labels from the scRNAseq object were transferred to the scATAC data. Cells that did not have an assignment score *≥ .*5 were discarded. Assigned barcodes were split by cell class( EXC, IN or GL). ChromA was run again for each cell class generating 3 sets of cell class specific peaks.

Aggregated chromatin accessibility profiles were used with TRANSFAC v2020.1 motifs and the inferelator-prior (v0.3.0) pipeline to create prior knowledge connectivity matrices between TFs and target genes for excitatory neurons, interneurons, and glial cells. Vascular cells were not present in the scATAC data sufficiently to allow construction of a vascular cell prior with this method, and so vascular cells were assigned the glial prior for network inference.

### 1.5. Saccharomyces cerevisiae prior knowledge networks

A prior knowledge matrix consists of a signed or unsigned connectivity matrix between regulatory transcription factors (TFs) and target genes.

This matrix can be obtained experimentally or by mining regulatory databases. For a TF gene relationships to be directly causal, the TF must localize to the gene, and gene expression must change in response to perturbations in the TF. However, these criteria do not have to be met at all times. It is reasonable to expect that in many (or most) cell states, a TF may not localize to a target gene, or expression of the gene may not be affected by perturbations in the TF.

Prior knowledge and gold standard networks are selected with these criteria in mind. The YEASTRACT prior knowledge network was obtained from the YEASTRACT database (Teixeira *et al*., 2018; Monteiro *et al*., 2020) (http://www.yeastract.com/; Downloaded 07/13/2019) which is constructed from published yeast TF localization and gene expression data. This prior knowledge network has 11,486 TF gene edges from the YEASTRACT database for which evidence exists that the TF localizes to the target gene, and that the target gene expression changes upon TF perturbation. The yeast gold standard network was constructed in an earlier work (Tchourine *et al*., 2018) and consists of 1,403 edges, which have multiple pieces of both DNA localization and target gene perturbation evidence.

### 1.6. TF Motif-Based Connectivity Matrix (inferelator-prior)

Scanning genomic sequence near promoter regions for TF motifs allows for the construction of motif-derived priors which can be further constrained experimentally by incorporating information about chromatin accessibility (Miraldi *et al*., 2019). We have further refined the generation of prior knowledge matrices with the python inferelator-prior package, which takes as input a gene annotation GTF file, a genomic FASTA file, and a TF motif file, and generates an unsigned connectivity matrix. It has dependencies on the common scientific computing packages NumPy (Harris *et al*., 2020), SciPy (Virtanen *et al*., 2020), and scikit-learn (Pedregosa *et al*., 2011). In addition, it uses the BEDTools kit (Quinlan and Hall, 2010) and associated python interface pybedtools (Dale *et al*., 2011). The inferelator-prior package (v0.3.0 was used to generate the networks in this manuscript) is available on github (https://github.com/flatironinstitute/inferelator-prior) and can be installed through the python package manager pip.

#### 1.6.1. Motif Databases

DNA binding motifs were obtained from published databases. CISBP (Lambert *et al*., 2019) motifs were obtained from CIS-BP (http://cisbp. ccbr.utoronto.ca/; Build 2.00; Downloaded 11/25/2020) and processed into a MEME-format file with the PWMtoMEME module of inferelator-prior. JASPAR (Fornes *et al*., 2020) motifs were obtained as MEME files from JASPAR (http://jaspar.genereg.net/; 8th Release; Downloaded 11/25/2020) . TRANSFAC (Matys *et al*., 2006) motifs were licensed from geneXplain (http://genexplain.com/transfac/; Version 2020.1; Downloaded 09/13/2020) and processed into a MEME-format file with the inferelator-prior motif parsing tools.

#### 1.6.2. Motif Scanning

Genomic regions of interest are identified by locating annotated Transcription Start Sites (TSS) and opening a window that is appropriate for the organism. For microbial species with a compact genome (e.g. yeast), regions of interest are defined as 1000bp upstream and 100bp downstream of the TSS. For complex eukaryotes with large intergenic regions (e.g. mammals), regions of interest are defined as 50000bp upstream and 2500bp downstream of the TSS. This is further constrained by intersecting the genomic regions of interest with a user-provided BED file, which can be derived from a chromatin accessibility experiment (ATAC-seq) or any other method of identifying chromatin of interest. Within these regions of interest, motif locations are identified using the Find Original Motif Occurrences (FIMO) (Grant *et al*., 2011) tool from the MEME suite (Bailey *et al*., 2009), called in parallel on motif chunks to speed up processing. Each motif hit identified by FIMO is then scored for information content (IC) (Kim *et al*., 2003). IC*_i_*, ranging between 0 and 2 bits, is calculated for each base *i* in the binding site, where *p_b,i_* is the probability of the base *b* at position *i* of the motif and *p_b,bg_* is the background probability of base *b* in the genome (Equation 1). Effective information content (EIC) (Equation 2) is the sum of all motif at position *i* is IC*_i_* penalized with the *£*_2_-norm of the hit IC*_i_* and the consensus motif base at position *i*, IC*_i,_*_consensus_.

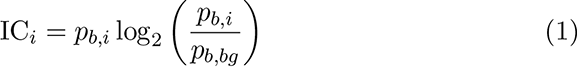

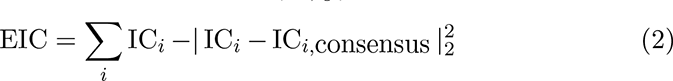

#### 1.6.3. Connectivity Matrix

A TF-gene binding score is calculated separately for each TF and gene. Each motif hit for a TF within the region of interest around the gene is identified. Overlapping motif hits are resolved by taking the maximum IC for each overlapping base, penalized with the *£*_2_-norm of differences from the motif consensus sequence. To account for cooperative TF binding effects, any motif hits within 100 bases (25 bases for yeast) are combined, and their EIC scores are summed. The TF-gene binding score is the maximum TF EIC after accounting for overlapping and adjacent TF motifs, and all TF-gene scores are assembled into a Genes x TFs score matrix.

This unfiltered TF-gene score matrix is not sparse as motifs for many TFs are expected to occur often by chance, and TF-gene scores for each TF are not comparable to scores for other TFs as motif position-weight matrices have differing information content. Scores for each TF are clustered using the density-based k-nearest neighbors algorithm DBSCAN (Ester *et al*., 1996) (MinPts = 0.001 * number of genes, eps = 1). The cluster of TF-gene edges with the highest score values, and any high-score outliers, are retained in the connectivity matrix, and other TF-gene edges are discarded.

#### 1.6.4. CellOracle Connectivity Matrix

CellOracle (Kamimoto *et al*., 2020) was cloned from github (v0.6.5; https://github.com/morris-lab/CellOracle; a0da790). CellOracle was provided a BED file with promoter locations for each gene (200bp upstream of transcription start site to 50bp downstream of transcription start site) and the appropriate MEME file for each motif database. Connectivity matrices were predicted using a false positive rate of 0.02 and a motif score threshold of 6. The inferelator-prior pipeline was run using the same promoter locations and MEME files so that the resulting networks are directly comparable, and the Jaccard index between each network and the YEASTRACT network was calculated. Each motif-based network was used as a prior for inferelator network inference on *Saccharomyces cerevisiae*, with the same 2577 genome-wide expression microarray measurements (Tchourine *et al*., 2018). 20% of the genes were held out of the prior networks and used for scoring the resulting network inference. The motif-based network files have been included in Supplemental Data 1.

### 1.7. Network Inference (The Inferelator)

The Inferelator modeling of gene regulatory networks relies on three main modeling assumptions. First, because many transcription factors (TFs) are post transcriptionally controlled and their expression level may not reflect their underlying biological activity, we assume that the activity of a TF can be estimated using expression levels of known targets from prior interactions data (Arrieta-Ortiz *et al*., 2015; Fu *et al*., 2011). Second, we assume that gene expression can be modeled as a weighted sum of the activities of TFs (Bonneau *et al*., 2006; Castro *et al*., 2019). Finally, we assume that each gene is regulated by a small subset of TFs and regularize the linear model to enforce sparsity.

The Inferelator was initially developed and distributed as an R package (Bonneau *et al*., 2006; Greenfield *et al*., 2010; Madar *et al*., 2010; Greenfield *et al*., 2013). We have rewritten it as a python package with dependencies on the common scientific computing packages NumPy (Harris *et al*., 2020), SciPy (Virtanen *et al*., 2020), pandas (Wes McKinney, 2010), AnnData (Wolf *et al*., 2018), and scikit-learn (Pedregosa *et al*., 2011). Scaling is implemented either locally through python or as a distributed computation with the Dask (Rocklin, 2015) parallelization library. The inferelator package (v0.5.6 was used to generate the networks in this manuscript) is available on github (https://github.com/flatironinstitute/inferelator) and can be installed through the python package manager pip. The Inferelator takes as input gene expression data and prior information on network structure, and outputs ranked regulatory hypotheses of the relative strength and direction of each interaction with an associated confidence score.

### 1.8. Transcription Factor Activity

The expression level of a TF is often not suitable to describe its activity (Schacht *et al*., 2014). Transcription factor activity (TFA) is an estimate of the latent activity of a TF that is inducing or repressing transcription of its targets in a sample. A gene expression dataset (**X**) is a Samples x Genes matrix where *X_i,j_* is the observed mRNA expression level (*i ∈* Samples and *j ∈* Genes), measured either by microarray, RNA-seq, or single cell RNA sequencing (scRNA-seq).

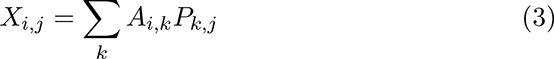

We estimate TFA by solving (Equation 3) for activity (*A_i,k_* ), where *k ∈* TFs, and **P** is a TFs x Genes prior connectivity matrix. *P_k,j_* is non-zero if gene *j* is regulated by TF *k* and 0 if it is not. In matrix notation, **X** = **AP**, and **A^^^** is estimated by minimizing *I* **A^^^ P** *−* **X** *I*^2^. This is calculated by the pseudoinverse **P***^†^* and solving **A^^^** = **XP***^†^*. The resulting **A^^^** is a Samples x TF activities matrix where *A*^^^*_i,k_* is the estimated latent TFA for sample *i* and

TF *k*. In cases where all values in **P** for a TF are 0, that TF is removed from **P** and the expression **X** of that TF is used in place of activity.

### 1.9. Inferelator Network Inference

Linear models (Equation 4) are separately constructed for each gene *j*.

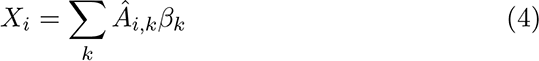

In addition to the model selection methods described here, we have implemented a module which takes any scikit-learn regression object (for example, elastic net (Zou and Hastie, 2005)). Model selection and regularization techniques are applied to enforce the biological property of sparsity. If the coefficient *β_j,k_* is non-zero, it is evidence for a regulatory relationship between TF *k* and gene *j*.

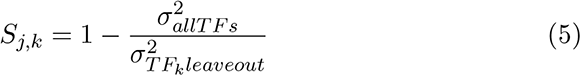

For each gene *j*, the amount of variance explained by each regulatory TF *k* is calculated as the ratio between the variance of the residuals in the full model and the variance of the residuals when the linear model is refit by ordinary least squares (OLS) and *k* is left out (Equation 5).

In order to mitigate the effect of outliers and sampling error, model selection is repeated multiple times using input expression data **X** that has been bootstrapped (resampled with replacement). Predicted TF-gene interactions are ranked for each bootstrap by amount of variance explained and then rank-combined into a unified network prediction. Confidence scores are assigned based on the combined rank for each interaction, and the overall network is compared to a gold standard and performance is evaluated by area under the precision-recall curve.

The effects of setting hyperparameters can be tested by cross-validation on the prior and gold standard networks. This strategy holds out a subset of genes (rows) from the prior knowledge network **P**. Network inference performance is then evaluated on only those held-out genes, using the gold standard network.

#### 1.9.1. Model Selection: Bayesian Best Subset Regression

Bayesian Best Subset Regression (BBSR) is a model selection method described in detail in (Greenfield *et al*., 2013). Initial feature selection for this method is necessary as best subset regression on all possible combinations of hundreds of TF features is computationally intractible. We therefore select ten TF features with the highest context likelihood of relatedness between expression of each gene and activity of each TF. This method is described in detail in (Madar *et al*., 2010).

First, gene expression and TF activity are discretized into equal-width bins (n=10) and mutual information is calculated based on their discrete probability distributions (Equation 6) to create a mutual information matrix

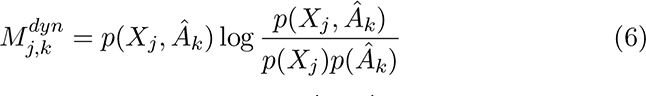

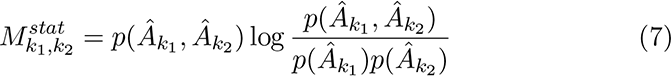

Mutual information is also calculated between activity of each TF (Equation 7) to create a mutual information matrix **M^stat^**.

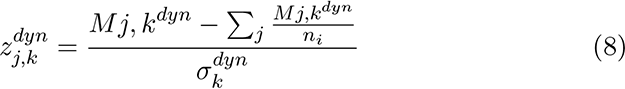

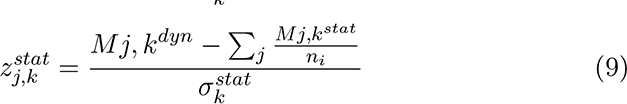

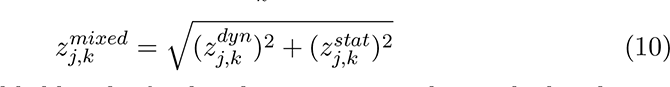

A mixed context likelihood of relatedness score is then calculated as a pseudo-zscore by calculating **Z^dyn^** (Equation 8) and **Z^stat^** (Equation 9). Any values less than 0 in **Z^dyn^** or **Z^stat^** are set to 0, and then they are combined into a mixed context likelihood of relatedness matrix **Z^mixed^** (Equation 10). For each gene *j*, the 10 TFs with the highest mixed context likelihood of relatedness values are selected for regression.

For best subset regression, a linear model is fit with OLS for every combination of the selected predictor variables.

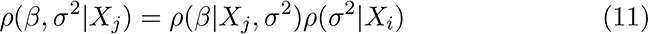

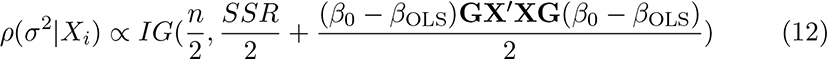

We define *β*_0_ as our null prior for the model parameters (zeros), *β*_OLS_ as the model coefficients from OLS, *SSR* as the sum of squared residuals, and **G** as a *g*-prior diagonal matrix where the diagonal values represent a weight for each predictor variable. *g*-prior weights in **G** close to 0 favor *β* values close to *β*_0_. Large *g*-prior weights favor *β* values close to *β*_OLS_. By default, we select *g*-prior weights of 1 for all predictor variables. From the joint posterior distribution (Equation 11) we can calculate the marginal posterior distribution of *σ*^2^ (Equation 12), where IG is the inverse gamma distribution.

The Bayesian information criterion (BIC) is calculated for each model, where *n* is the number of observations and *k* is the number of predictors (Equation 13).

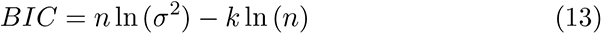

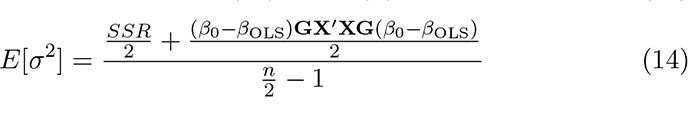

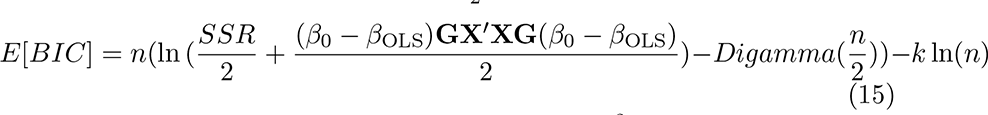

We calculate the expected posterior distribution of *σ*^2^ (Equation 14) for each subset of predictors, and use it to determine the model BIC (Equation 15). We then select the model with the smallest *E*[*BIC*]. The predictors in the selected subset model for gene *j* are TFs which regulate its expression.

#### 1.9.2. Model Selection: StARS-LASSO

Least absolute shrinkage and selection operator (LASSO) (Zou, 2006) combined with the Stability Approach to Regularization Selection (StARS) (Liu *et al*., 2010) is a model selection method described in detail in (Miraldi *et al*., 2019). In short, the StARS-LASSO approach is to select the optimal *λ* parameter for (Equation 16). *N* random subsamples of *X* and *A*^^^ without replacement subnetworks *S_n,λ_* are defined as the non-zero coefficients *β_n,λ_* after LASSO regression. Initially, *λ* is set large, so that each subnetwork *S_n_* is highly sparse, and is then decreased, resulting in increasingly dense networks. Edge instability is calculated as the fraction of times subnetworks disagree about the presence of an network edge. As *λ* decreases, the subnetworks are expected to have increasing edge instability initially and then decreasing edge instability as *λ* approaches 0, as (Equation 16) reduces to OLS and each subnetwork becomes dense.

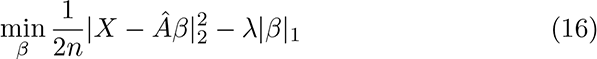

We choose the largest value of *λ* such that the edge instability is less than 0.05, which is interpretable as all subnetworks share *>* 95% of edges. This selection represents a balance between increasing the network size and minimizing the instability that occurs when data is sampled.

### 1.10. Multiple Task Network Inference

We separate biological samples which represent different states into separate tasks, learn networks from these tasks, and then combine task-specific networks into an ensemble network. One method of solving these states is to sequentially apply a single-task method for network inference (i.e. 1.9.1 or 1.9.2). The networks generated for each task are then rank-combined into a unified network. The Adaptive Multiple Sparse Regression (AMuSR) method, described in detail in (Castro *et al*., 2019), uses a multi-task learning framework, where each task is solved together.

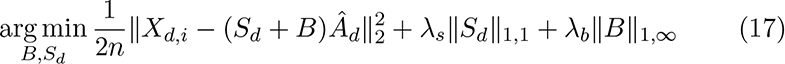

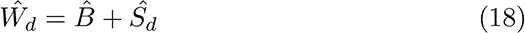

In (Equation 17), *B* is a block-sparse weight matrix in which the weights for any feature are the same across all tasks. *S_d_* is a sparse weight matrix for task *d*, allowing weights for features to vary between tasks. The combination *W_d_* of *B* and *S_d_* (Equation 18) are model weights representing regulatory interactions between TFs and genes for task *d*. In short, this method uses adaptive penalties to favor regulatory interactions shared across multiple tasks in *B*, while recognizing dataset specific interactions in *S_d_*. Model hyperparameters *λ_s_* and *λ_b_* are identified by grid search, selecting the model that minimizes the extended Bayesian Information Criterion (eBIC) (Equation 19), where *D* is the number of task datasets, and for dataset *d*, *n_d_* is the number of observations, *X*^(^*^d^*^)^ is gene expression for gene *i*, *A*^^(^*^d^*^)^ is TF activity estimates, *W_∗,d_* is model weights, *k_d_* is the number of non-zero predictors, and *p_d_* is the total number of predictors. For this work, we choose to set the eBIC paramater *γ* to 1.

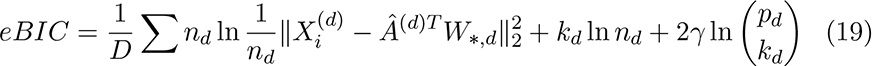

### 1.11. Network Performance Metrics

Prior work has used the area under the Precision (Equation 20) Recall (Equation 21) curve to determine performance, by comparing to some known, gold-standard network. Here we add two metrics; Matthews correlation coefficient (Matthews, 1975) (MCC) (Equation 22) and F1 score (Equation 23). MCC can be calculated directly from the confusion matrix

True Positive (TP), False Positive (FP), True Negative (TN), and False Negative (FN) values.

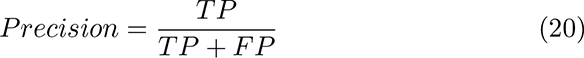

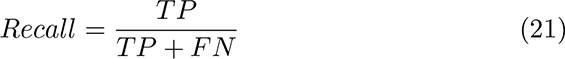

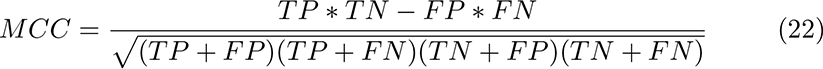

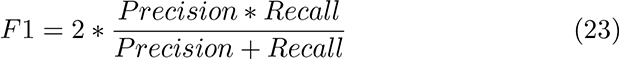

*Precision* + *Recall*

We compute an MCC and F1 score for each cutoff along ranked interactions in order to generate MCC and F1 scores for all possible networks in growing ranked order. The maximum MCC along ranked interactions gives the subnetwork that has maximum similarity to the comparison network, accounting for TP, FP, TN, and FN. The maximum F1 along ranked interactions gives the subnetwork that has maximum similarity to the comparison network accounting for TP, FP, and FN.

### 1.12. Visualization

Figures were generated with R (R Core Team, 2020) and the common ggplot2 (Wickham, 2016), umap (McInnes *et al*., 2018), and tidyverse packages (Wickham *et al*., 2019). Additional figures were generated with python using scanpy (Wolf *et al*., 2018), matplotlib (Hunter, 2007), and seaborn (Waskom, 2021). Network diagrams were created with the python package jp gene viz (Watters, 2019). Schematic figures were created in Adobe Illustrator, and other figures were adjusted in Illustrator to improve panelling and layout.

## Availability of Data and Materials

The datasets supporting the conclusions of this article are available in the NCBI GEO repository with accession IDs: GSE125162, GSE144820, GSE67023, GSE27219, GSE142864. A large number of GEO records were compiled and normalized in a previous work Tchourine *et al*. (2018) into a combined dataset which is available on Zenodo (DOI: 10.5281/zenodo.3247754). The scRNAseq expression matrix, metadata, prior knowledge network, and gold standard network for the yeast network inference benchmarking is available on Zenodo (DOI: 10.5281/zenodo.5272314). Single-cell mouse datasets are publicly available from 10x genomics 10x Genomics (2017, 2019c,a,b) under a Creative Commons Attribution (CC-BY 4.0) license. Software packages developed for this article are available on github (https://github. com/flatironinstitute/inferelator and https://github.com/flatironinstitute/ inferelator-prior) and have been released as python packages through PyPi (https://pypi.org/project/inferelator/ and https://pypi.org/ project/inferelator-prior/). Specific analysis scripts for this work have been included in Supplemental Data 1.

## Author’s contributions

CSG contributed to Methodology, Software, Validation, Formal Analysis, Writing – Original Draft Preparation, and Visualization. CJ and GS contributed to Conceptualization, Methodology, Software, Validation, Investigation, Resources, Data Curation, Formal Analysis, Writing – Original Draft Preparation, and Visualization. AS contributed to Validation, Data Curation, Formal Analysis, and Visualization. AW contributed to Software and Visualization. AT contributed to Software, Writing – Original Draft Preparation, and Formal Analysis. DC and KT contributed to Software, Data Curation, and Conceptualization. NDV, NC, RY, and TH contributed to Software. DG contributed to Supervision, Project Administration, and Funding Acquisition. EM contributed to Conceptualization, Writing – Original Draft Preparation, and Software. RB contributed to Conceptualization, Writing – Original Draft Preparation, Supervision, Project Administration, and Funding Acquisition.

## Additional Files

- Supplemental Data 1 is a .tar.gz file containing the prior knowledge networks used in this work, the gold standard networks used in this work, and the python scripts used to generate the learned networks in this work
- Supplemental Data 2 is a .tar.gz file containing the mouse E18 neuronal network learned in Figure 6 of this work
- Supplemental Table 1 is a .tsv file containing the crossvalidation performance results from Figure 2
- Supplemental Table 2 is a .tsv file containing the crossvalidation performance results from Figure 3
- Supplemental Table 3 is a .tsv file containing the crossvalidation performance results from Figure 4B-D
- Supplemental Table 4 is a .tsv file containing the crossvalidation performance results from Figure 4G
- Supplemental Table 5 is a .tsv file containing the crossvalidation performance results from Supplemental Figure 5A
- Supplemental Table 6 is a .tsv file containing the crossvalidation performance results from Figure 4H
- Supplemental Table 7 is a .tsv file containing the crossvalidation performance results from Supplemental Figure 3

---

**Supplemental Figure 1:**
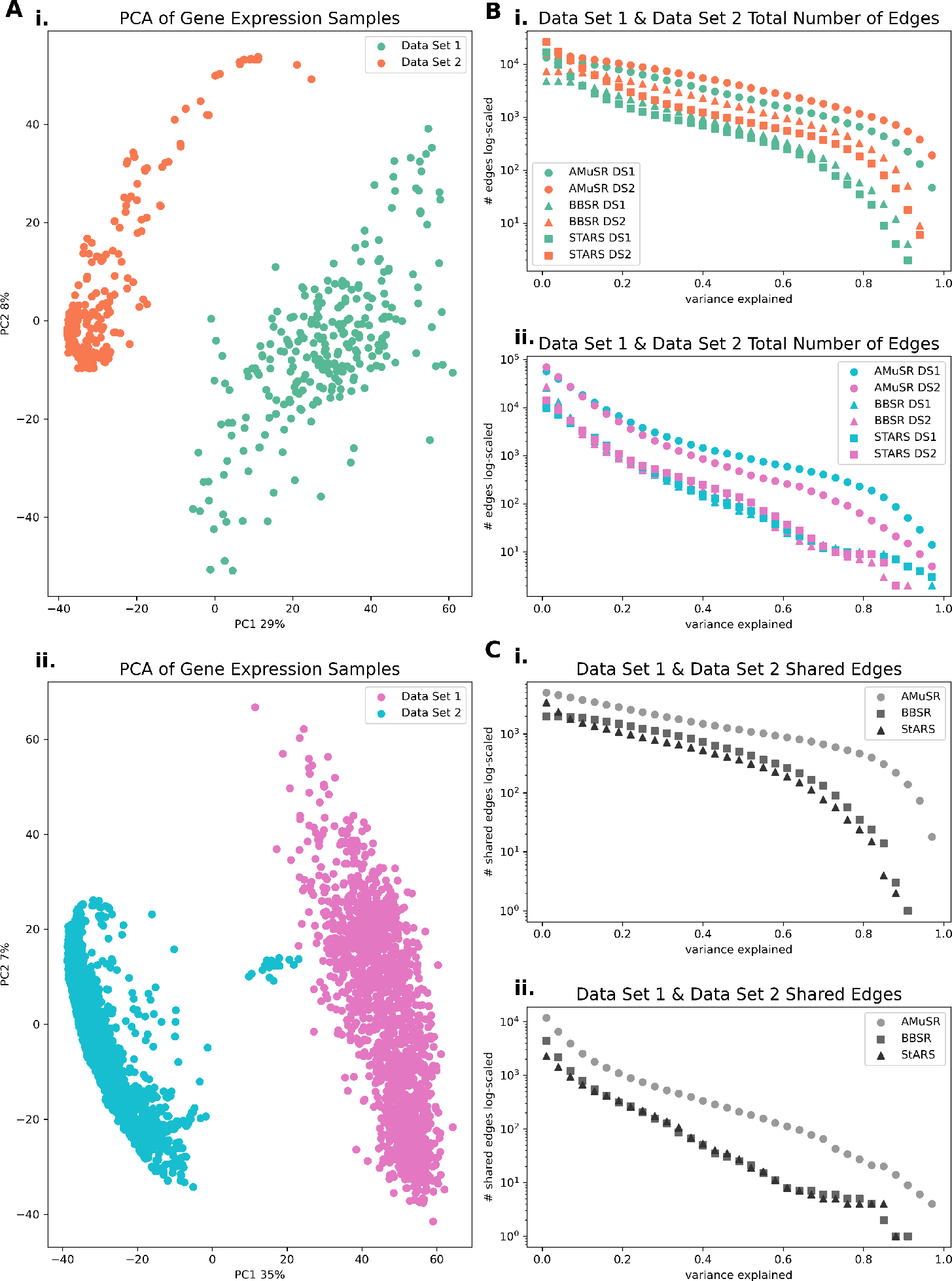
Learning *Bacillus subtilis* and *Saccharomyces cerevisiae* networks by tasks. (**A**) PCA depicts batch effects between datasets for both (i) *Bacillus subtilis* and (i) *Saccharomyces cerevisiae*. Learning networks by treating the independently collected datasets as separate tasks allows for sharing regulatory commonalities while respecting experimental variance. (**B**) The number of shared edges between the two datasets, for both model organisms (i) and (ii), shows a high number of overlapping edges. Edges are ranked by their corresponding variance explained for each of the three different model selection approaches: AMuSR, BBSR, and StARS-LASSO. (**C**) Across the three different model selection approaches, AMuSR learns the highest number of overlapping edges between the respective datasets for model organisms (**i**) and (**ii**).

**Supplemental Figure 2:**
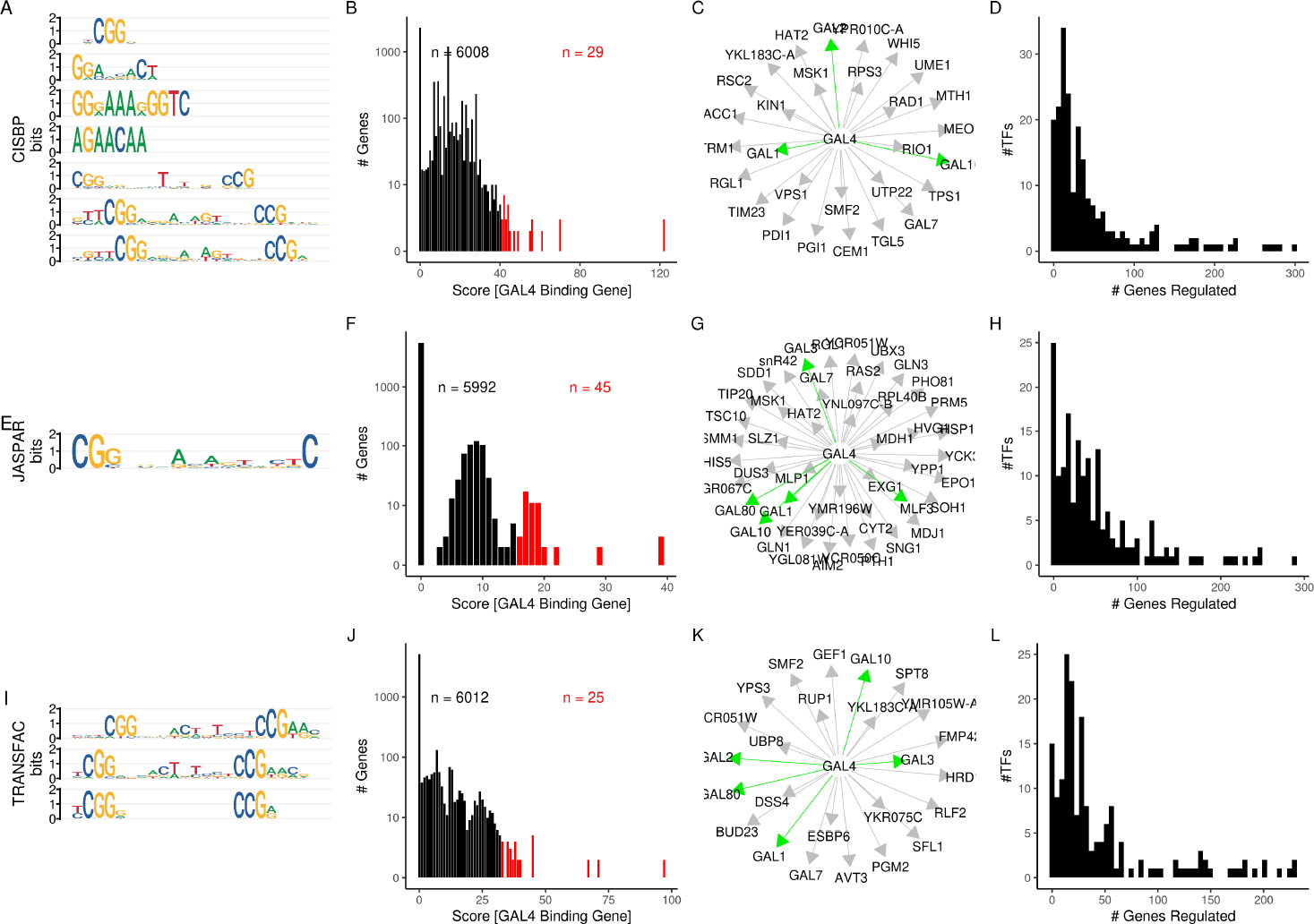
Network construction using TF motifs in Saccharomyces cerevisiae. (**A**) Motifs annotated for GAL4 in the CIS-BP motif database. (**B**) Histogram of scores linking GAL4 to target genes. Genes in black have been omitted from the final connectivity matrix, and genes in red have been included. (**C**) Network connecting GAL4 and target genes. Green edges are present in the YEASTRACT database. (**D**) Histogram of out degree for each TF in the complete network. (**E-H**) Network analysis as A-D for the JASPAR motif database. (**I-L**) Network analysis as A-D for the TRANSFAC PRO motif database.

**Supplemental Figure 3:**
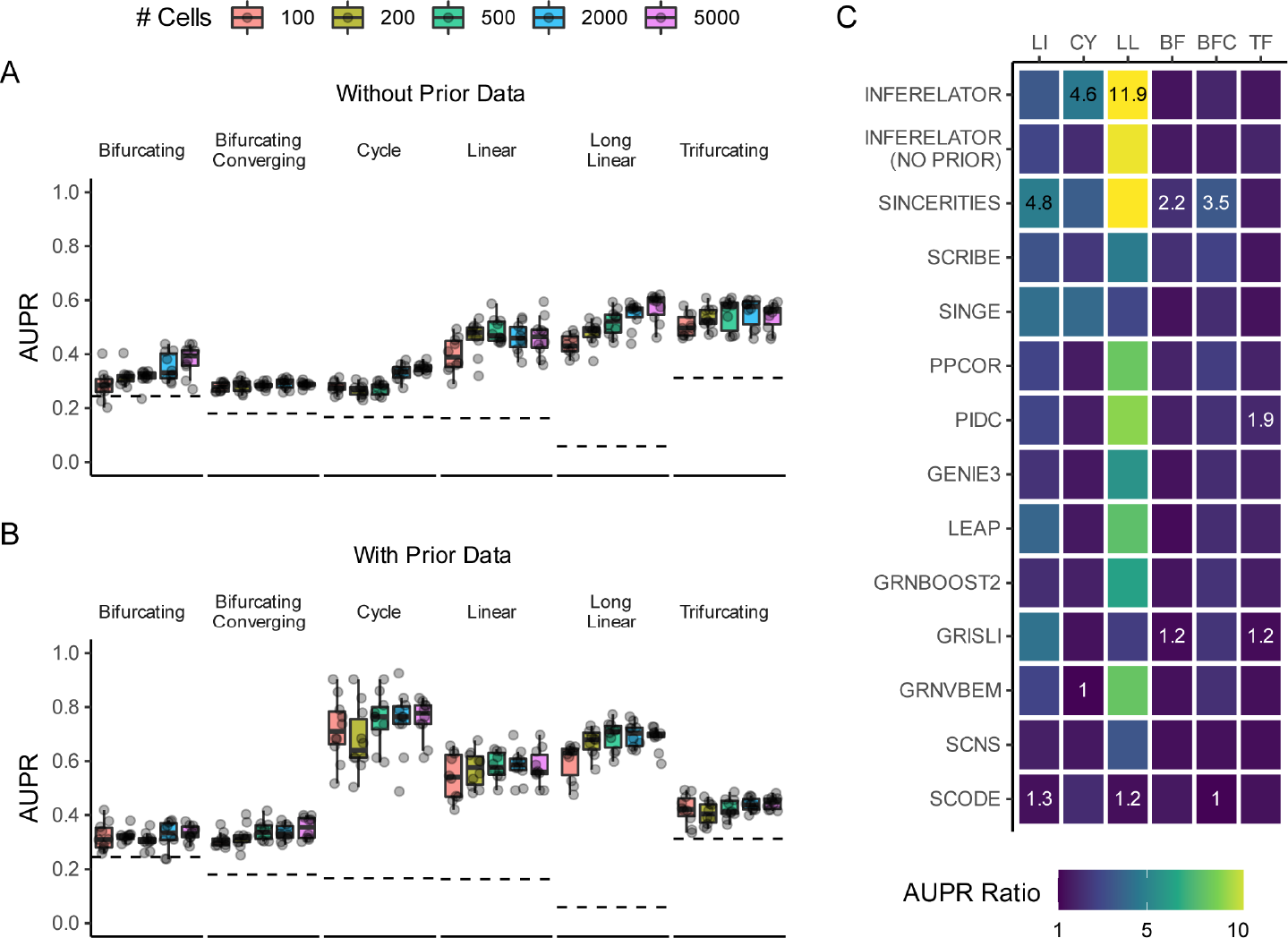
Inferelator performance on BEELINE simulated network data. (A) Network inference performance of the Inferelator with BBSR model selection as measured by AUPR against the ground truth with no prior network information provided. Dashed lines are the expected baseline of a random predictor. (**B**) Network inference performance of the Inferelator with BBSR model selection as measured by AUPR against half of the ground truth with the other half of the ground truth provided as prior network information. Each point is the median performance of 10 differently-seeded splits. (**C**) Comparison of the AUPR ratio over the baseline for the Inferelator to each of the network inference methods used in the original BEELINE benchmark.

**Supplemental Figure 4:**
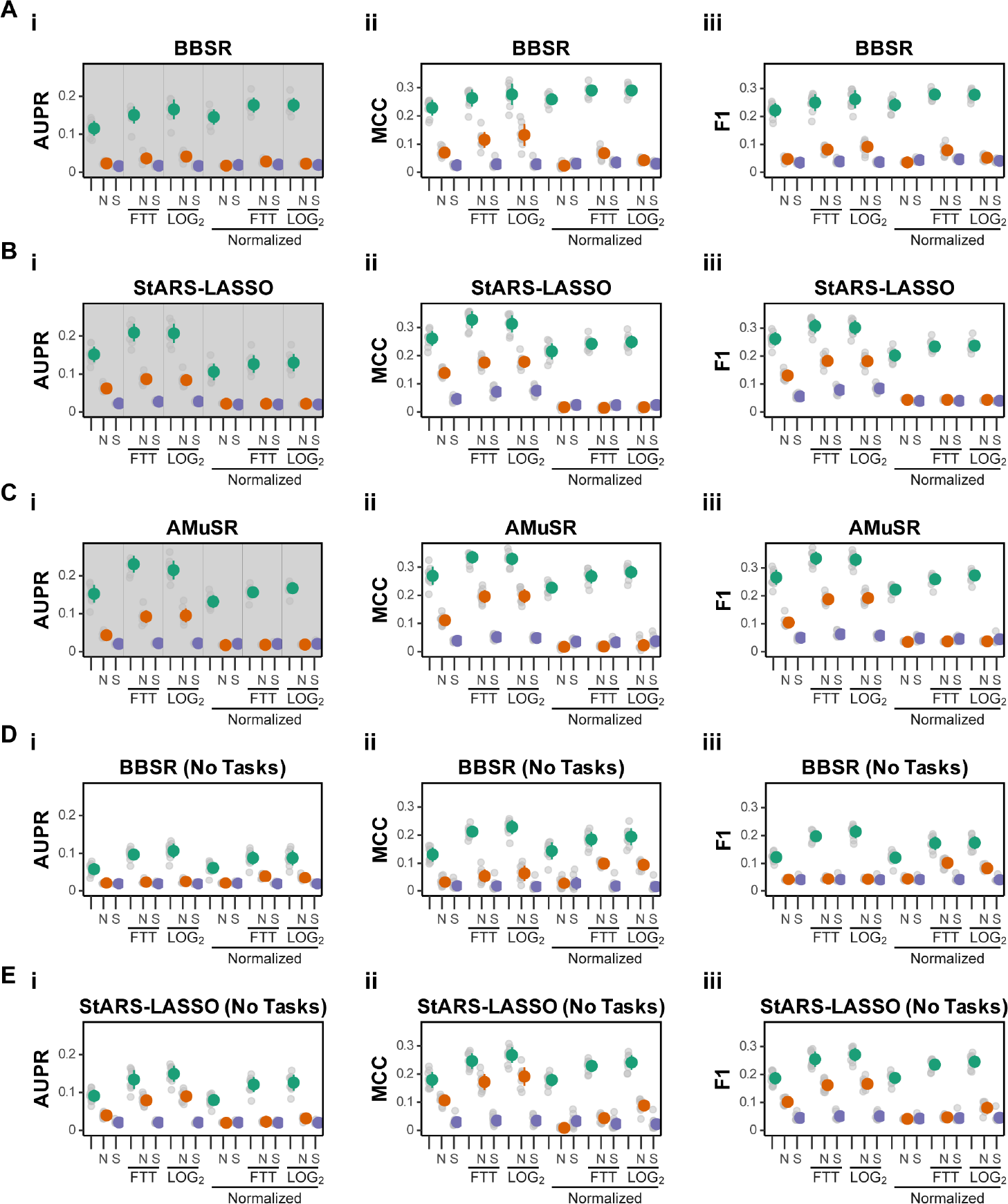
Extended single-cell yeast network performance metrics as measured by (**i**) AUPR, (**ii**) Matthews Correlation Coefficient (MCC), and (**iii**) F1 score. Each gray dot represents performance of one network inference run. Colored dots represent the mean and standard deviation. (**A**) Single-cell yeast network inference performance of BBSR model selection Plots with a gray background are the same plots as used in maintext Figure 4. (**B**) Performance of StARS-LASSO model selection. (**C**) Performance of AMuSR model selection. (**D**) Performance of BBSR model selection where all cells are used without splitting into multiple tasks. (**E**) Performance of StARS-LASSO model selection where all cells are used without splitting into multiple tasks.

**Supplemental Figure 5:**
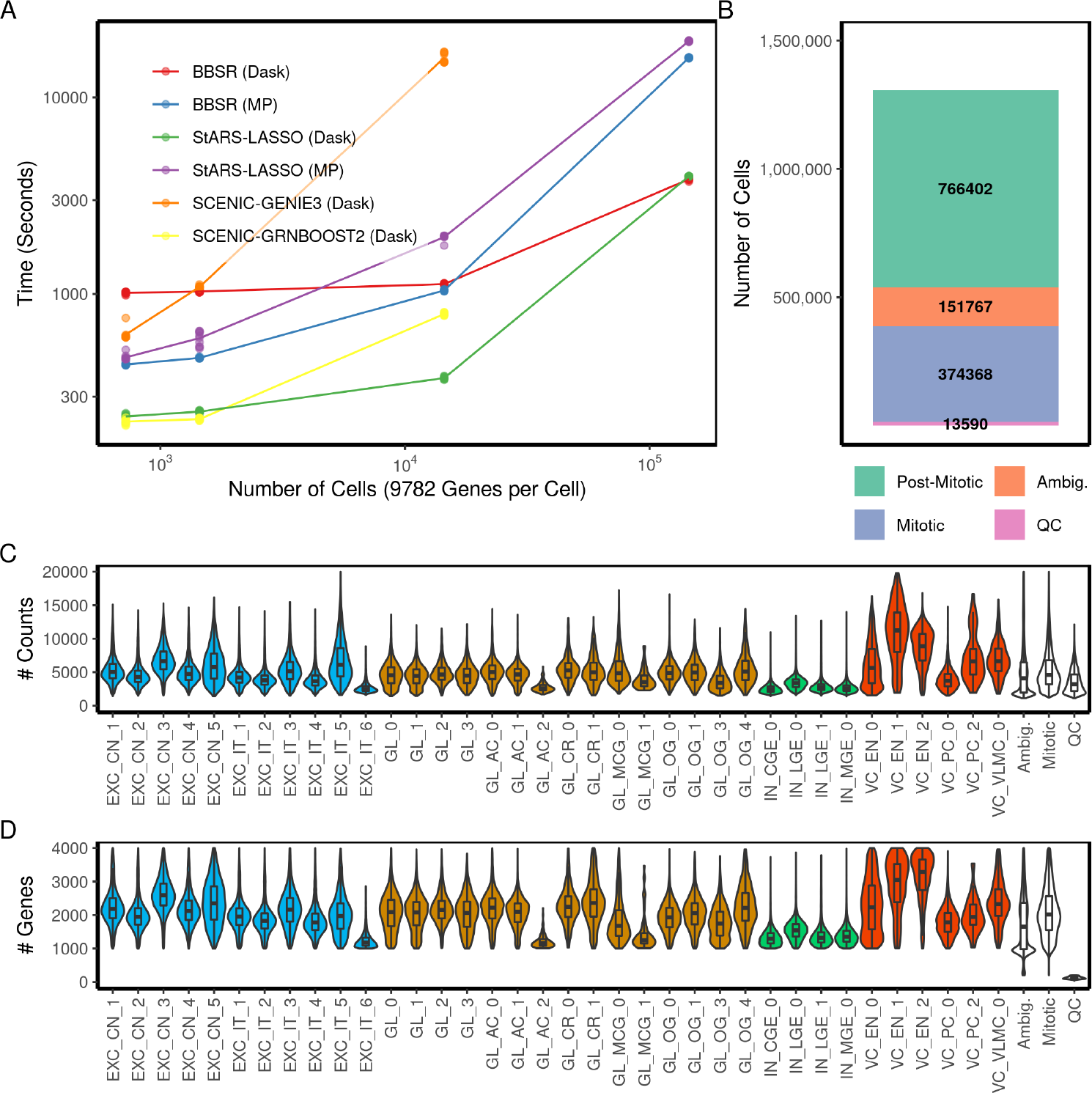
(**A**) Computational performance as measured by runtime in seconds using the Dask engine (140 cpu cores) for the Inferelator 3.0 (BBSR or StARSLASSO), and for SCENIC (GENIE3 or GRNBOOST2). Performance is also measured for the Inferelator 3.0 or using the python-based multprocessing (MP) engine (28 cpu cores). Expression data is sampled from 144,000 mouse cells and 9,782 genes are modeled for network inference. Runtime is shown for 10 replicate runs for each quantity of cells. (**B**) Number of cells removed during preprocessing for Quality Control (QC), as Mitotic, and as Ambiguous by neuronal marker. Post-mitotic, non-ambiguous cells are retained and clustered. (**C**) Number of single-cell counts per cell in each of 36 cell type-specific groups, and in the groups removed during preprocessing. (**D**) Number of genes per cell in each of 36 cell type-specific groups, and in the groups removed during preprocessing

**Supplemental Figure 6:**
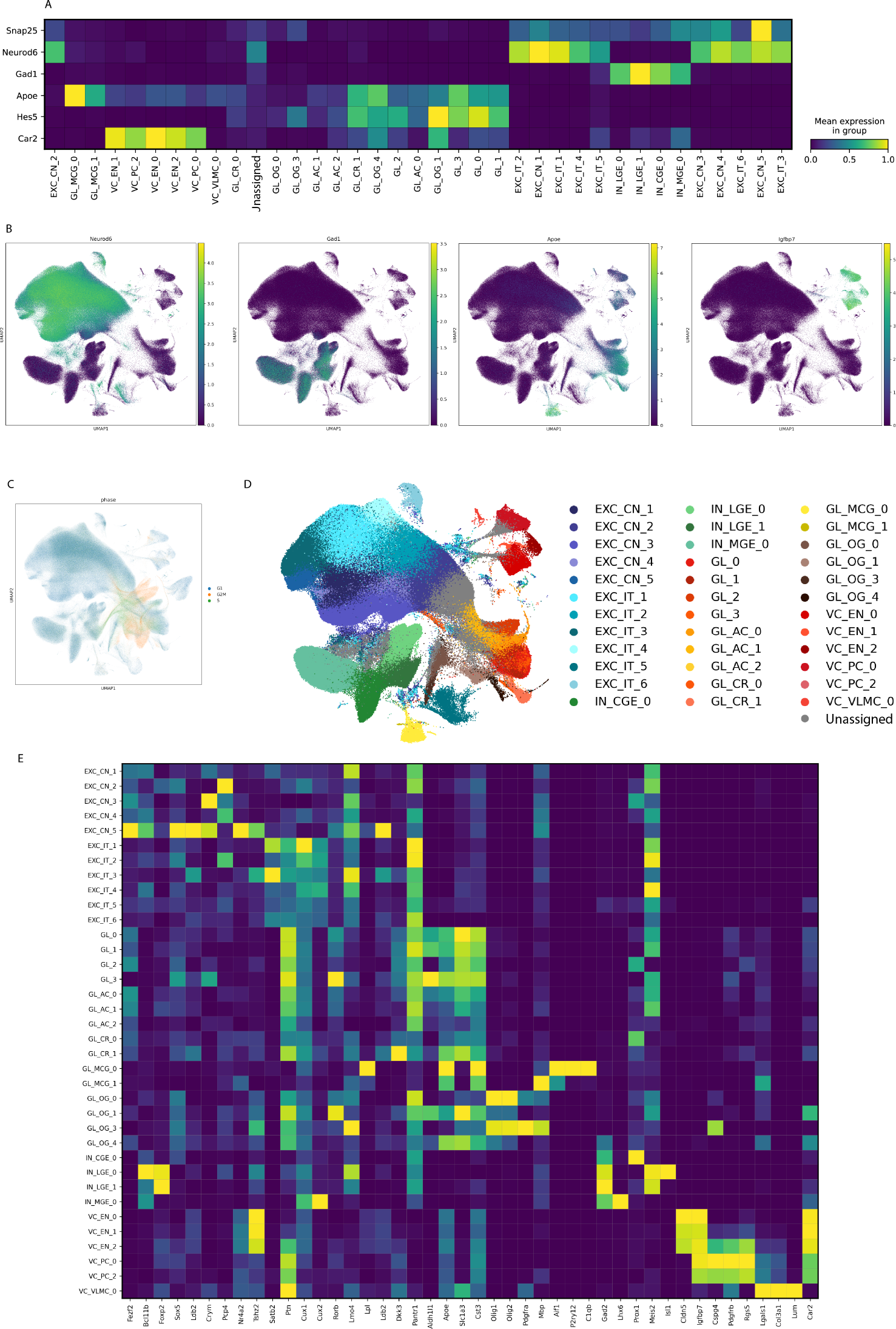
(**A**) Cell class marker expression for each annotated subcluster in mouse single-cell brain data. (**B**) UMAP of 766,402 mouse brain cells colored by cell class marker expression. (**C**) UMAP of 1.3M mouse brain cells colored by the assigned cell cycle phase. (**D**) UMAP of 766,402 mouse brain cells colored by 36 assigned subcluster. (**E**) Cell type marker expression by assigned subcluster.

**Supplemental Figure 7:**
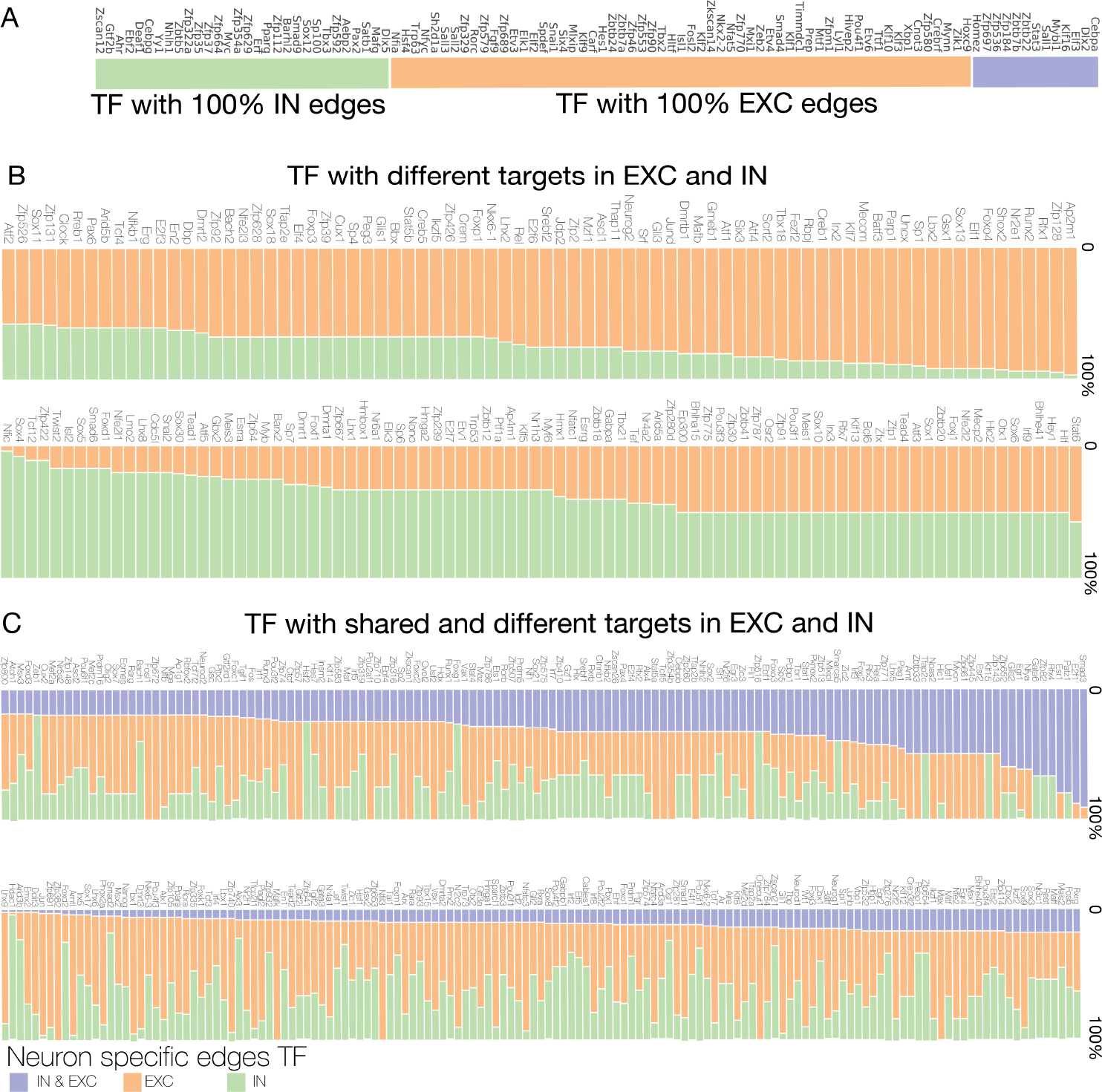
(**A**) List of TFs that have identical target genes in GRNs for both Excitatory neurons (EXC) and Interneurons (IN), that have only target genes in Excitatory neurons, and that have only target genes in Interneurons. (**B**) List of TFs that have no shared target genes in GRNs for Excitatory neurons and in GRNs for interneurons. (**C**) TFs that have some shared target genes in GRNs for Excitatory neurons and interneurons, but also have some target genes specific to Excitatory neurons or interneurons.

## 2. Response to Reviewers

We’d like to thank all of the reviewers for the time that they’ve spent evaluating this manuscript. We believe that the revised manuscript is substantially improved thanks to these comments. To summarize, the most important concern raised by reviewers 1, 3, and 4 is that there is no adequate benchmark against other network inference algorithms. Reviewer 1 has also raised several textual concerns, suggested tests for robustness, and requested clarification on two points related to model design. Reviewer 2 has raised a mathematical argument suggesting that this method is flawed in concept. Reviewer 3 has also raised several specific concerns about the prior and testing networks and the interpretation of inferred networks. Finally, Reviewer 4 has raised several interesting points related to some subtle observations in our model performance.

### 2.1. Summary of Changes

As the most general concern, we address benchmarking first. We initially chose not to include competitive benchmarks against other network inference methods. A neutral benchmarking panel (as recommended by Reviewer 1) is an excellent suggestion and we have included an evaluation of the Inferelator on the BEELINE standard as a new supplemental figure (Supplemental Figure 3). We note that the BEELINE benchmarking is not designed for network inference tools which utilize prior network knowledge during inference (it is a benchmark built around pseudotime). While the Inferelator is adequate to that benchmark, additional benchmarking is necessary.

We have additionally tested two other single-cell network inference tools which utilize prior network knowledge (SCENIC and CellOracle) on the yeast single-cell network inference problem as a benchmark. Yeast is a model organism with real-world single-cell data and which has a reliable gold standard that we can use for performance quantification. We report these results in figure 4, panels G-H. We also report the performance of the GRNBOOST2 network inference method which does not utilize prior data (one component of the SCENIC pipeline) in figure 4H.

In short, the Inferelator is the only method which can learn edges for genes which no prior knowledge is known, and is robust to noise in the prior knowledge network. CellOracle performs very well when given a prior knowledge network and asked to make predictions within that network, although it is more sensitive to noise in the prior knowledge network. We have revised our runtime benchmark in Supplemental Figure 5A to include SCENIC. We have also revised the discussion to include the comparative results and to emphasize the importance of the model organism benchmark we’ve chosen for this work.

In accordance with Reviewer 1’s suggestions, we have revised the introduction to cover prior work and community benchmarks. We have also revised the discussion to better justify the modeling strategy in the context of the results we show. Supplemental Figure 4 now includes performance metrics for the yeast benchmark when networks are learned on all cells together, instead of by task group. We have modified figure 1 to emphasize that we are scoring on information held out of the modeling.

We have predominantly responded to Reviewer 2 in this document, providing specific theoretical and experimental results to contradict the assertion that our modeling strategy is fatally flawed. We have added a prior knowledge network experiment where false positive edges are added prior to modeling in Figure 4H in part to specifically refute the reviewer’s assertions. We have added a section to our methods to answer Reviewer 3’s questions about the selection of our prior knowledge and gold standard networks. Reviewer 4 requested interpretation of several subtle observations in our results. We have modified Figure 4B-D and added runtime benchmarks for SCENIC to Supplemental Figure 5.

We also note that during this revision, we identified a minor error in the construction of the yeast single-cell expression data (several genes were inadvertently dropped when different data sets were merged). We have fixed that error and repeated all analyses that used that data set; no conclusions have changed.

Point by point responses to the reviewer comments follow.

### 2.2. Reviewer 1

Comments to the Author Inferelator 3.0 is a new version of the Inferelator that provides a workflow for five different regression and model selection modules. This version supports single-cell gene expression data and has better scalability, as shown through experiments with the 10x 1.3 million cell mouse neuronal dataset. The authors highlight their method for selecting regulatory edges to retain in a GRN ranking regulatory edges by the amount of target gene variance explained, and selecting a threshold that maximizes MCC against a known gold standard. The Inferelator tool seems to be well-documented and available through PyPi and Github.

Some major comments suggested for revision:

1. Introduction needs a lot of work. Lacks comprehensive discussion of previous work and of many related methods (such as those in this benchmarking paper https://www.nature.com/articles/s41592-019-0690-6) and further explanation of 3 model selection methods used in paper.

*•* We have revised the introduction to give a clearer description of the inferelator, as well as the two most comparable other methods (CellOracle and SCENIC). We note that in the interest of space, we now rely on the excellent work of three benchmarking papers, including the BEELINE benchmarking paper, to describe the many other extant methods for network inference.

2. The paper does no comparison (of performance, time, memory, or other measures) of Inferelator to other existing methods, including SCENIC and others mentioned. Please see benchmarking paper here for ideas on metrics: https://www.nature.com/articles/s41592-019-0690-6

*•* This is an excellent suggestion. We have chosen to apply the inferelator to the simulated BEELINE benchmarks, and report those results in Supplemental Figure 3. Only the BBSR method for model selection was tested, as there are no separable tasks for AMuSR in the BEELINE simulated data, and the overall network size is too small to use a stability-based model selection method like StARS-LASSO. We do note however, that the BEELINE framework was not developed for network inference methods which utilize prior network knowledge (this is why the BEELINE benchmark evaluates the GENIE3 and GRNBOOST2 components of SCENIC without running the full SCENIC pipeline; SCENIC requires prior knowledge).

We have therefore chosen to also benchmark SCENIC and CellOracle on the yeast single-cell network inference problem which has a reliable gold standard. We report those results in Figure 4G-H. In summary, CellOracle has a number of desirable characteristics in a network inference method, and performs well at evaluating a prior network for edges to retain. However, it is not capable of making predictions outside the prior network. The inferelator performance is somewhat lower than CellOracle when scored against a gold standard which was not held out of the prior network, but is capable of making novel predictions outside of the prior network (and therefore performs well when scored against a gold standard held out of the prior network). SCENIC is not capable of making predictions outside of a prior network, and performs poorly when making predictions within a prior network. We have also added a set of runtime benchmarks for SCENIC to Supplemental Figure 5 (CellOracle has not reached a development stage where it would be fair to include in a benchmark for runtime).

1. 3. The paper more or less proposes to port their existing regression methods to single cell data without assessing how peculiarities of single cell data are affected by their approaches. For example, the authors discuss the noise inherent in single cell data, robustness of their regression methods to varying levels of dropout noise (as these can vary from experiment to experiment) can be shown on known ground truth data generated artificially or using benchmarks from the DREAM GRN challenge.

*•* This is largely correct we believe that single-cell data is undersampled, but the increased scale of data collection makes that drawback less critical. We have found Svensson 2020 (https://doi.org/10.1038/s41587-019-0379-5) to be generally correct in all aspects when it comes to interpreting single-cell count data. We note that the most successful methods for single-cell network inference generally do not use models which include single-cell peculiarities (like zero-inflation), but instead rely on models that are robust to noise (CellOracle, for example, uses bagging regression, which is in our opinion an elegant choice to minimize the influence of noise, and that method performs quite well).

We have added several sentences to the results to explain this: Single-cell data is undersampled and noisy, but large numbers of observations are collected in parallel. As network inference is a population-level analysis which must already be robust against noise, we reason that data preprocessing techniques that improve per-cell analyses (like imputation) are unnecessary. We demonstrate that this is valid by quantitatively evaluating networks learned from *Saccharomyces cerevisiae* scRNAseq data with a previously-defined yeast gold standard.”

4. Another interesting experiment is to assess the robustness of networks using subsampling of the single cell data, networks should be robust between subsampling strategies.

*•* This is an excellent suggestion, and the reviewer’s point related to noise is something we have considered at length. We have performed the suggested subsampling experiment in prior work and found that performance increases as a function of cell count up to a point where it plateaus (https://doi.org/10.7554/eLife.51254 Fig 5B). This is consistent with our expectation is that sampling noise in single-cell expression data is manageable via increasing N.

We therefore choose instead to investigate the effect of noise on the prior knowledge network, which is noise that we cannot compensate for experimentally (the effect of noise in the prior was a question raised by Reviewer 4). We have tested the performance of the Inferelator on yeast single-cell network inference when the prior network has random noise added and reported the results in Figure 4H. We find that addition of spurious, false edges to the prior knowledge network does decrease performance, but only modestly, indicating that the Inferelator is robust to noise in the prior knowledge network. A comparison to SCENIC and CellOracle has been provided, in addition to negative controls.

1. 5. Another single-cell specific concern I have is the time lag between TF activity and target expression within a cell. Due to mixing in bulk samples this seems to be less of a concern, but within a single cell sample simultaneous observation of both activities may be sparse.

*•* We are unfortunately unable to directly observe TF activity (direct measurement of activity would be exceptionally useful, and we hope to have that data someday). Instead, we estimate TF activity based on the expression of known gene targets. This estimate is done per-cell and depends on the current cell gene expression, and not the TF expression in the past. We therefore do not expect there to be a ’time lag’ between TF activity and target expression, as we do not currently incorporate time or pseudotime information in our single-cell network modeling. Applying an explicitly dynamic model to network inference is an area we are actively exploring, but represents an entirely different modeling approach and would not be suitable for addition to this work.

1. 6. Finally, what is the justification of doing the inference ”per cell type”, clustering or partitioning data to some arbitrary level using Leiden or Louvain does not necessarily define regulatory program-specific cells. Indeed other approaches such as SCENIC are more local in their learning of regulatory networks. What effect does the resolution of this clustering or the neighborhood have on their inference?

*•* SCENIC does not locally estimate GRNs. SCENIC is explicitly a global method, using prior network knowledge to identify regulatory units in a provisional draft network created from global gene ”adjacencies”. This global GRN is then applied to each cell (with the AUCell function) to determine how well each regulatory unit explains gene expression in that cell as a metric, not as part of the learning process.

We propose (as does CellOracle, which clusters as part of its core workflow) that using a neighborhood-based clustering approach allows us to identify groups of cells which are running different gene regulatory programs. This is of particular value when we are unable to directly observe chromatin state in complex eukaryotes, as TF gene relationships are likely to be dependent on having the ability to access specific enhancer or promoter regions. Treating these cells with different chromatin states as separate learning tasks allows our method to learn common regulatory network components which are active in multiple tasks as well as cluster-specific network components which are active in a limited number of clusters.

To illustrate the value of task-wise learning, we have added performance metrics for network inference on the yeast single-cell data without task separation to Supplemental Figure 4. We see that overall performance is substantially diminished when learning a network on all cells together, without tasks.

Minor comments:

1. The authors state in the introduction ”a major difficulty is that biological systems have large numbers of both regulators and targets; there is poor network identifiability because many plausible networks can explain observed expression data and the regulation of gene expression in an organism” It is unclear if the difficulty is due to the large numbers of regulators and targets (as it was previously stated that only 6% of the human genomes are TFs) or due to redundancy of networks/pathways.

*•* Network size is a difficulty but many large problems exist in machine learning, and so is not insurmountable. Many pathways are redundant or interdependent in ways that simply cannot be deconvoluted computationally (instead requiring careful biological perturbation, which may or may not be possible). We can realistically generate thousands of networks which offer approximately equal explanatory power, and determining which network is correct is an unsolved problem. We have revised the introduction to make this point clearer.

2. The claim in the discussion that ”many of the performance differences between gene regulatory network inference methods are not due to clever methods for model selection, but are instead the result of differences in data cleaning and preprocessing” is a strong one and requires further citation or evidence.

*•* We refer to Figure 4, where preprocessing differences dwarf the differences between model selection methods (despite using three model selection methods which have very different characteristics). This statement is intended to emphasize the importance of using common preprocessing and scoring techniques when comparing network inference methods, as these techniques can introduce or obscure correlations in both predictable and unpredictable ways. We understand this to be commonly accepted wisdom in the statistical learning field (An early warning about data preprocessing from the 19th century is an interesting read: https://doi.org/10.1098/rspl.1896.0076). We have revised the statement to be more specific: ”For example, we find that performance differences between our methods of model selection may be smaller than differences caused by data cleaning and preprocessing.”

3. Please report AUPRC ratio (to the random baseline) instead of AUPRC for better understanding of model performance.

*•* We have reported an AUPRC ratio in addition to AUC for the BEELINE comparison in Supplemental Figure 3. However, we respectfully decline to do so for other analysis in this work. Reporting AUC as a ratio to baseline is a practice that we do not feel is advisable. We can generate several model baselines for example, a model baseline from shuffling labels and a model baseline from replacing data are not identical, and may not be equal to a model baseline calculated based on the gold standard density. It is a best practice to generate multiple baselines to control for different things and report them separately. Furthermore, the interpretation of a model that reports an AUPR of 0.5 over a baseline of 0.05 would differ from a model that reports an AUPR of 0.01 over a baseline of 0.001 and this substantial difference would be lost with ratios.

4. List as a limitation that model is not able to add or learn edges that do not exist in prior networks

*•* This is not a limitation of this modeling strategy. A key advantage of our work is that we are able to add or learn edges, even when there is no information about a gene in the prior. Model performance as reported in figures 2-4 is based on holding genes out of the prior networks entirely and scoring on these genes for which the model has no prior information. We have modified Figure 1 to clarify this.

### 2.3. Reviewer 2

This manuscript discusses an update to Inferelator (version 3.0). This manuscript builds on several other work by the authors (e.g. Inferelator-Amusr) and utilizes these methods that are previously developed as part of the study.

Due to this reliance on previous methods, the issues present in the authors’ previous work (PMID: 30677040, Catro et al 2019) is also inherited in this work and has tainted the results. Consequently, unless theses major issues are addressed, there is not much point in reviewing other aspects of the manuscript. As a result, I focus on detailing these issues and hope that the authors would address and rectify them before moving forward.

The main issue is with the algorithm Inferelator-AMuSr. From the algorithmic side, this method (PMID: 30677040) is quite interesting and utilizes block sparsity and different regularization techniques to learn gene regulatory networks. Unfortunately, the problem formulation is flawed and follows a circular logic. This method uses gene (and TF) expression values across different conditions + a prior network of gene-TF associations (e.g. from ChIP-seq data) as its input. It first uses these datasets to learn TF activity and then uses TF activities (in place of TF expression) to reconstruct the network. However, it is relatively easy to show that in the best case scenario, this algorithm recovers the prior network (without discovering anything new). While in the practical case in which the algorithms themselves rely on various assumptions and add errors, it finds the original prior network + added errors, but treats the added errors as new discoveries (which is quite dangerous to the research community). I have provided a two-page document attached, focusing on the single-task learning version of the method, describing and showing this flaw. The same problem also exists in the multi-task version of it, but for simplicity I focused here on the single task version.

*•* For the sake of brevity, we will focus our response on the specific claims in the accessory PDF without reproducing it in its entirety 1. The issue here, however, is that **W = P^T^** is trivially a solution to the two-step procedure above. We can see that by replacing this choice of W in Eq 3 to have **X = PA’**. But remembering that **A’** was found by solving **X = PA** (matrix A that satisfies this equation), we can see that **X = PA’** is trivially satisfied. This implies that **W = P^T^** is the solution to the AMUSR two-step procedure.

*•* The reviewer has identified a very valid concern; overfitting is a very real danger for any machine or statistical learning method. In this work, we explore the use of several regularization methods that produce sparse model coefficients (BBSR, StARS-LASSO, and AMuSR) to mitigate overfitting risks. Model selection methods which regularize **W** will result in recovery of a sparse **W** where **W** may or may not have the same structure as **P**.

As a trivial conceptual counterexample to illustrate this point, allow **P** to be a TFs by genes prior matrix where every value is

1. The activity estimate **A’** will then have a rank of 1, where all TF activities are co-linear. As additional predictors provide no additional information, regularization should result in a matrix W which has at most one non-zero entry for each gene, and **W** */*= **P**.

As a second conceptual counterexample to illustrate this point, allow **P** to be a TFs by genes prior matrix where for half of the columns, every value is 0 (as a note, every value is 0 for 43% of the genes in our YEASTRACT prior knowledge network **P**). The corresponding rows of the pseudoinverse **P***^†^* will then also be all zeros. **A’** will be entirely independent of gene *g* which has no non-zero values in the prior matrix, as the gene *g* row in **P***^†^* is all zeros. **A’** will still be a valid predictor matrix, and we can regress expression of gene *g* against **A’** to select TF activities which predict expression of *g*. These selected predictors will be represented as non-zero entries in weight matrix **W** for this gene *g*, and **W** */*= **P**.

As a real-world counterexample, we have performed a number of tests where the expression matrix **X** is replaced with noise (the Noise controls, labeled ’N’ in Figure 4 and Supplemental Figure 4), and we see that performance on held-out genes drops as expected. To further explore this, we have performed a test where we take prior matrix **P** and randomly add false positive edges (reported in Figure 4H), evaluating performance against the gold standard network without holding out any genes from the prior network. If the reviewer’s assertion of circularity is correct, we would expect that **W** would also be filled with false positive edges, and performance would drop dramatically as noise increases. We see that this is not the case.

2. In the best-case scenario, when the algorithms used to solve the twostep procedure above do not use any approximation and do not add errors, one simply recovers matrix P, which we already knew. In the more dangerous practical case, algorithms (those that use different regularization terms with block sparsity, etc.), add errors and find W that is P^T^ + added error. Then, this focuses the attention to the difference of W and P^T^ as new discoveries, while in reality these are simply added errors by algorithms

*•* While some connections added by network inference are undoubtedly spurious, it is not the case that all must be. As a trivial counterexample, imagine three genes (A, B, and C) where genes A and B are strongly positively correlated and genes A and C are strongly negatively correlated. If the prior network contains an edge linking TF-1 to gene A, the activity of TF-1 will correlate with expression of gene A. The activity of TF-1 is then likely to be a useful predictor for the expression of genes B and C, able to explain a substantial amount of the variance observed in the data. An output network **W** where TF-1 is connected to genes A, B, and C is therefore a perfectly reasonable learned network which has new edges which are not present in the prior **P**.

As a real-world counterexample, we note that the results reported in Figures 2-4 are reported on genes for which no prior information was provided. If the reviewer’s assertion that all learned edges are errors by the algorithm is correct, we would expect this to perform no better than the negative controls where labels have been shuffled which are presented in figure 4 (the Shuffled control, labeled ’S’ in Figure 4 and Supplemental Figure 4). We see that this is not the case.

*•* We have shown that the specific mathematical concerns here are addressed in our modeling, but would also like to emphasize that the overall point that this reviewer is making is VERY valid. In the absence of some constraints, which invariably take the form of prior knowledge related to the network structure, the only information available from expression is correlative in nature, yielding networks edge that represent co-expression and have no association with causality. For this reason, the other methods we have benchmarked both incorporate the same prior information SCENIC requires a prior TF-Gene ranking file and TF-Gene binary motif connection file, and CellOracle requires a genes by TFs prior matrix of the same type as the prior we use. We explicitly embed our prior information into a latent TF activity layer.

We believe that it is very important to be clear about this inclusion, as it does create risks (as the reviewer has intuited). The modeling may recover the existing network information that we put in, and little else. This is a systemic problem for the network inference field and highlights the importance of the negative controls which we have included in this work (and which are sadly not ubiquitous when evaluating network inference tools). A comprehensive examination of the circularity problems in the current state of the art for network inference would be a very interesting paper that would add substantially to the literature, but would effectively be an entirely new manuscript and therefore would not fit into this work (I would love to read it if the reviewer were interested in writing it).

### 2.4. Reviewer 3

INTRODUCTION This paper describes Inferelator 3.0, the latest iteration of the Inferelator family of GRN inference algorithms. The latest version differs from the previous version in that it is a Python implementation that uses large-scale parallelization to enable processing of single-cell RNA-Seq (scRNA-Seq) data from up to 10^5^ cells. Otherwise, its basic pipeline and gene-expression modeling methodology are similar to those previously reported in Castro (et al., 2019) from the same lab. The paper does not make any claims about how accurate this new algorithm is compared to Inferelator 2.0, compared to any of the other leading algorithms that are available, or on any absolute scale. Primarily, it describes and evaluates several variants the authors tried before settling on the final Inferelator 3.0 algorithm.

INTEREST TO POTENTIAL READERS It is not clear who the intended audience for this paper is. Logical possibilities would be other researchers working on network inference, potential users of network inference algorithms, and possibly those interested in the biology of the networks produced. The first two groups will be interested only if the paper provides rigorous performance comparisons to other algorithms, including Inferelator 2 and many or most of the leading competitors. Those interested in the biological implications of the networks themselves would require a much deeper analysis of the resulting networks than is currently provided.

MAJOR CLAIMS I was not able to identify any claims other than that certain alternative ways of implementing components of Inferelator 3.0 worked better than others. Looking at the subsections of Results:

1. 2.1 The natural claim here would be that the new Python implementation runs faster than the previous implementation. However, no statements regarding speed or other desirable qualities are made.

2. 2.2. This section compares two expression modeling algorithms the authors considered using, BBSR and StARS-LASSO, and concludes that there is no difference. It also describes AMuSR, published by many of the same authors in 2019, as being better than either of BBSR or StARS-LASSO at dealing with batch effects, so they use AMuSR in Inferelator 3.0. This reports on the authors’ thought process during the design of Inferelator 3.0, but it does not make any claims about Inferelator 3.0 itself.

3. 2.3. This section compares different ways the authors considered putting together a prior network for Inferelator. They observe that two of the methods produce networks that are similar to each other but not similar to the network obtained from the Yeastract database.

This raises questions about the status of Yeastract as a gold standard (see below), but it does not make any specific claims. For example, it does not claim that the Inferelator-prior accessory package they implemented is any better than the existing CellOracle package.

4. 2.4 This section reports on various preprocessing approaches the authors considered when implementing Inferelator 3.0, but it does not make any claims about Inferelator 3.0 itself.

1. 5. 2.5 This section describes how Inferelator 3.0 was run on large datasets comprising mouse single-cell RNA-Seq and ATAC-Seq data. There is no validation of the network. A few sentences are devoted to describing the targets of TFs Egr1 and Atf4. While some readers may be interested in these two TFs, there is little introduction or explanation of why they are of particular interest, among 1500 other TFs.

*•* We thank the reviewer for these comments. The manuscript has been revised to clarify the major claims related to performance in our manuscript, and we have added a number of benchmarks against comparable network inference tools. The reviewer will find this revised manuscript greatly improved by their suggestions for explicit comparisons to other network inference leading methods. Based on this high-quality benchmarking, we claim several specific advantages over other extant network inference methods related to discovering information not present in the prior knowledge network and robustness to noise in that network.

We would like to note that CellOracle is a contemporaneously developed method (it is currently in an alpha state with an associated preprint). Both the inferelator-prior and CellOracle methods for generating prior knowledge networks from motif data are functional, although they generate different prior knowledge networks using different selection criteria. We do not claim that our method for generating prior knowledge networks is superior (their methodology is quite sound). We do claim that our benchmarking (using real-world model-organism data, and testing on a reliable gold standard using information held out of the modeling process) is superior to other network inference benchmarks which do not adhere to good practices for machine learning.

The reviewer’s note that we have not validated the large mouse neuronal network in this work is correct; unfortunately, no rigorous gold standard exists or can be reasonably constructed (a systematic problem which afflicts all work on mammalian network inference). Several network-wide analyses for the mouse neuronal network are provided in Supplemental Figure 7, but the most appropriate validation for this network is experimental. We will add a reference to our manuscript currently in-press which learns new biology by experimentally validating an inferred network.

### RIGOROUS EVIDENCE TO SUPPORT THE CLAIMS

1. Both the Inferelator-internal claims that are made in the current version of the paper and the comparative claims that might be made in a revision require rigorous evaluation of network accuracy. That starts with a clear definition of what it means for a network edge to be correct. For instance, is the binding of the TF in the regulatory DNA of the target gene necessary for correctness? Is it sufficient for correctness? What about if the predicted target changes in expression level when the TF is perturbed? Such a change could be caused by many mechanisms, including mechanisms that are mediated by cell states such as growth rate or metabolic state rather than regulatory networks. Would such changes be considered sufficient for an edge to be correct? Is a change in expression necessary for an edge to be correct?

*•* The reviewer has identified a subtle, but very important point. In the Inferelator framework, an edge is an hypothesis supported by the input data, for which we report summary statistics such as variance explained, and ranked confidence over bootstraps. Our statistical learning explanation is that the framework does not make any assumptions about the interpretation of an edge; this is the purview of the user, who should select a prior knowledge network and a gold standard based on how they expect their biological system to function.

As biologists, we argue that binding to DNA is not necessary, which is fortunate even in a well studied model organism like *Saccharomyces cerevisiae*, the number of TFs which have been conclusively shown to bind DNA is very limited (most in vivo studies of TF binding are, strictly speaking, studies of localization only). We do expect that a TF which causally regulates a gene will localize to that gene in some cellular states. Differential expression of a target gene after a TF is perturbed is also not strictly necessary, although we expect that it will occur in some cellular states. The most accurate answer to the reviewer’s question is that both localization and expression changes are conditionally necessary for a TF gene regulatory edge, but in any arbitrary cellular state it is not necessary that they occur. We have added a clarification on this point to the methods section.

1. 2. Once the intended meaning of the network is made clear, the gold standard for evaluation must match the intended meaning. If binding is considered necessary for correctness, the network should be evaluated against evidence of binding. If functional effect is considered necessary, it should be evaluated against perturbation-response data.

*•* We have selected a prior knowledge network based on criteria that match our biological interpretation. The YEASTRACT prior knowledge network is consists of TF gene edges for which some evidence exists for both localization and for gene expression changes upon TF perturbation. The yeast gold standard which we use was selected for the same criteria, although with a more rigorous requirement for experimental support.

Unfortunately, rigorous celltype-specific genome-scale TF perturbation data is still unavailable for many mammalian systems, and consequently the prior knowledge networks we use from the inferelator-prior pipeline represent predicted TF gene localization. This highlights why we consider experimental validation to be important, as expression changes when we perturb the TF provides strong supporting evidence.

The gold standards the authors use for B. subtilis and S. cerevisiae are described as being curated and/or literature derived. Most edges in Yeastract are derived from a small number of large scale, highthroughput datasets. To the best of my knowledge, no judgments are made as to the quality of the data or the conclusions. Thus, Yeastract is better described as a compilation of (mostly) high-throughput datasets with references, rather than a curated network. While it is literature derived in the sense that there are papers associated with the high-throughput datasets, one should not conclude from this that these literature-derived edges are in any sense more accurate or reliable than high-throughput datasets typically are. And Yeastract includes datasets that are quite old and generally believed to be less reliable than more some more recent datasets.

*•* The reviewer is correct about the YEASTRACT database. While the YEASTRACT prior knowledge network is useful, we do agree that it is not ideally suited for use as a gold standard (largely for the reasons that the reviewer has identified). We therefore use a curated *S. cerevisiae* curated gold standard, as described in https://doi.org/10.1016/j.celrep.2018.03.048.

This gold standard has edges which have evidence from at least three experiments, and which have evidence of both TF localization and gene expression changes after perturbation. We note that this results in a relatively small gold standard network, but as these are (we believe) the highest confidence edges, it is still a valid way to benchmark using ranked measures (e.g. AUPR). We are careful not to use unranked metrics (like Jaccard) when evaluating network performance against this gold standard. We have clarified this in the methods section.

1. 3. Potential readers who are interested in using network inference algorithms need to know which algorithm they should choose, based on accuracy comparison and possibly resource requirements. They also need to know what level of performance they should expect if choose Inferelator 3.0. For example, if they take all edges scoring above some threshold, what fraction of those edges can they expect to be supported by evidence from the gold standard?

*•* A key aspect of this work is how to properly threshold a regulatory network. Metrics like the F1 score or the matthews correlation coefficient proposed here use information from the gold standard or prior knowledge network to identify optimal thresholds for retaining edges. We argue that this principled method of choosing thresholds is superior to selection of some threshold, provided that the network used for scoring is of useful quality. These metrics are valuable as they take into account true positives, false positives, and false negatives in a way that an accuracy measure would not particularly as biological networks are highly imbalanced in positive and negative edges, a situation where an accuracy metric is generally unwise.

To directly address the concern of the reviewer, we have chosen to compare our work to SCENIC and CellOracle as they are the most comparable alternatives for single-cell network inference. The preprocessing (e.g. TF activity) and model selection methods built for older versions of the Inferelator developed in R (e.g. the BBSR model selection method) have been reimplemented in the python-based package which we present here. Based on our extensive software testing framework, we are confident that the output of these reimplemented methods are valid and equivalent to those in the Inferelator 2.0. Our expectation is that the performance of the original R package and the current python package would be very similar when using the same preprocessing and model selection methods, if the out-of-date R package were capable of handling data at this scale (it is not able to handle the staggering number of observations present in single-cell data sets).

### 2.5. Reviewer 4

Major:

1. Using the prior network reconstruction from both CellOracle and Inferelator-prior results in lower AUPR than using one from YEASTRACT. Do the authors have an explanation for this? How accurate/complete does this prior need to be?

*•* This is a very interesting observation on a topic that we’ve considered at some length. To put it simply the strategy of using TF motifs to scan regulatory regions for potential binding will result in poor results for many (or perhaps most) TFs. We suspect the reasons for this are twofold first is that TF motifs themselves are of highly variable reliability. Some TFs (e.g. GAL4) have been extensively studied and the DNA binding has been directly measured, but most TF motifs are derived from ChIP data, which is more indirect. Lower quality motifs will just give poorer estimates of regulation.

The second reason is that both motif-scanning pipelines treat TFs as discrete units that can be modeled in isolation, and that’s just not reflective of the underlying biology in many cases. Some TFs bind cooperatively with other TFs or chromatin readers, and we are unable to account for these types of interaction effects. We also suspect that motifs derived only from ChIP localization data for TFs are less likely to be reliable, as localization is driven by factors other than DNA sequence, but we have not directly tested that hypothesis.

That said, we do not believe that the prior for the inferelator needs to be particularly accurate or complete. TFs for which no accurate predictions have been made in the prior network will unfortunately likely be poorly modeled in the final network, but so long as there is some signal in the noise we believe that modeling performance will be reasonable. We’ve tested this in Figure 4H by taking a the YEASTRACT prior network (which we believe to be the most accurate prior knowledge network we have available) and filling it with randomly generated edges. The resulting network inference performance is quite stable, given that the true prior network edges are outnumbered (up to 10:1) by false positive edges.

2. Interestingly, in applying Inferelator 3.0 to single-cell yeast data, the authors found decreases in performance associated with depth-normalized data, suggesting total counts per cell carries some information in inference. This doesn’t seem to be the case when using BBSR model selection. Can the authors speculate on why this is the case?

*•* This is also a very interesting observation of a subtle effect. As a best-subset regression method that uses the Bayesian Information Criterion, BBSR model selection favors simpler models. There is an initial feature selection based on mutual information which greatly restricts the number of considered features prior to best-subset regression (this is unfortunately necessary as best-subset regression scales exponentially with the number of predictors). Predictor variables (TFs) which are only weakly linked to gene expression through correlation from total count depth are likely to be excluded in this intial filter and not considered during regression. We note that the performance of AMuSR and BBSR are very similar when cell count depth is normalized the difference is that AMuSR performs better on non-depth-normalized expression data, and BBSR performance does not change. Interpretation of the original Figure 4 was needlessly difficult as the y-axis was scaled differently in panels B, C, and D. We have fixed the y-axis scaling in panels B, C, and D in the revised Figure 4 so that they are identical.

3. I’d be interested to understand the limits of Inferelator 3.0 in terms of scalability, which seems to be the main draw of this tool. Reconstruction on 1.3 million single-cells seems impressive (even if divided into 36 clusters), I wonder how long that took, and how scalability compares to previous versions and other single-cell based methods.

*•* This is an excellent question, as this is a lot of data. Our inference approach uses bootstrapping networks (internally ranking network edges by variance explained), and the full network reported in figures 5 & 6 took approximately 3350 cpu-hours to calculate each bootstrap network (around 10 minutes per cpu per gene). We tested this again on the newest version of the Inferelator (which has some additional optimizations) and the newest version of Dask and found it decreased to 1400 cpu-hours (the output is identical). We’re fortunate to have excellent computational resources, but this is a lot of computational time.

We have included a runtime benchmark (without task learning) as Supplemental Figure 5A that compares runtime between the Inferelator and SCENIC, the most scalable of the existing net-work inference tools. At 140k cells, the Inferelator can complete network runs in around an hour, but with equal resources the runtime of SCENIC using GENIE3 is out of a testable range, and SCENIC using GRNBOOST2 dies with cryptic memory errors. Prior iterations of the Inferelator were written for bulk RNA-seq data at a much lower scale. We are quite confident, based on how much of it had to be rewritten to efficiently utilize memory, that earlier versions of the Inferelator are not able to handle 140k cells either. That having been said, we intend to continue developing the Inferelator, as every time we catch up to the size of large single-cell data sets, someone publishes something 10 times larger. There are a number of techniques for scalability that we think we can take advantage of, now that we are built around a powerful (dask) parallelization library.

1. 4. Benchmarking: it would be useful to put this tool in context of others in terms of AUPR, runtime, etc. (i.e. some of the ones mentioned in the background section)

*•* This is a suggestion raised by (all) other reviewers, and we have added several benchmarks. We have included performance benchmarks against the synthetic data in BEELINE (Supplemental Figure 4), and added SCENIC and CellOracle to the yeast singlecell benchmarking in Figure 4. We have also contextualized the advantage of task-based learning by adding the non-task performance against the yeast single-cell benchmark to Supplemental Figure 4. Finally, we have added a runtime benchmark of SCENIC to our runtime benchmarking in Supplemental Figure 5.

Minor

1. missing pointer in line 193

2. References seem to be garbled in lines 284-7

*•* We have corrected these errors.

## 3. Response to Reviewers 2

For consistency, we number the reviewers as they were labeled during the first round of peer review. We’d like to thank Reviewer 1 for their useful suggestions, and have incorporated the changes they advise into Figures 2 and 3. Reviewers 3 and 4 have no new questions or suggestions.

Reviewer 2’s comments restate their first round comments, without considering our response in any meaningful way or providing any new criticism of substance. There are no comments on the manuscript under review, which we note that they have explicitly declined to read, and there are no suggestions for improvements or alternatives. The issues raised by Reviewer 2 are contradicted by the careful tests we have performed and included in the manuscript, both initially and added in response to the first round of peer review. Reviewer 2’s comments comprise an expression of concern by the reviewer regarding an earlier work that was reviewed and published in PLoS Computational Biology.

In response to these concerns, we have edited Figure 2 to improve readability. We have added the performance of the GRNBOOST2 algorithm on bulk *B. subtilis* and *S. cerevisiae* data to Figures 2 and 3 to provide a direct comparison to the results from this method. We have also made minor text edits to clarify methodology.

### 3.1. Reviewer 1

Figure 2A: Type is too small to be read, even at high magnification

*•* We apologize, this figure was poorly scaled. We have reformatted the figure to be easier to read; it is effectively identical content-wise.

2. Figure 2B,C PR curves: Lines for many different methods and datasets are heavily overlapping, making it impossible to compare most of the methods to each other. Rather than plotting all the different holdout sets, it would be more useful to just plot the mean precision as a function of the recall for each method. The fact that the scales on the two axes are different also makes it hard to read. I suggest making the plots square, with both axes being the length of the current vertical axis. The horizontal axis can be cut off at the point where all methods read essentially 0, to save horizontal space if needed.

*•* We have made the aggregate result lines clearer by increasing the alpha transparency on the individual replicates. We have also made these plots square, so that the axes are easier to interpret.

3. All PR plots: Please show a baseline for selecting edges at random (this should be a horizontal line in which precision is independent of recall).

*•* We have added a dashed line that reflects the number of positive network edges out of the total in the gold standard that is used for testing as a convenience. We caution that we would not interpret this as a model baseline. The shuffled controls (labeled ”S” in Figures 2-4 and supplements) should be interpreted as a baseline for selecting edges at random. Calculating the AUPR baseline as the density of the gold standard (the ratio of positive to total edges) is a suboptimal practice for statistical learning in general and network inference in particular. It is generally accurate where the gold standard for testing is uniform in density, or where the model selects edges entirely at random. That is often not the case in biological systems where one TF may have many known target genes and others may have very few, and network inference methods often enforce sparsity in ways that bias edge selection. As an example, consider a gold-standard network of 1000 genes and 100 TFs, where 20 TFs have 100 targets and 80 TFs have 5 targets. The density of this network is 0.024, and the expected number of correct edges if 100 are selected at random is 2.4. However, if the model randomly selects one ”top” target for each TF, the expected number of correct edges is 6. Even though this random model has no predictive ability, it would still perform 250% better than a density baseline.

The density of our yeast gold standard, for example is 0.01442, but the median AUPR of our shuffled controls is 0.01884 (30% higher), and we attribute the difference to non-uniform density within the gold standard. In this work the model AUPR is many times the baseline AUPR and small differences in the baseline AUPR do not meaningfully change the interpretation of our results. There are cases where using density in place of negative controls would suggest that a model has performed over baseline when it has not. The baseline strategy via label shuffling that we use is a best practice which avoids this risk.

1. 4. Also, it is hard to know how good these results are. It would be helpful to have some external point of comparison in this figure.

“These motif-derived prior networks from both the inferelator-prior and CellOracle methods perform well as prior knowledge for GRN inference using the Inferelator pipeline (Figure 3C).” In what sense is an AUPR between 0.1 and 0.15 performing well? Please explain the basis for describing this performance as good. Best would be comparison to other methods.

*•* This is an excellent suggestion. We have added results from the GRNBOOST2 algorithm, which does not use prior information, to Figures 2 and 3. In figure 2, the GRNBOOST2 method performance is above the random change baseline, but is substantially lower than the inferelator performance on held-out genes for 3 of the 4 data sets. We do note that performance on the heavily preprocessed yeast data set 1 (the change over time of the median of three replicates of log2 FC between an experimental and a control channel in a microarray) is much closer to the inferelator than the other data sets. This does suggest that very careful experimental design which minimizes noise may significantly improve the results of correlative network inference methods, although the other three data sets cannot be processed in the same way to test this theory.

In figure 3, The GRNBOOST2 method performance is above the random chance baseline, but only modestly (AUPR of 0.03818 for the GRNBOOST2 method compared to 0.01884 for baseline). Using motif-derived prior networks in the inferelator is therefore significantly better than a correlative network inference method, when scored against genes for which no prior knowledge was provided (hold-outs). Motif-derived prior networks are not as good as using literature database derived prior knowledge (YEASTRACT).

1. 5. Figure 4, legend “. . . two baseline controls; data which has been replaced by Gaussian noise (N) and network inference using shuffled labels in the prior network (S).” Why does Gaussian noise data without count normalization consistently outperform shuffled prior? Shouldn’t both N and S be information-free and thus yield similar results? Is information from the held out portion of the prior network somehow leaking, yielding better-than-random performance even when the gene expression data contain no information?

*•* This is an excellent question (Reviewer 4 asked a related question during the first round of review). The method by which we replace the data with white noise was chosen to retain the mean and standard deviation of each gene, and the total count depth for each cell. There is some information remaining in this control as a result (mean and SD of each gene and count depth of each cell).

We note that when the count depth for each cell is normalized (the right half of Figure 4B-D), the performance of the Noise control (N) drops to equal that of the shuffled control (S). No information about the tested genes was provided to the model (they are true hold-outs; all processing occurs after the split, to ensure that there is no information leakage). Our interpretation is: ”The performance of the randomly generated Noise control

1. (N) is higher than the performance of the shuffled (S) control when counts per cell are not normalized, suggesting that total counts per cell provides additional information during inference.”

### 3.2. Reviewer 2

1. I thank the authors for engaging in this discussion. Below, I have provided my points to their rebuttal, hoping that it helps to improve the methodology.

1. We are always happy to have discussions about our published work, and we appreciate that the reviewer is clear that they are engaged in a post-publication discussion and not in peer review of the manuscript under consideration. We do think it adds greatly to the scientific literature when existing work can be refined. We strongly encourage the reviewer to submit their concerns about Castro 2019 to the PLoS editorial office at. We think that they would be well-received; the underlying issue of model circularity when prior knowledge is embedded is a field-wide problem which would benefit from additional scrutiny.

2. First, I believe the authors agree with my point that W = PT is one of the solutions (if not the only possible solution) of the two-step procedure of the AMUSR.

*•* The reviewer is correct that recovering the prior is a possible solution. It is not clear how the reviewer is defining ’the only possible solution’ to this overdetermined system. We balance model error against sparsity and determine model performance on genes which are not included in P.

The reviewer’s assertion that we must only recover the prior knowledge in P would mean that the genes which we test on would not be incorporated into our final learned network. We see from the results in the manuscript, our model performance is considerably above the control baselines. This contradicts the reviewer’s claim.

3. The authors have mentioned two pathological examples for the choice of P (the prior network): first an all-One matrix and second a low-rank matrix of 0s and ones. Then they have argued that in these cases, W is not equal to P (or to be consistent with my document, W is not equal to PT).

*•* We offered a number of careful controls in the original manuscript, and added additional relevant testing in the revised manuscript. The thought experiments referred to here were only offered in the response to reviewer section as a complement to our manuscript results. They are conceptually easy to grasp counterexamples to specific claims of this reviewer.

With the exception of Figure 4H, which does not score on genes held out of the prior knowledge P (the methods we compare to are not capable of identifying edges not in P, as shown in Figure 4G), all results in Figures 2-4 are shown on genes held out of the prior knowledge P. These results directly show that, when there is no prior knowledge for a gene (every edge for that gene in the prior network is zero), we are still able to identify correct regulators, scored against a gold standard network.

This directly contradicts the concerns of the reviewer, who repeatedly states that the the model output weights W will be equal to the input prior matrix P. If this were true, the model output weights for all genes scored in Figures 2-4 would be zero, and performance would be equal to the model baselines. No suggestion has been made as to how to reconcile the reviewer’s concerns with these results, which are not considered in these comments. We also note that, in response to this concern (and to comments from other reviewers), we have added an experiment (Figure 4H) where a large number of random edges are added to the prior matrix P, and performance is evaluated based on the ability to recover correct, gold standard edges. While we do see a decrease in the Inferelator-AMuSR performance as noise is added to P, the decrease is modest (AUPR decreases from 0.42 to 0.32 when 10 random noise edges are added for every valid edge in P).

If the reviewer’s concerns are valid, we would expect performance to drop dramatically as the noise in P increases, as W would be equal to P and therefore valid edges would be greatly outnumbered by spurious edges. No suggestion has been made as to how to reconcile the reviewer’s concerns with these results, which are not considered in these comments, despite being explicitly referenced in our first response to this reviewer.

4. To answer this, I should first mention that I agree that if one solves the regularization problem that the authors have proposed to solve the two-step procedure of the AMUSR, W will not be equal to PT. This however does not mean that PT is not one (maybe out of several) solutions to the problem. To see this, no matter what the properties of P are, one can replace PT in the equations and see a trivially true statement. For example, consider P to be an all-one matrix (first example). To show this, we only need to show that the choice of W = PT, independent of the nature and structure of PT, is a solution. From my previous document, recall that W is the solution to X = WT A’, where A’ is a matrix that satisfies X = P A’. We only need to show that the choice of W = PT satisfies the equations above. By doing so, we obtain X = PA’ where X =PA’, which is trivially satisfied by definition of A’. So it really does not matter what structure P has. It could be an all-0 matrix or an all-one matrix or any type of example. W = PT is ALWAYS a solution to the two-step procedure of the AMUSR (even though other solutions may exist).

*•* We appreciate that the reviewer sees that their principal concern does not apply to the model which we use in this work (bolded above). Their argument that follows is based on a model formulation that we do not use, for reasons that the reviewer points out, although they incorrectly refer to their toy model as AMUSR. We agree that the toy model that the reviewer is describing is not useful.

*•* 5. I am not claiming that W = PT is the only solution as in the case of pathological examples above, there may be infinitely many solutions. The authors may argue that their regularization allows them to find one of the OTHER solutions of the equations above (which I would agree with). However, given that the W = PT is ALWAYS one of the solutions, and P was obtained from prior information (and is treated as gold standard to guide the rest of inference), why a variation of that (a regularized version of that) is a better solution?

*•* As clearly noted in our work, we have no expectation that the prior information is complete or noise-free. We discuss at length the difficulties in generating prior knowledge networks (it is the topic of several figures and a considerable portion of the results section in this manuscript). There are also specific tests shown that measure the impact of spurious noise in the prior knowledge network on the inference performance.

For the yeast single-cell benchmarking in this manuscript, we do not use the prior knowledge network as a gold standard. The gold standard is a carefully hand-curated set of the highest-confidence interactions, and is not used as the prior knowledge network.

6. Also, I should add that the coefficients of regularization terms influence how strongly the sparsity of S and block sparsity of B (where B + S = W) are imposed. By changing the coefficients, a user can ARTIFICIALLY ensure that the solution W will not be equal to PT. This is what I was referring to in my previous document as the more dangerous case, since an algorithmic artifact will be treated as new discoveries. To make this point more clear, if the sparsity constraints of B and S are not too strict, one can always find a decomposition for P such that S + B = P, where B is “block-sparse” and S is “sparse”. In such a case, W = PT will still be a solution not only to the nonregularized problem, but also to the regularized version of the problem. Depending on how strict these sparsity constraints are, however, such decomposition may or may not be possible. In other words, one can make them so strict to ensure that W found by solving the regularized problem is not equal to PT. This however, is an artifact of how one chooses the regularization coefficients and does not mean that the regularized version of P is more biologically relevant than the nonregularized version of it. In my opinion, the only place that AMUSER can potentially claim usefulness is if P cannot be decomposed as a block sparse matrix B + sparse matrix S (e.g., the pathological all-one example), though again, this depends on some arbitrary definition of sparsity which is imposed by the user through coefficients.

*"* These concerns are specific to the AMuSR multi-task learning method. The coefficients of regularization are set using a principled model selection method, as detailed in this manuscript. They are not chosen arbitrarily by the user. In the case of AMuSR, coefficients are chosen that minimize a variant of the Bayesian information criterion.

We also note that the prior information network P does not have to be the same in each task, as noted and performed in this manuscript. It is not clear how the reviewer’s concerns would be extended to multiple different versions of P with different structures, as well as different expression matrices X with different structures. When applied to joint regression problems, the model weights are not *W* = *B* + *S*, but are as follows, where *n* is the number of regression tasks:

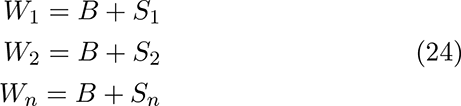

Decomposing model weights into *B* and *S* would be an ill-posed problem if this method were applied to a single regression problem. However, it is explicitly a method for joint regression, where P and X can both vary in each joint regression task, and each task *n* is optimized with different model weights *W_n_*. We have clarified this in Figure 2 and in the supplemental methods section.

Also, even in that case, the authors MUST show that 1) it is often the case that the real-world prior matrices P are not sparse and cannot be written as B+S (however, in my experience most biologically driven prior TRNs are sparse and hence can be written that way)

*•* It is not clear on why the reviewer would suggest that this is a requirement. The basis for all network inference modeling is that biological networks are sparse.

2) the solution of their algorithm is NOT INDEPENDENT of matrix X (mathematically needs to be shown)

*•* We have specifically tested the dependence of our model on matrix X by replacing matrix X with white noise. These results are included in Figure 4 and associated supplements. We do not believe that the standard in this field requires a formalized proof. We note that this standard, applied broadly, would preclude the publication of almost every work that uses iterative optimization instead of a closed-form model solution.

3) a regularized solution that modifies P (given that P itself is a solution) is more biologically relevant than P itself. Again, to reiterate, P is a solution and it comes from prior information (which we are to trust), so why seek a regularized version?

*•* The prior information network should not be trusted. It is incomplete and noisy. A complete and error-free gene regulatory network has not been constructed in any organism.

We do agree that if the prior information network was complete, the modeling that we perform would be superfluous. However, the technology to experimentally determine a complete gene regulatory network does not exist. This is a core motivation for the method presented in our manuscript and is directly covered in the introduction, results, and discussion sections.

Given the issues above, I am not convinced that this narrow case of a matrix P that cannot be written as a sparse + bock-sparse matrix situation is common enough that even if item 2 and 3 is shown to be correct would make AMUSER useful.

*•* We agree that this method would not be appropriate to decompose a single task into a block-sparse and a sparse weight matrix. This is a method not suited to learning from a single prior matrix P and expression matrix X. As such, we do not use it on single-task regression problems.

11. Response: I agree that the solution to the regularized problem will be different IF the coefficients of the regularization term are selected strict enough to make sure P cannot be written as S + B. However, it does not answer why we need to modify P anyways and also does not answer why we should trust a regularized version of P more than P that was our original prior information.

*•* This point has been raised and answered above, and is a core topic of this manuscript. The prior knowledge is incomplete and noisy. The goal is to identify new edges which explain gene expression and to eliminate edges from our prior knowledge that do not explain gene expression.

We quantify ’why we should trust’ the proposed model by holding gene information out of the prior knowledge network and then evaluating our ability to identify regulatory edges for those genes from a hand-curated gold standard. This hold-out testing is a standard in the field of supervised learning. We also include several baseline tests, which are detailed in the manuscript.

In response to the reviewer comments from the first round of revision, we have added a comparison to two other single-cell network inference methods which incorporate prior network knowledge, and ten other single-cell network inference methods which do not. We believe that this method is sufficiently described and tested in this manuscript that a reader can make an informed judgement about its utility.

## Notes

### Competing Interest Statement

The authors have declared no competing interest.

### Summary of Updates

Revision in response to peer review comments. Figures 2 and 3 updated.

