## Supplemental Data 1 (.tar.gz) for "High performance single-cell gene regulatory network inference at scale: The Inferelator 3.0": bsubtilis_mtl_AMUSR2021.py

```
from inferelator import utils
from inferelator.distributed.inferelator_mp import MPControl

from inferelator import workflow
from inferelator.regression.bbsr_multitask import BBSRByTaskRegressionWorkflowMixin
from inferelator.postprocessing.results_processor import InferelatorResults
from inferelator.crossvalidation_workflow import CrossValidationManager

DATA_DIR = '~/fi-inferelator_sc/data/bsubtilis'
OUTPUT_DIR = '~/fi-inferelator_sc/bsubtilis_2021/multitask'
InferelatorResults.curve_data_file_name = "pr_curve.tsv"
CV_SEEDS = list(range(42, 52))

PRIORS_FILE_NAME = 'gold_standard.tsv.gz'
GOLD_STANDARD_FILE_NAME = 'gold_standard.tsv.gz' 
TF_LIST_FILE_NAME = "tf_names.tsv"

BSUBTILIS_1_EXPRESSION = 'expression.tsv.gz'

BSUBTILIS_2_EXPRESSION = 'GSE67023_expression.tsv.gz'

METADATA_1_FILE_NAME = 'meta_data.tsv'
METADATA_2_FILE_NAME = 'GSE67023_meta_data.tsv'
#GENE_METADATA_COLUMN = 'SystematicName'

if __name__ == '__main__':

    utils.Debug.vprint("CV Multitask-AMUSR (5fold)", level=0)
    MPControl.set_multiprocess_engine("dask-cluster")
    MPControl.client.use_default_configuration("rusty_ccb", n_jobs=5)
    WALLTIME = '24:00:00'
    MPControl.client.walltime = WALLTIME
    MPControl.client.add_worker_conda(cmd='~/.local/miniconda3/bin/activate inferelator')
    MPControl.connect()

cv_wrap = CrossValidationManager()
cv_wrap.add_gridsearch_parameter('random_seed', CV_SEEDS)


worker = workflow.inferelator_workflow(regression="amusr", workflow="multitask")
worker.set_file_paths(input_dir=DATA_DIR, output_dir=OUTPUT_DIR,
                      gold_standard_file = GOLD_STANDARD_FILE_NAME,
                      priors_file=PRIORS_FILE_NAME, tf_names_file=TF_LIST_FILE_NAME)
#worker.meta_data_handlers = ['nonbranching','branching']
#worker.shuffle_prior_axis = 0
worker.create_task(task_name="ArrietaOrtiz",
                   input_dir=DATA_DIR,
                   expression_matrix_file=BSUBTILIS_1_EXPRESSION,
                   #expression_matrix_metadata = ['isTs', 'is1stLast', 'prevCol', 'del.t', 'condName'],
                   extract_metadata_from_expression_matrix=True,
                   tf_names_file = "tf_names.tsv",
                   priors_file=PRIORS_FILE_NAME,
                   meta_data_file = METADATA_1_FILE_NAME,
                   expression_matrix_columns_are_genes = False,
                   workflow_type='tfa')
worker.create_task(task_name="Madar",
                   input_dir=DATA_DIR,
                   expression_matrix_file=BSUBTILIS_2_EXPRESSION,
                   #expression_matrix_metadata = ['isTs', 'is1stLast', 'prevCol', 'del.t', 'condName'],
                   extract_metadata_from_expression_matrix=True,
                   tf_names_file = "tf_names.tsv",
                   priors_file=PRIORS_FILE_NAME,
                   meta_data_file = METADATA_2_FILE_NAME,
                   expression_matrix_columns_are_genes = False,
                   workflow_type='tfa')
worker.set_run_parameters(num_bootstraps=5)
worker.set_crossvalidation_parameters(split_gold_standard_for_crossvalidation=True, cv_split_ratio=0.2)
worker.append_to_path("output_dir", "AMUSR")

cv_wrap.workflow = worker
cv_wrap.run()
                   
```
