## Supplemental Data 1 (.tar.gz) for "High performance single-cell gene regulatory network inference at scale: The Inferelator 3.0": bsubtilis_stl_bbsr_DS1_2021.py

```
from inferelator import utils
from inferelator import workflow
from inferelator.distributed.inferelator_mp import MPControl
import os
from inferelator.postprocessing.results_processor import InferelatorResults
# Set verbosity level to "Talky"
utils.Debug.set_verbose_level(1)

# Set the location of the input data and the desired location of the output files

DATA_DIR = '~/fi-inferelator_sc/data/bsubtilis'
OUTPUT_DIR = '~/fi-inferelator_sc/data/bsubtilis_2021/ArrietaOrtiz'

EXPRESSION_FILE_NAME = 'GSE67023_expression.tsv.gz'
#EXPRESSION_FILE_NAME = 'expression.tsv.gz'
PRIORS_FILE_NAME = 'gold_standard.tsv.gz'
GOLD_STANDARD_FILE_NAME = 'gold_standard.tsv.gz'
META_DATA_FILE_NAME = 'GSE67023_meta_data_v2.tsv'
TF_LIST_FILE_NAME = 'tf_names.tsv'
#TFA_FILE = "TF_activities_with_labels_transposed.tsv"
CV_SEEDS = list(range(42, 52))
InferelatorResults.curve_data_file_name = "pr_curve.tsv"

# Multiprocessing uses the pathos implementation of multiprocessing (with dill instead of cPickle)
# This is suited for a single computer but will not work on a distributed cluster

if __name__ == '__main__':
    MPControl.set_multiprocess_engine("multiprocessing")
    MPControl.client.set_processes(10)
    #CONDA_ACTIVATE_PATH = '~/.local/miniconda3/bin/activate inferelator'
    #MPControl.client.use_default_configuration("rusty_ccb", n_jobs=5)
    #MPControl.client.set_processes(os.environ["SLURM_NTASKS"]) 
    #MPControl.client.add_worker_conda("source ~/.local/miniconda3/bin/activate inferelator")
    #MPControl.client.walltime = '48:00:00'
    #MPControl.client.add_worker_env_line('module load slurm')
    #MPControl.client.add_worker_env_line('module load gcc/8.3.0')
    #MPControl.client.add_worker_env_line('source ' + CONDA_ACTIVATE_PATH)
    MPControl.connect()


# Define the general run parameters

def set_up_workflow(wkf):
    wkf.input_dir = DATA_DIR
    wkf.output_dir = OUTPUT_DIR
    wkf.expression_matrix_file = EXPRESSION_FILE_NAME
    wkf.priors_file = PRIORS_FILE_NAME
    wkf.meta_data_file = META_DATA_FILE_NAME
    wkf.tf_names_file = TF_LIST_FILE_NAME
    wkf.gold_standard_file = GOLD_STANDARD_FILE_NAME
    wkf.expression_matrix_columns_are_genes = False
    wkf.num_bootstraps = 5
    return wkf

# Inference with BBSR (crossvalidation)
# Run the regression 10 times and hold 20% of the gold standard out of the priors for testing each time
for random_seed in CV_SEEDS:
    worker = workflow.inferelator_workflow(regression="bbsr", workflow="tfa")
    worker = set_up_workflow(worker)
    worker.split_gold_standard_for_crossvalidation = True
    worker.cv_split_ratio = 0.2
    worker.random_seed = random_seed
    worker.append_to_path('output_dir', 'stl_bbsr_cv_' + str(random_seed))
    worker.run()
    del worker
```
