## Supplemental Data 1 (.tar.gz) for "High performance single-cell gene regulatory network inference at scale: The Inferelator 3.0": yeast_stl_stars_bbsr_D1_2021.py

```
from inferelator import utils
from inferelator.distributed.inferelator_mp import MPControl

from inferelator import workflow
from inferelator.regression.bbsr_multitask import BBSRByTaskRegressionWorkflowMixin
from inferelator.postprocessing.results_processor import InferelatorResults
from inferelator.crossvalidation_workflow import CrossValidationManager


DATA_DIR = '~/fi-inferelator_sc/data/Yeast'
OUTPUT_DIR = '~/fi-inferelator_sc/data/yeast_2021/Hackett'

EXPRESSION_FILE_NAME = 'calico_expression_matrix_log2.tsv.gz'
#META_DATA_FILE_NAME = 'yeast_microarray_meta_data.tsv'
PRIORS_FILE_NAME = 'YEASTRACT_20190713_BOTH.tsv'
GOLD_STANDARD_FILE_NAME = 'gold_standard.tsv'
TF_LIST_FILE_NAME = 'tf_names_restrict.tsv'

GENE_METADATA_FILE_NAME = 'orfs.tsv'
GENE_METADATA_COLUMN = 'SystematicName'

CV_SEEDS = list(range(42,52))
InferelatorResults.curve_data_file_name = "pr_curve.tsv"


if __name__ == '__main__':

    MPControl.set_multiprocess_engine("dask-cluster")
    MPControl.client.use_default_configuration("rusty_ccb", n_jobs=5)
    MPControl.client.add_worker_conda(cmd='~/.local/miniconda3/bin/activate inferelator')
    MPControl.connect()

#cv_wrap = CrossValidationManager()
#cv_wrap.add_gridsearch_parameter('random_seed', CV_SEEDS)


def set_up_workflow(wkf):
    wkf.set_file_paths(input_dir=DATA_DIR,
                       output_dir=OUTPUT_DIR,
                       tf_names_file=TF_LIST_FILE_NAME,
                       #meta_data_file=META_DATA_FILE_NAME,
                       priors_file=PRIORS_FILE_NAME,
                       gold_standard_file=GOLD_STANDARD_FILE_NAME)
    wkf.set_expression_file(tsv=EXPRESSION_FILE_NAME)
    wkf.set_file_properties(expression_matrix_metadata = ['TF', 'strain', 'date', 'restriction', 'mechanism', 'time'],
                            extract_metadata_from_expression_matrix=True,
                            metadata_handler='nonbranching')
    wkf.set_run_parameters(num_bootstraps=5)
    wkf.set_crossvalidation_parameters(split_gold_standard_for_crossvalidation=True, cv_split_ratio=0.2)
    return wkf

for random_seed in CV_SEEDS:
    worker = workflow.inferelator_workflow(regression="bbsr", workflow="tfa")
    worker = set_up_workflow(worker)
    worker.split_gold_standard_for_crossvalidation = True
    worker.cv_split_ratio = 0.2
    worker.random_seed = random_seed
    worker.append_to_path('output_dir', 'stl_bbsr_cv_' + str(random_seed))
    worker.run()
    del worker

for random_seed in CV_SEEDS:
    worker = workflow.inferelator_workflow(regression="stars", workflow="tfa")
    worker = set_up_workflow(worker)
    worker.split_gold_standard_for_crossvalidation = True
    worker.cv_split_ratio = 0.2
    worker.random_seed = random_seed
    worker.append_to_path('output_dir', 'stl_stars_cv_' + str(random_seed))
    worker.run()
    del worker
```
