## Supplemental Data 1 (.tar.gz) for "High performance single-cell gene regulatory network inference at scale: The Inferelator 3.0": figure_3_co.py

```
from inferelator import utils
from inferelator import workflow
from inferelator import crossvalidation_workflow
from inferelator.distributed.inferelator_mp import MPControl


CONDA_ACTIVATE_PATH = '~/.local/anaconda3/bin/activate'

YEASTRACT_PRIOR = "YEASTRACT_20190713_BOTH.tsv"

TF_NAMES = "tf_names_gold_standard.txt"
YEASTRACT_TF_NAMES = "tf_names_yeastract.txt"

EXPRESSION_FILE_NAME = 'yeast_microarray_expression.tsv.gz'
META_DATA_FILE_NAME = 'yeast_microarray_meta_data.tsv'

PRIOR_FILES = edge_files = {"CISBP_1000": 'CO_CISBP_1000_150.tsv.gz',
                            "CISBP_200": 'CO_CISBP_200_50.tsv.gz',
                            "JASPAR_1000": 'CO_JASPAR_1000_150.tsv.gz',
                            "JASPAR_200": 'CO_JASPAR_200_50.tsv.gz',
                            "TRANSFAC_1000": 'CO_TRANSFAC_1000_150.tsv.gz',
                            "TRANSFAC_200": 'CO_TRANSFAC_200_50.tsv.gz',
                            "YEASTRACT": YEASTRACT_PRIOR}

INPUT_DIR = '/mnt/ceph/users/cjackson/inferelator/data/yeast'
OUTPUT_PATH = '/mnt/ceph/users/cjackson/gsj_2020_fig2_CO'

def set_up_workflow(wkf):
    wkf.set_file_paths(input_dir=INPUT_DIR,
                       output_dir=OUTPUT_PATH,
                       tf_names_file="tf_names_yeastract.txt",
                       meta_data_file=META_DATA_FILE_NAME,
                       gold_standard_file="gold_standard.tsv")
    wkf.set_expression_file(tsv=EXPRESSION_FILE_NAME)
    wkf.set_file_properties(expression_matrix_columns_are_genes=False)
    wkf.set_run_parameters(num_bootstraps=5)
    wkf.set_crossvalidation_parameters(split_gold_standard_for_crossvalidation=True, cv_split_ratio=0.2)
    return wkf


def set_up_cv_seeds(wkf):
    cv = crossvalidation_workflow.CrossValidationManager(wkf)
    cv.add_gridsearch_parameter('random_seed', list(range(42, 52)))
    return cv


def set_up_dask(n_jobs=2):
    MPControl.set_multiprocess_engine("dask-cluster")
    MPControl.client.use_default_configuration("rusty_ccb", n_jobs=n_jobs)
    MPControl.client.add_worker_conda("source ~/.local/anaconda3/bin/activate inferelator")
    MPControl.client.add_slurm_command_line("--constraint=broadwell")
    MPControl.connect()


if __name__ == '__main__':
    set_up_dask()

    utils.Debug.vprint("Testing priors", level=0)

    for expt_name, prior_file in PRIOR_FILES.items():

        worker = workflow.inferelator_workflow(regression="bbsr", workflow="tfa")
        set_up_workflow(worker)
        worker.set_file_paths(priors_file=prior_file)
        worker.append_to_path('output_dir', expt_name)
        cv_wrap = set_up_cv_seeds(worker)
        cv_wrap.run()

        del cv_wrap
        del worker
```
