## Supplemental Data 1 (.tar.gz) for "High performance single-cell gene regulatory network inference at scale: The Inferelator 3.0": co_experimental_priornoise.py

```
from inferelator import utils
from inferelator import workflow
from inferelator import crossvalidation_workflow
from inferelator.benchmarking.celloracle import CellOracleWorkflow ,CellOracleRegression
from inferelator.distributed.inferelator_mp import MPControl

import os

YEASTRACT_PRIOR = "YEASTRACT_20190713_BOTH.tsv"
YEASTRACT_TF_NAMES = "tf_names_yeastract.txt"

INPUT_DIR = '/home/cj59/Documents/celloracle'
OUTPUT_PATH = '/scratch/cj59/gsj_2020_celloracle/prior_noise'

utils.Debug.set_verbose_level(1)

MPControl.set_multiprocess_engine("local")
MPControl.connect()

def set_up_workflow(wkf):
    wkf.set_file_paths(input_dir=INPUT_DIR,
                       output_dir=OUTPUT_PATH,
                       priors_file=YEASTRACT_PRIOR,
                       tf_names_file=YEASTRACT_TF_NAMES,
                       gold_standard_file='gold_standard.tsv')
    wkf.set_expression_file(h5ad='GSE144820_GSE125162.h5ad')
    wkf.append_to_path('output_dir', 'experimental')

def set_up_cv_seeds(wkf):
    cv = crossvalidation_workflow.CrossValidationManager(wkf)
    cv.add_gridsearch_parameter('random_seed', list(range(51, 52)))
    return cv


if __name__ == '__main__':

    if 'SLURM_ARRAY_TASK_ID' in os.environ:
        i = int(os.environ['SLURM_ARRAY_TASK_ID'])
        sl = slice(i-1, i)
    else:
        sl = slice(0,5)

    for prior_noise in [0, 0.01, 0.025, 0.05, 0.1][sl]:
        worker = workflow.inferelator_workflow(regression=CellOracleRegression, workflow=CellOracleWorkflow)
        set_up_workflow(worker)
        worker.append_to_path('output_dir', 'noise_' + str(prior_noise))
        worker.set_shuffle_parameters(add_prior_noise=prior_noise) if prior_noise > 0 else None

        cv_wrap = set_up_cv_seeds(worker)
        cv_wrap.run()

        del cv_wrap
        del worker
```
