## Supplemental Data 1 (.tar.gz) for "High performance single-cell gene regulatory network inference at scale: The Inferelator 3.0": figure_4_preprocessing_amusr.py

```
from inferelator import utils
from inferelator import workflow
from inferelator import crossvalidation_workflow
from inferelator.preprocessing import single_cell
from inferelator.distributed.inferelator_mp import MPControl


CONDA_ACTIVATE_PATH = '~/.local/anaconda3/bin/activate'
EXPRESSION_MATRIX_METADATA = ['Genotype', 'Genotype_Group', 'Replicate', 'Condition', 'tenXBarcode']

YEASTRACT_PRIOR = "YEASTRACT_20190713_BOTH.tsv"

TF_NAMES = "tf_names_gold_standard.txt"
YEASTRACT_TF_NAMES = "tf_names_yeastract.txt"

INPUT_DIR = '/mnt/ceph/users/cjackson/inferelator/data/yeast'
OUTPUT_PATH = '/mnt/ceph/users/cjackson/gsj_2021_amusr_fig4'

utils.Debug.set_verbose_level(1)


def set_up_workflow(wkf):
    wkf.set_file_paths(input_dir=INPUT_DIR,
                       output_dir=OUTPUT_PATH,
                       priors_file=YEASTRACT_PRIOR,
                       tf_names_file=YEASTRACT_TF_NAMES,
                       gold_standard_file='gold_standard.tsv')

    task = worker.create_task(task_name="Jackson_2019",
                              workflow_type="single-cell",
                              count_minimum=0.05,
                              tasks_from_metadata=True,
                              meta_data_task_column="Condition")
    task.set_expression_file(h5ad='GSE144820_GSE125162.h5ad')

    wkf.set_crossvalidation_parameters(split_gold_standard_for_crossvalidation=True,
                                       cv_split_ratio=0.2)
    wkf.set_run_parameters(num_bootstraps=5)
    wkf.set_count_minimum(0.05)


def set_up_cv_seeds(wkf, seeds=list(range(42, 52))):
    cv = crossvalidation_workflow.CrossValidationManager(wkf)
    cv.add_gridsearch_parameter('random_seed', seeds)
    return cv


def set_up_dask(n_jobs=5):
    MPControl.set_multiprocess_engine("dask-cluster")
    MPControl.client.use_default_configuration("rusty_ccb", n_jobs=n_jobs)
    MPControl.client.add_worker_conda("source ~/.local/anaconda3/bin/activate inferelator")
    MPControl.client.add_slurm_command_line("--constraint=broadwell")
    MPControl.connect()


if __name__ == '__main__':
    set_up_dask()

    utils.Debug.vprint("Testing preprocessing", level=0)

    # Figure 5D: BBSR By Task Learning

    worker = workflow.inferelator_workflow(regression="amusr", workflow="multitask")
    set_up_workflow(worker)
    worker.append_to_path('output_dir', 'figure_4_count')
    cv_wrap = set_up_cv_seeds(worker)
    cv_wrap.run()

    del cv_wrap
    del worker

    worker = workflow.inferelator_workflow(regression="amusr", workflow="multitask")
    set_up_workflow(worker)
    worker.add_preprocess_step("log2")
    worker.append_to_path('output_dir', 'figure_4_log2')
    cv_wrap = set_up_cv_seeds(worker)
    cv_wrap.run()

    del cv_wrap
    del worker

    worker = workflow.inferelator_workflow(regression="amusr", workflow="multitask")
    set_up_workflow(worker)
    worker.add_preprocess_step("ftt")
    worker.append_to_path('output_dir', 'figure_4_fft')
    cv_wrap = set_up_cv_seeds(worker)
    cv_wrap.run()

    del cv_wrap
    del worker

    worker = workflow.inferelator_workflow(regression="amusr", workflow="multitask")
    set_up_workflow(worker)
    worker.add_preprocess_step(single_cell.normalize_expression_to_median)
    worker.append_to_path('output_dir', 'figure_4_median')
    cv_wrap = set_up_cv_seeds(worker)
    cv_wrap.run()

    del cv_wrap
    del worker

    worker = workflow.inferelator_workflow(regression="amusr", workflow="multitask")
    set_up_workflow(worker)
    worker.add_preprocess_step(single_cell.normalize_expression_to_median)
    worker.add_preprocess_step("log2")
    worker.append_to_path('output_dir', 'figure_4_median_log2')
    cv_wrap = set_up_cv_seeds(worker)
    cv_wrap.run()

    del cv_wrap
    del worker

    worker = workflow.inferelator_workflow(regression="amusr", workflow="multitask")
    set_up_workflow(worker)
    worker.add_preprocess_step(single_cell.normalize_expression_to_median)
    worker.add_preprocess_step("ftt")
    worker.append_to_path('output_dir', 'figure_4_median_fft')
    cv_wrap = set_up_cv_seeds(worker)
    cv_wrap.run()

    del cv_wrap
    del worker
```
