## Supplemental Data 1 (.tar.gz) for "High performance single-cell gene regulatory network inference at scale: The Inferelator 3.0": scenic_experimental.py

```
from inferelator import utils
from inferelator import workflow
from inferelator import crossvalidation_workflow
from inferelator.benchmarking.scenic import SCENICWorkflow, SCENICRegression
from inferelator.distributed.inferelator_mp import MPControl

YEASTRACT_PRIOR = "YEASTRACT_20190713_BOTH.tsv"
YEASTRACT_TF_NAMES = "tf_names_yeastract.txt"

INPUT_DIR = '/home/cj59/Documents/scenic'
OUTPUT_PATH = '/scratch/cj59/gsj_2020_scenic'

utils.Debug.set_verbose_level(1)

def set_up_workflow(wkf):
    wkf.set_file_paths(input_dir=INPUT_DIR,
                       output_dir=OUTPUT_PATH,
                       priors_file=YEASTRACT_PRIOR,
                       tf_names_file=YEASTRACT_TF_NAMES,
                       gold_standard_file='gold_standard.tsv')
    wkf.set_expression_file(h5ad='GSE144820_GSE125162.h5ad')
    wkf.dask_temp_path = '/scratch/cj59/'
    wkf.set_crossvalidation_parameters(split_gold_standard_for_crossvalidation=True,
                                       cv_split_ratio=0.2)


def set_up_cv_seeds(wkf):
    cv = crossvalidation_workflow.CrossValidationManager(wkf)
    cv.add_gridsearch_parameter('random_seed', list(range(42, 52)))
    return cv

    
if __name__ == '__main__':

    MPControl.set_multiprocess_engine("dask-cluster")
    MPControl.client.use_default_configuration("greene", n_jobs=2)
    MPControl.client.add_worker_conda("source /scratch/cgsb/gresham/no_backup/Chris/.conda/bin/activate scenic")
    MPControl.connect()

    worker = workflow.inferelator_workflow(regression=SCENICRegression, workflow=SCENICWorkflow)
    set_up_workflow(worker)
    worker.append_to_path('output_dir', 'experimental')
    
    cv_wrap = set_up_cv_seeds(worker)
    cv_wrap.run()
```
