## Supplemental Data 1 (.tar.gz) for "High performance single-cell gene regulatory network inference at scale: The Inferelator 3.0": figure_6.py

```
from inferelator import utils
from inferelator import workflow
from inferelator.preprocessing import single_cell
from inferelator.distributed.inferelator_mp import MPControl

import os
import csv
import time
from dask.distributed import performance_report

utils.Debug.set_verbose_level(1)


CONDA_ACTIVATE_PATH = '~/miniconda3/bin/activate'
INPUT_DIR =  '/mnt/ceph/users/sysbio/GSM/data'
OUTPUT_DIR = '/mnt/ceph/users/cjackson/gsj_2021_fig6'
TF_NAMES =   'TF_e18.tsv'
OUT_PREFIX = 'APR14_counts_10'
GS =         'gold_std_apr_8_rec.tsv'

PRIORS =     {'EXC_P':'priors/E18_EXC_apr_8_rec.tsv',
              'IN_P':'priors/E18_IN_apr_8_rec.tsv',
              'GL_P':'priors/E18_GL_apr_8_rec.tsv'}

TASKS =      {'EXC_CN_1': {'file':'l1p_filtered/EXC_CN_1_Apr9.h5ad','tasker':None, 'prior':'EXC_P'},
              'EXC_CN_2': {'file':'l1p_filtered/EXC_CN_2_Apr9.h5ad','tasker':None, 'prior':'EXC_P'},
              'EXC_CN_3': {'file':'l1p_filtered/EXC_CN_3_Apr9.h5ad','tasker':None, 'prior':'EXC_P'},
              'EXC_CN_4': {'file':'l1p_filtered/EXC_CN_4_Apr9.h5ad','tasker':None, 'prior':'EXC_P'},
              'EXC_CN_5': {'file':'l1p_filtered/EXC_CN_5_Apr9.h5ad','tasker':None, 'prior':'EXC_P'},
              'EXC_IT_1': {'file':'l1p_filtered/EXC_IT_1_Apr9.h5ad','tasker':None, 'prior':'EXC_P'},
              'EXC_IT_2': {'file':'l1p_filtered/EXC_IT_2_Apr9.h5ad','tasker':None, 'prior':'EXC_P'},
              'EXC_IT_3': {'file':'l1p_filtered/EXC_IT_3_Apr9.h5ad','tasker':None, 'prior':'EXC_P'},
              'EXC_IT_4': {'file':'l1p_filtered/EXC_IT_4_Apr9.h5ad','tasker':None, 'prior':'EXC_P'},
              'EXC_IT_5': {'file':'l1p_filtered/EXC_IT_5_Apr9.h5ad','tasker':None, 'prior':'EXC_P'},
              'EXC_IT_6': {'file':'l1p_filtered/EXC_IT_6_Apr9.h5ad','tasker':None, 'prior':'EXC_P'},
              'IN_MGE_0': {'file':'l1p_filtered/IN_MGE_0_Apr9.h5ad','tasker':None, 'prior':'IN_P'},
              'IN_CGE_0': {'file':'l1p_filtered/IN_CGE_0_Apr9.h5ad','tasker':None, 'prior':'IN_P'},
              'IN_LGE_0': {'file':'l1p_filtered/IN_LGE_0_Apr9.h5ad','tasker':None, 'prior':'IN_P'},
              'IN_LGE_1': {'file':'l1p_filtered/IN_LGE_1_Apr9.h5ad','tasker':None, 'prior':'IN_P'},
              'GL_AC_0': {'file':'l1p_filtered/GL_AC_0_Apr9.h5ad','tasker':None, 'prior':'GL_P'},
              'GL_AC_1': {'file':'l1p_filtered/GL_AC_1_Apr9.h5ad','tasker':None, 'prior':'GL_P'},
              'GL_AC_2': {'file':'l1p_filtered/GL_AC_2_Apr9.h5ad','tasker':None, 'prior':'GL_P'},
              'GL_OG_0': {'file':'l1p_filtered/GL_OG_0_Apr9.h5ad','tasker':None, 'prior':'GL_P'},
              'GL_OG_1': {'file':'l1p_filtered/GL_OG_1_Apr9.h5ad','tasker':None, 'prior':'GL_P'},
              'GL_OG_3': {'file':'l1p_filtered/GL_OG_3_Apr9.h5ad','tasker':None, 'prior':'GL_P'},
              'GL_OG_4': {'file':'l1p_filtered/GL_OG_4_Apr9.h5ad','tasker':None, 'prior':'GL_P'},
              'GL_CR_0': {'file':'l1p_filtered/GL_CR_0_Apr9.h5ad','tasker':None, 'prior':'GL_P'},
              'GL_CR_1': {'file':'l1p_filtered/GL_CR_1_Apr9.h5ad','tasker':None, 'prior':'GL_P'},
              'GL_0': {'file':'l1p_filtered/GL_0_Apr9.h5ad','tasker':None, 'prior':'GL_P'},
              'GL_1': {'file':'l1p_filtered/GL_1_Apr9.h5ad','tasker':None, 'prior':'GL_P'},
              'GL_2': {'file':'l1p_filtered/GL_2_Apr9.h5ad','tasker':None, 'prior':'GL_P'},
              'GL_3': {'file':'l1p_filtered/GL_3_Apr9.h5ad','tasker':None, 'prior':'GL_P'},
              'GL_MCG_0': {'file':'l1p_filtered/GL_MCG_0_Apr9.h5ad','tasker':None, 'prior':'GL_P'},
              'GL_MCG_1': {'file':'l1p_filtered/GL_MCG_1_Apr9.h5ad','tasker':None, 'prior':'GL_P'},
              'VC_EN_1': {'file':'l1p_filtered/VC_EN_1_Apr9.h5ad','tasker':None, 'prior':'GL_P'},
              'VC_EN_0': {'file':'l1p_filtered/VC_EN_0_Apr9.h5ad','tasker':None, 'prior':'GL_P'},
              'VC_EN_2': {'file':'l1p_filtered/VC_EN_2_Apr9.h5ad','tasker':None, 'prior':'GL_P'},
              'VC_PC_0': {'file':'l1p_filtered/VC_PC_0_Apr9.h5ad','tasker':None, 'prior':'GL_P'},
              'VC_PC_2': {'file':'l1p_filtered/VC_PC_2_Apr9.h5ad','tasker':None, 'prior':'GL_P'},
              'VC_VLMC': {'file':'l1p_filtered/VC_VLMC_0_Apr9.h5ad','tasker':None, 'prior':'GL_P'}}

MPControl.set_multiprocess_engine("dask-cluster")
MPControl.client.use_default_configuration("rusty_rome", n_jobs=5)
MPControl.client.set_cluster_params(local_workers=1)
MPControl.client.add_worker_conda("source ~/.local/anaconda3/bin/activate inferelator")
MPControl.client.add_slurm_command_line("--constraint=rome")
MPControl.client.set_job_size_params(walltime="168:00:00")
MPControl.connect()

worker = workflow.inferelator_workflow(regression="amusr", workflow="multitask")
worker.set_file_paths(input_dir=INPUT_DIR, output_dir=OUTPUT_DIR, tf_names_file=TF_NAMES,gold_standard_file=GS)
worker.append_to_path('output_dir', OUT_PREFIX)
worker.count_minimum = None
worker.curve_data_file_name = '{}_curve.tsv'.format(OUT_PREFIX)
worker.set_run_parameters(use_mkl=True, num_bootstraps=10)

for task in TASKS.keys():
    TASKS[task]['tasker'] = worker.create_task(task_name=task,
                                               tf_names_file=TF_NAMES,
                                               tasks_from_metadata=False,
                                               expression_matrix_columns_are_genes=True,
                                               extract_metadata_from_expression_matrix=False,
                                               workflow_type="single-cell",
                                               priors_file=PRIORS[TASKS[task]['prior']])
    TASKS[task]['tasker'].set_expression_file(h5ad=TASKS[task]['file'], h5_layer='counts')
    TASKS[task]['tasker'].set_count_minimum(0.05)
    TASKS[task]['tasker'].add_preprocess_step(single_cell.log2_data)


result = worker.run()
```
