## Supplemental Data 1 (.tar.gz) for "High performance single-cell gene regulatory network inference at scale: The Inferelator 3.0": supp_fig_4_profiling_bbsr.py

```
from inferelator import utils
from inferelator import workflow
from inferelator import crossvalidation_workflow
from inferelator.distributed.inferelator_mp import MPControl

from dask.distributed import performance_report
import os, subprocess, signal, time, csv, gc
import numpy as np

CONDA_ACTIVATE_PATH = '~/.local/anaconda3/bin/activate'

PRIOR_FILE = "E18_EXC_apr_8_rec.tsv"
TF_NAMES = "TF_e18.tsv"

INPUT_DIR = '/mnt/ceph/users/sysbio/chris'
OUTPUT_PATH = '/mnt/ceph/users/cjackson/gsj_2020_profile_bbsr'

utils.Debug.set_verbose_level(1)


def set_up_workflow(wkf):
    wkf.set_file_paths(input_dir=INPUT_DIR,
                       output_dir=OUTPUT_PATH,
                       priors_file=PRIOR_FILE,
                       tf_names_file=TF_NAMES,
                       gold_standard_file=PRIOR_FILE)

    wkf.set_expression_file(h5ad='144k_by_10384_EXC_IT_1.h5ad')

    wkf.set_crossvalidation_parameters(split_gold_standard_for_crossvalidation=True,
                                       cv_split_ratio=0.2)
    wkf.set_run_parameters(num_bootstraps=5, use_mkl=True)
    wkf.set_count_minimum(0.01)


MPControl.set_multiprocess_engine("dask-cluster")
MPControl.client.use_default_configuration("rusty_ccb", n_jobs=5)
MPControl.client.add_worker_conda("source ~/.local/anaconda3/bin/activate inferelator_new")
MPControl.client.add_slurm_command_line("--constraint=broadwell")
MPControl.client.set_cluster_params(local_workers=0)
MPControl.client.set_job_size_params(walltime="168:00:00")
MPControl.connect()
MPControl.client.is_dask()


class DownsampleDataWorkflow(workflow._factory_build_inferelator(regression="bbsr", workflow="single-cell")):

    sample_ratio = None
    sample_seed = 1000

    def startup_run(self):
        super(DownsampleDataWorkflow, self).startup_run()

        if self.sample_ratio == 1.:
            return

        rgen = np.random.default_rng(self.sample_seed)

        n_keep = int(self.data.num_obs * self.sample_ratio)
        n_keep = 1 if n_keep < 1 else n_keep

        self.data.get_random_samples(n_keep, random_gen=rgen, inplace=True, with_replacement=False)

if __name__ == '__main__':

    os.makedirs(OUTPUT_PATH, exist_ok=True)
    with open(os.path.join(OUTPUT_PATH, "downsample_performance_bbsr.tsv"), "a+") as out_fh:

        csv_handler = csv.writer(out_fh, delimiter="\t", lineterminator="\n", quoting=csv.QUOTE_NONE)
        csv_handler.writerow(["Ratio", "Seed", "Num_Cells", "Time", "AUPR", "F1", "MCC"])

        for ratio in [0.005, 0.01, 0.1, 1.0]:

            for seed in range(49, 52):

                worker = DownsampleDataWorkflow()
                set_up_workflow(worker)
                worker.add_preprocess_step("log2")

                worker.set_output_file_names(network_file_name=None, confidence_file_name=None,
                                            nonzero_coefficient_file_name=None,
                                            pdf_curve_file_name=None,
                                            curve_data_file_name=None)
                                            
                worker.set_run_parameters(num_bootstraps=5)
                worker.append_to_path('output_dir', 'network_outputs')
                worker.sample_ratio = ratio
                worker.random_seed = seed
                worker.sample_seed = seed + 1000

                performance_filename = os.path.join(OUTPUT_PATH, "perf_" + str(ratio) + "_" + str(seed))

                #https://stackoverflow.com/questions/4789837/how-to-terminate-a-python-subprocess-launched-with-shell-true
                cmd = "python -m inferelator.utils.profiler -p {pid} -o {pfn}".format(pid=os.getpid(), pfn=performance_filename + "_mem.tsv")
                memory_monitor = subprocess.Popen(cmd, stdout=subprocess.PIPE, shell=True, preexec_fn=os.setsid) 

                start_time = time.time()
                with performance_report(filename=performance_filename + ".html"):
                    result = worker.run()
                
                csv_row = [str(ratio), str(seed), str(worker._num_obs), '%.1f' % (time.time() - start_time)]
                csv_row += [result.all_scores[n] for n in result.all_names]

                csv_handler.writerow(csv_row)

                del worker
                del result

                time.sleep(1)
                os.killpg(os.getpgid(memory_monitor.pid), signal.SIGTERM)

                MPControl.client.client.restart()
                gc.collect()

                out_fh.flush()
```
