## Supplemental Data 1 (.tar.gz) for "High performance single-cell gene regulatory network inference at scale: The Inferelator 3.0": README

Supplemental Data 1

Gold Standard Files:

./Figure2/B_subtilis_gold_standard.tsv.gz
	Bacillus subtilis gold standard TSV. 3148 unsigned edges [4891 genes x 239 TFs].

./Figure2/S_cerevisiae_gold_standard.tsv.gz
	Saccharomyces cerevisiae gold standard TSV. 1403 signed edges [993 genes x 98 TFs].

Network Prior Files:

./Figure3/CO_CISBP_200_50.tsv.gz
	Yeast prior connectivity matrix TSV generated with CellOracle and the CIS-BP motif database. 34381 edges [6037 genes x 193 TFs].

./Figure3/CO_JASPAR_200_50.tsv.gz
Yeast prior connectivity matrix TSV generated with CellOracle and the JASPAR motif database. 15625 edges [6037 genes x 137 TFs].

./Figure3/CO_TRANSFAC_200_50.tsv.gz
	Yeast prior connectivity matrix TSV generated with CellOracle and the TRANSFAC motif database. 24601 edges [6037 genes x 156 TFs].

./Figure3/Scer_cisbp_200_50_edge_matrix.tsv.gz
Yeast prior connectivity matrix TSV generated with inferelator-prior and the CIS-BP motif database. 16191 edges [6037 genes x 212 TFs].

./Figure3/Scer_jaspar_200_50_edge_matrix.tsv.gz
	Yeast prior connectivity matrix TSV generated with inferelator-prior and the JASPAR motif database. 15625 edges [6037 genes x 175 TFs].

./Figure3/Scer_transfac_200_50_edge_matrix.tsv.gz
	Yeast prior connectivity matrix TSV generated with inferelator-prior and the TRANSFAC motif database. 13817 edges [6037 genes x 175 TFs].

./Figure3/YEASTRACT_20190713_BOTH.tsv.gz
	Yeast prior connectivity matrix TSV downloaded from YEASTRACT July 19th 2019. 11486 edges [6885 genes x 152 TFs].

./Figure5/E18_EXC_apr_8_rec.tsv.gz
	Mouse E18 excitatory neuron prior connectivity matrix TSV generated with inferelator-prior and the TRANSFAC motif database. 39272 edges [17526 genes x 587 TFs]

./Figure5/E18_IN_apr_8_rec.tsv.gz
	Mouse E18 interneuron prior connectivity matrix TSV generated with inferelator-prior and the TRANSFAC motif database. 40363 edges [17526 genes x 587 TFs]

./Figure5/E18_GL_apr_8_rec.tsv.gz
	Mouse E18 glial prior connectivity matrix TSV generated with inferelator-prior and the TRANSFAC motif database. 39272 edges [17526 genes x 588 TFs]

Inferelator Workflow Scripts:

./InferelatorScripts/*
	Python scripts to run network inference with the Inferelator v0.5.6 to generate networks
