## Supplementary figures and images for "High performance single-cell gene regulatory network inference at scale: The Inferelator 3.0"

### combined_metrics.pdf

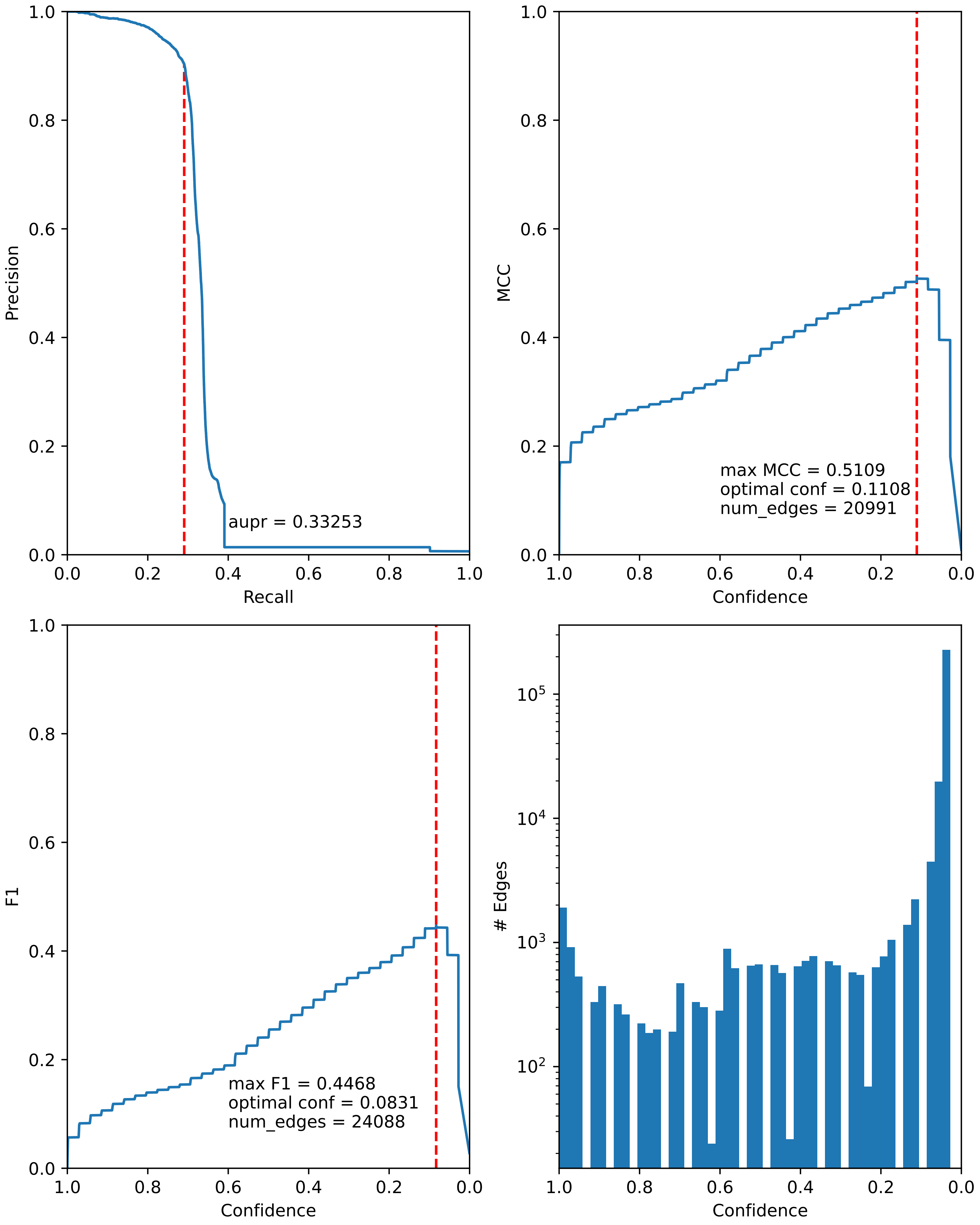

### combined_metrics.pdf

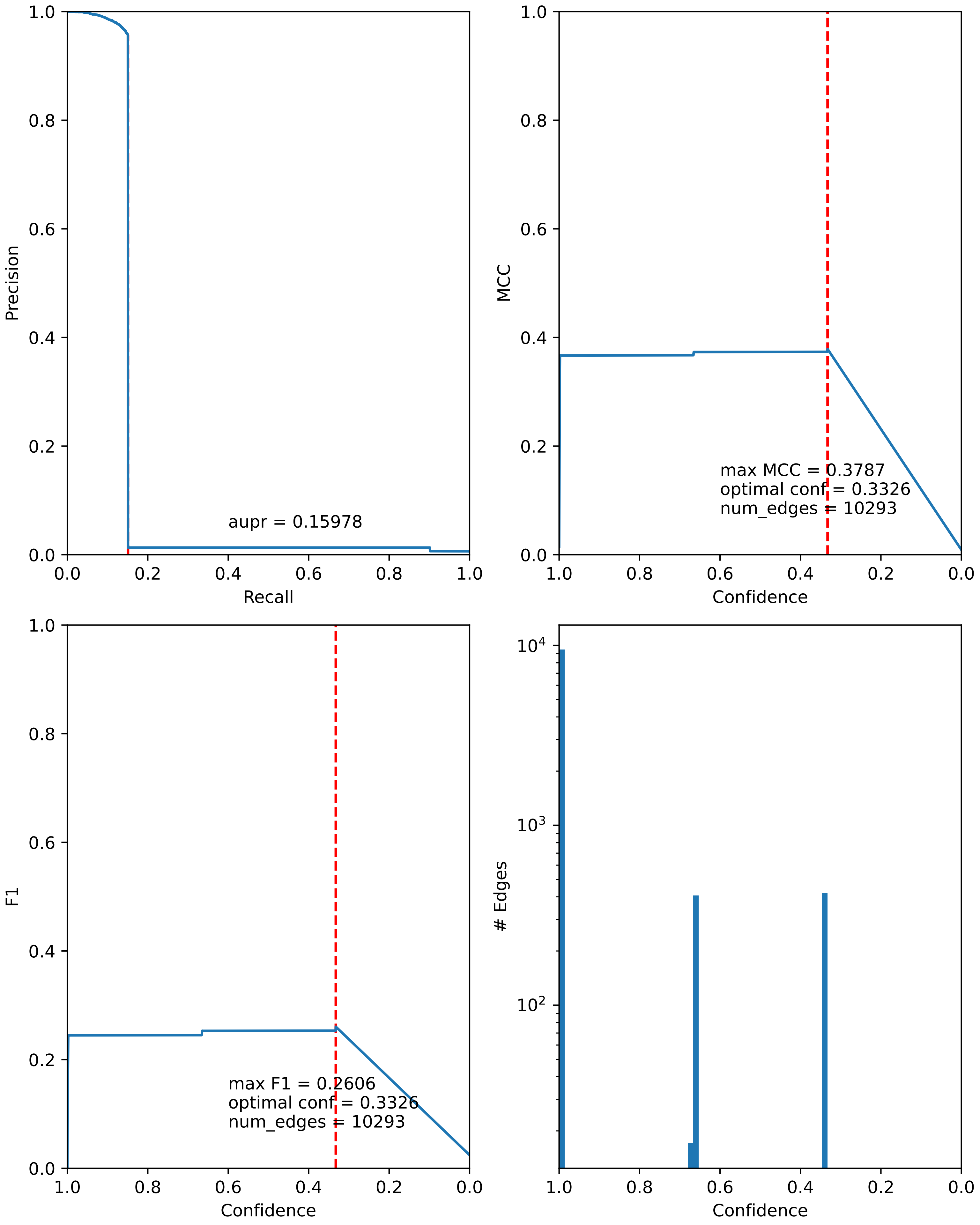

### combined_metrics.pdf

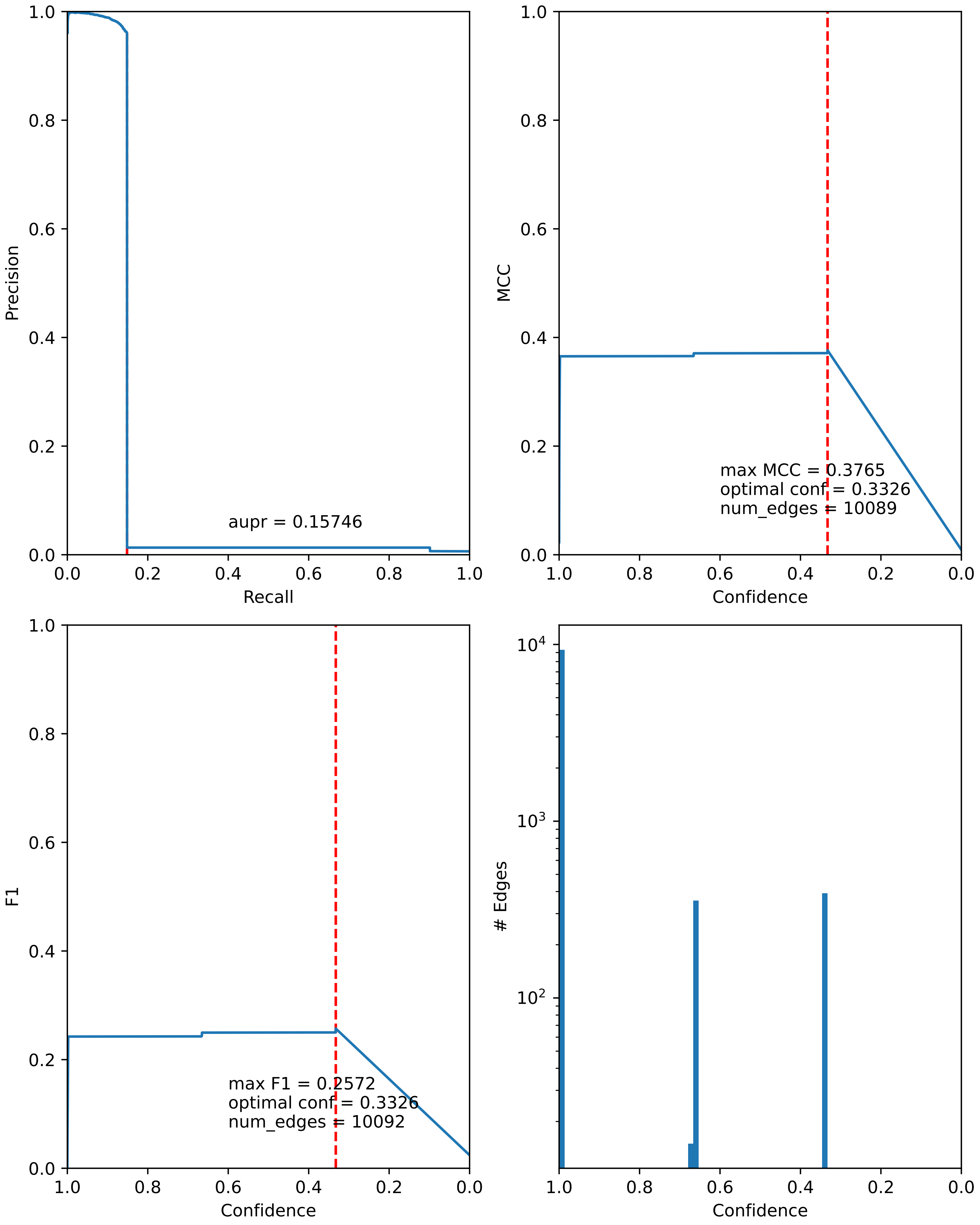

### combined_metrics.pdf

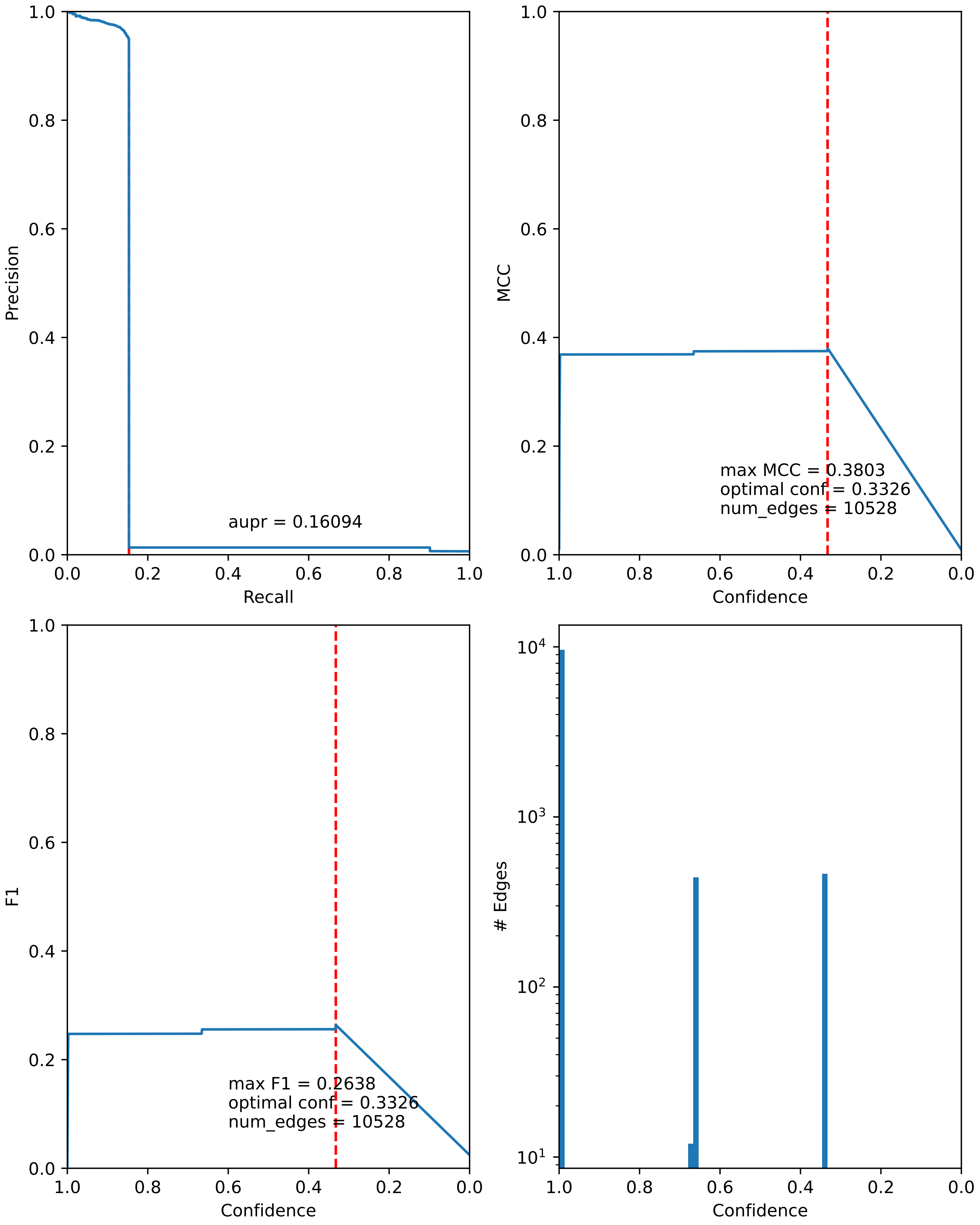

### combined_metrics.pdf

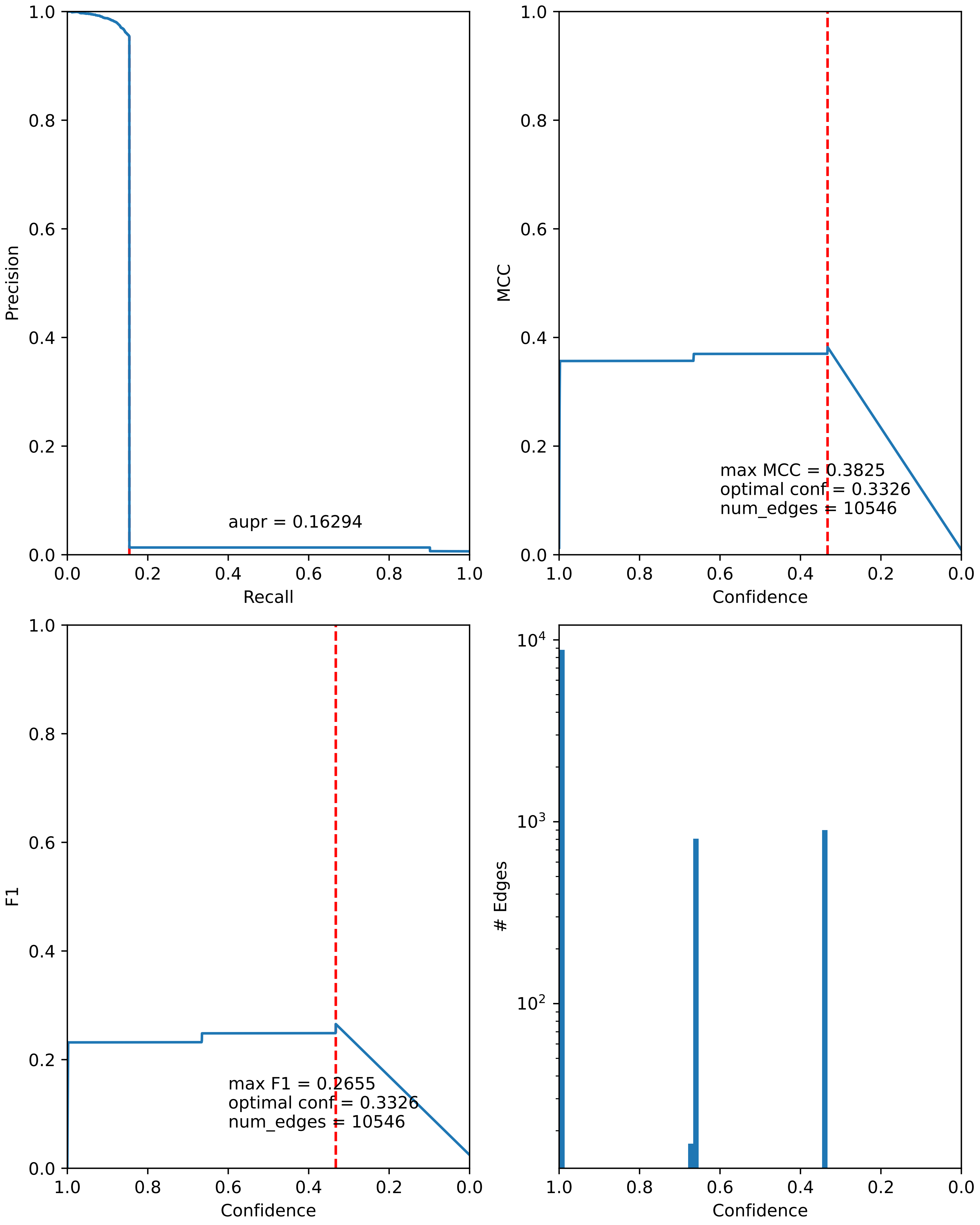

### combined_metrics.pdf

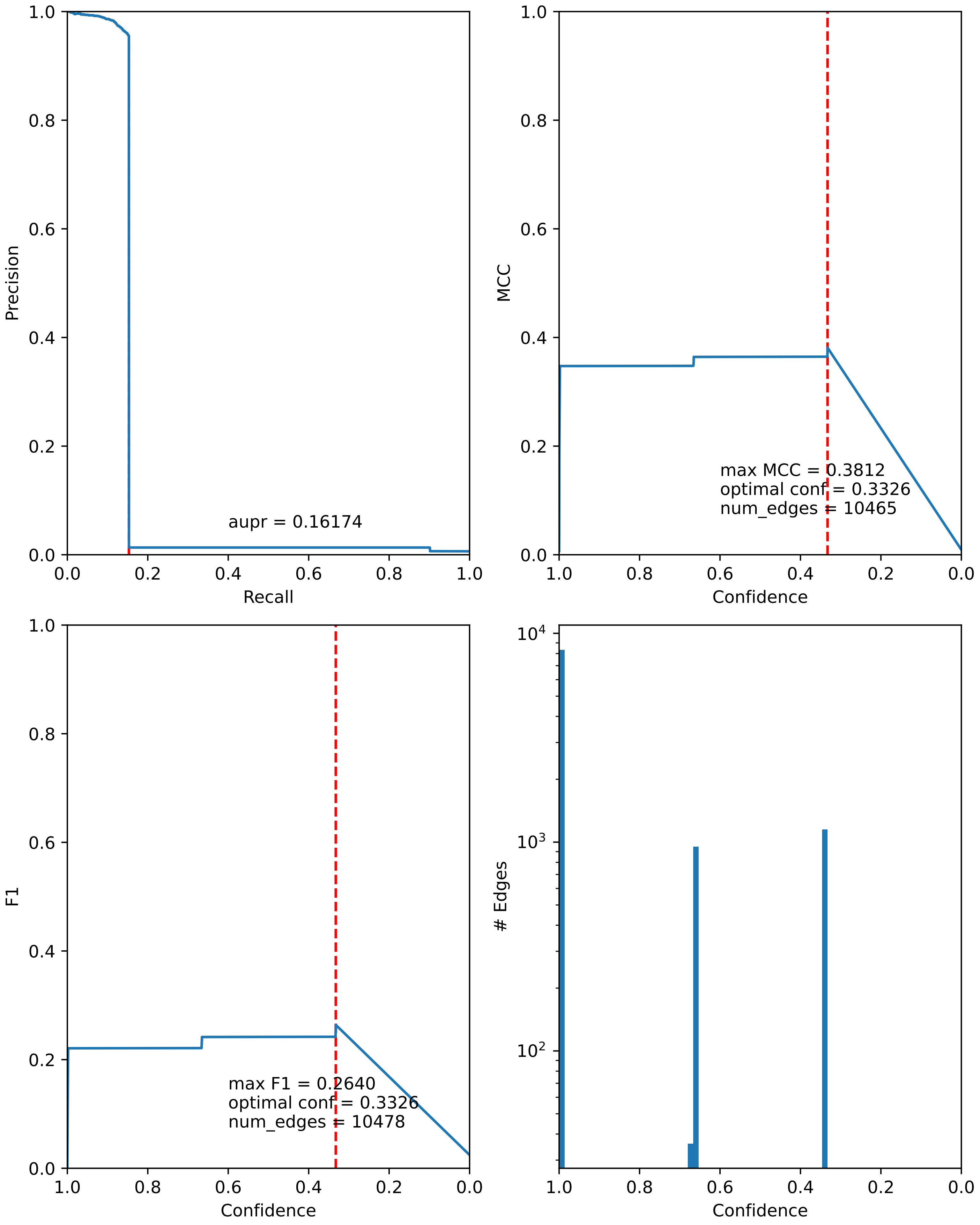

### combined_metrics.pdf

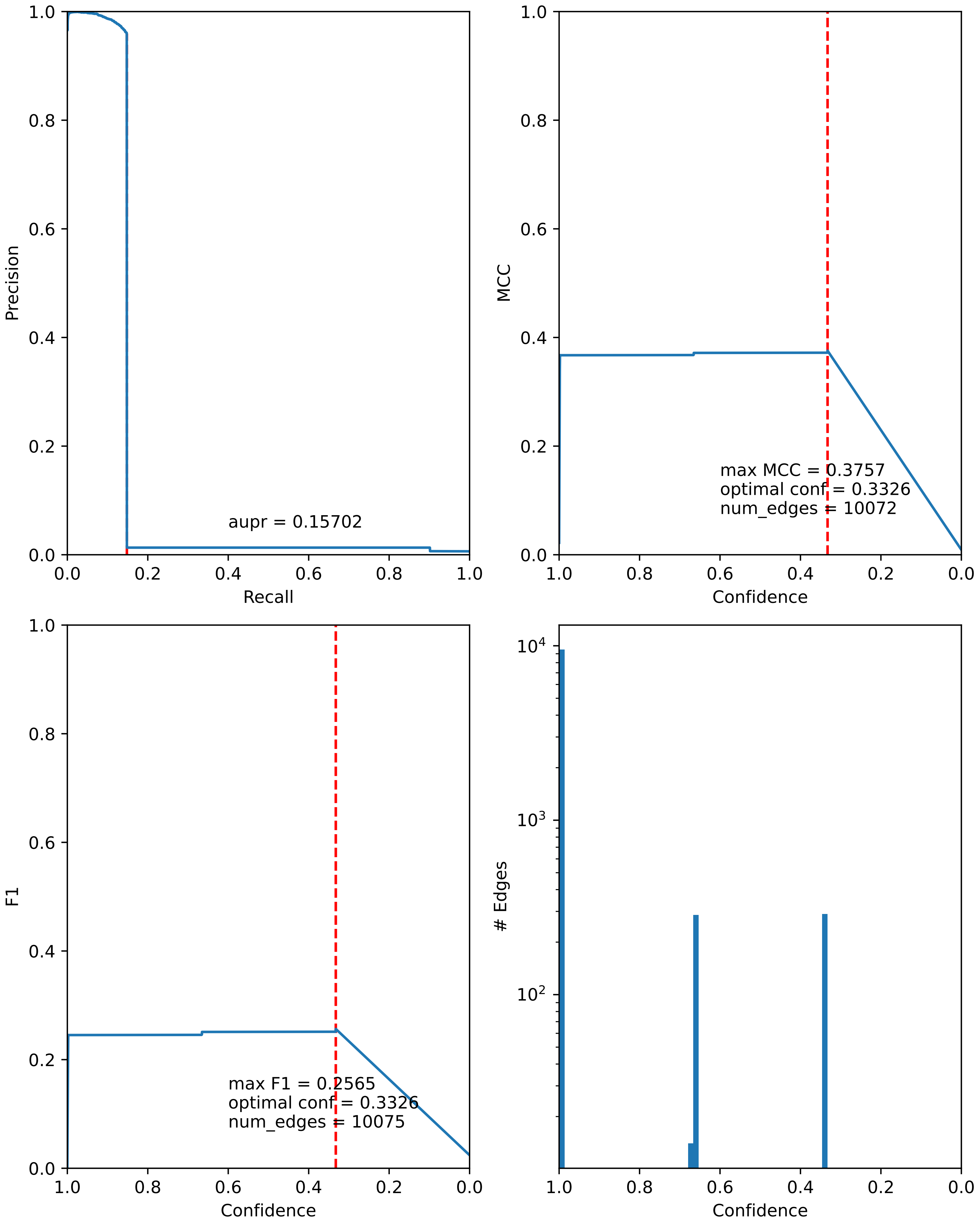

### combined_metrics.pdf

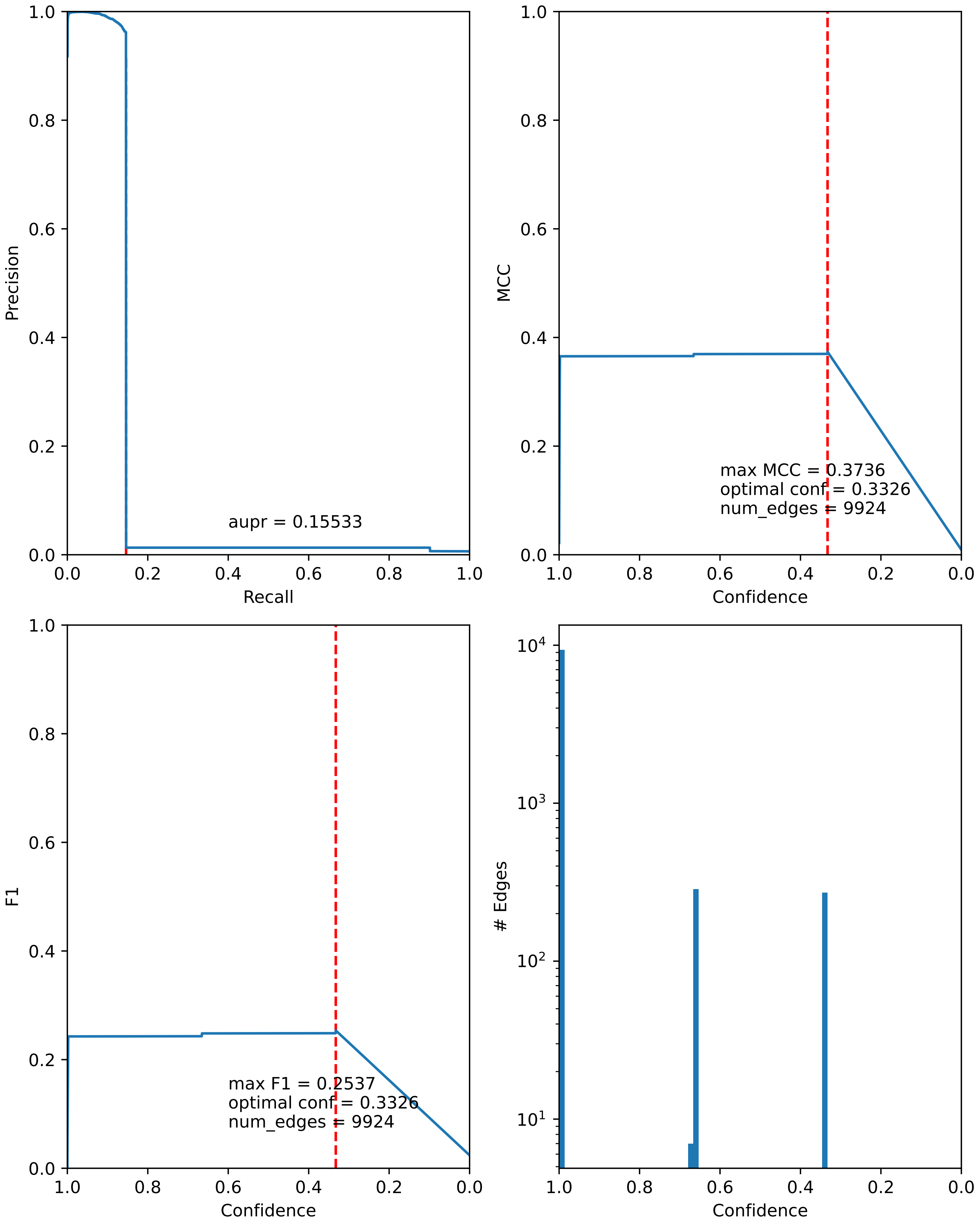

### combined_metrics.pdf

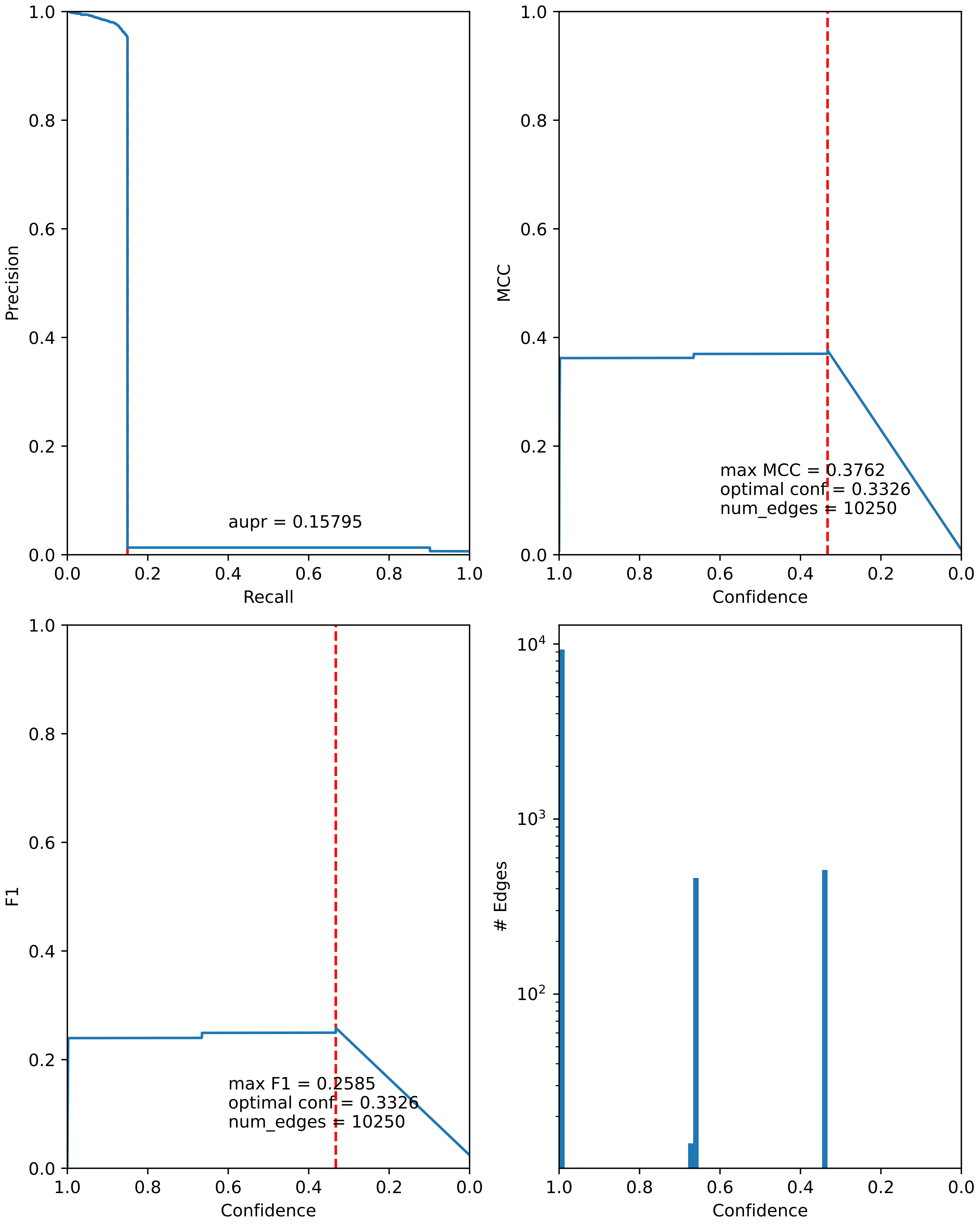

### combined_metrics.pdf

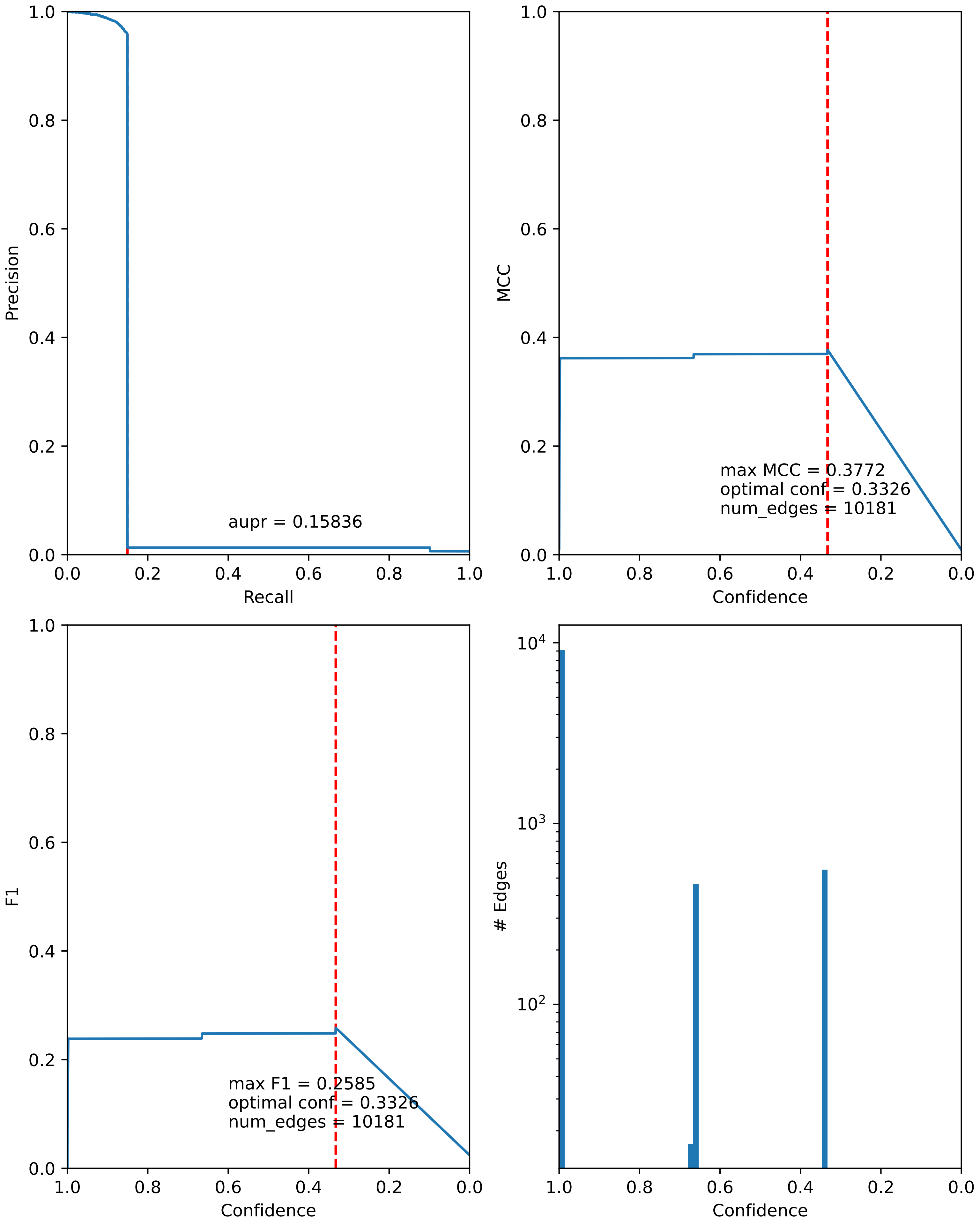

### combined_metrics.pdf

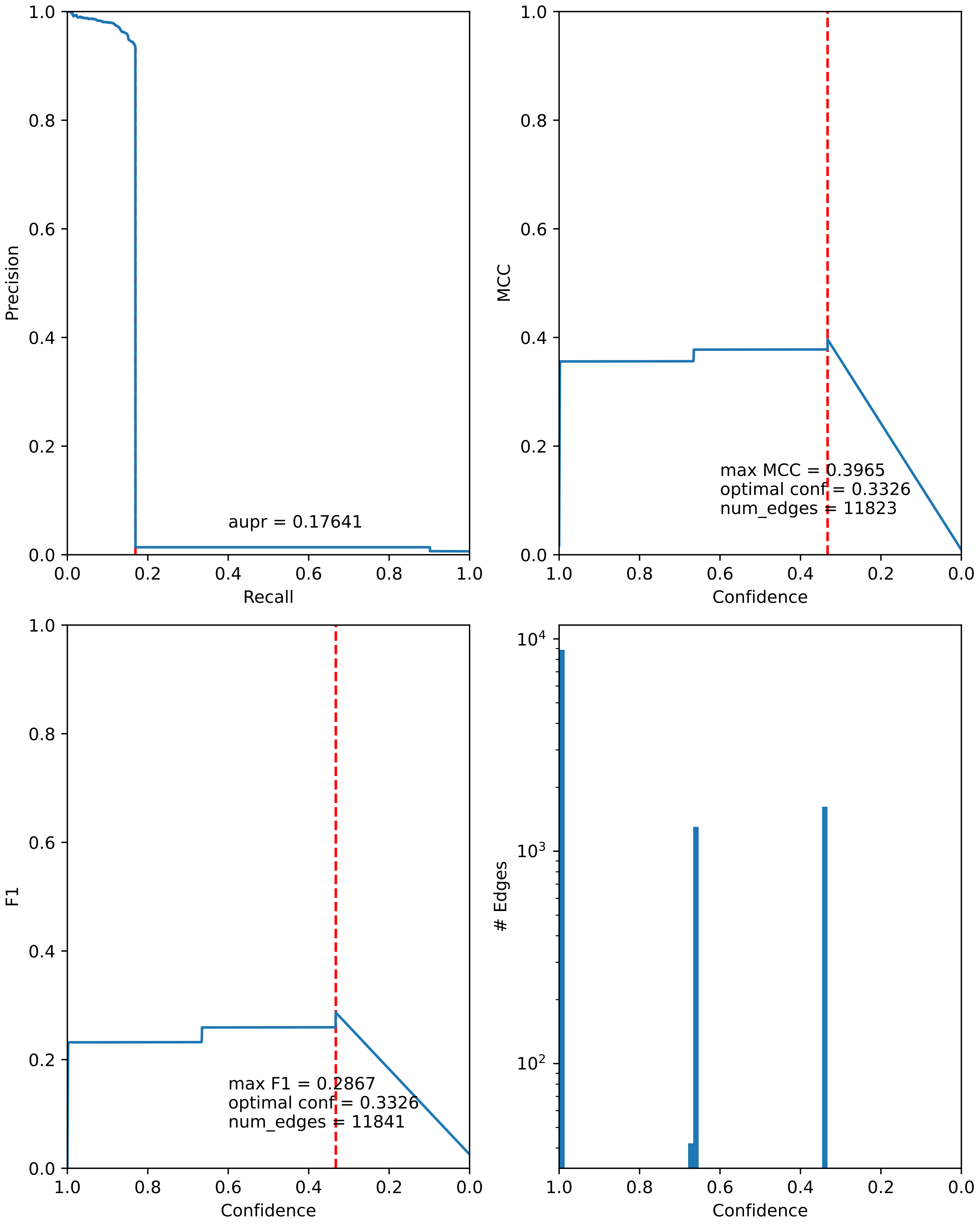

### combined_metrics.pdf

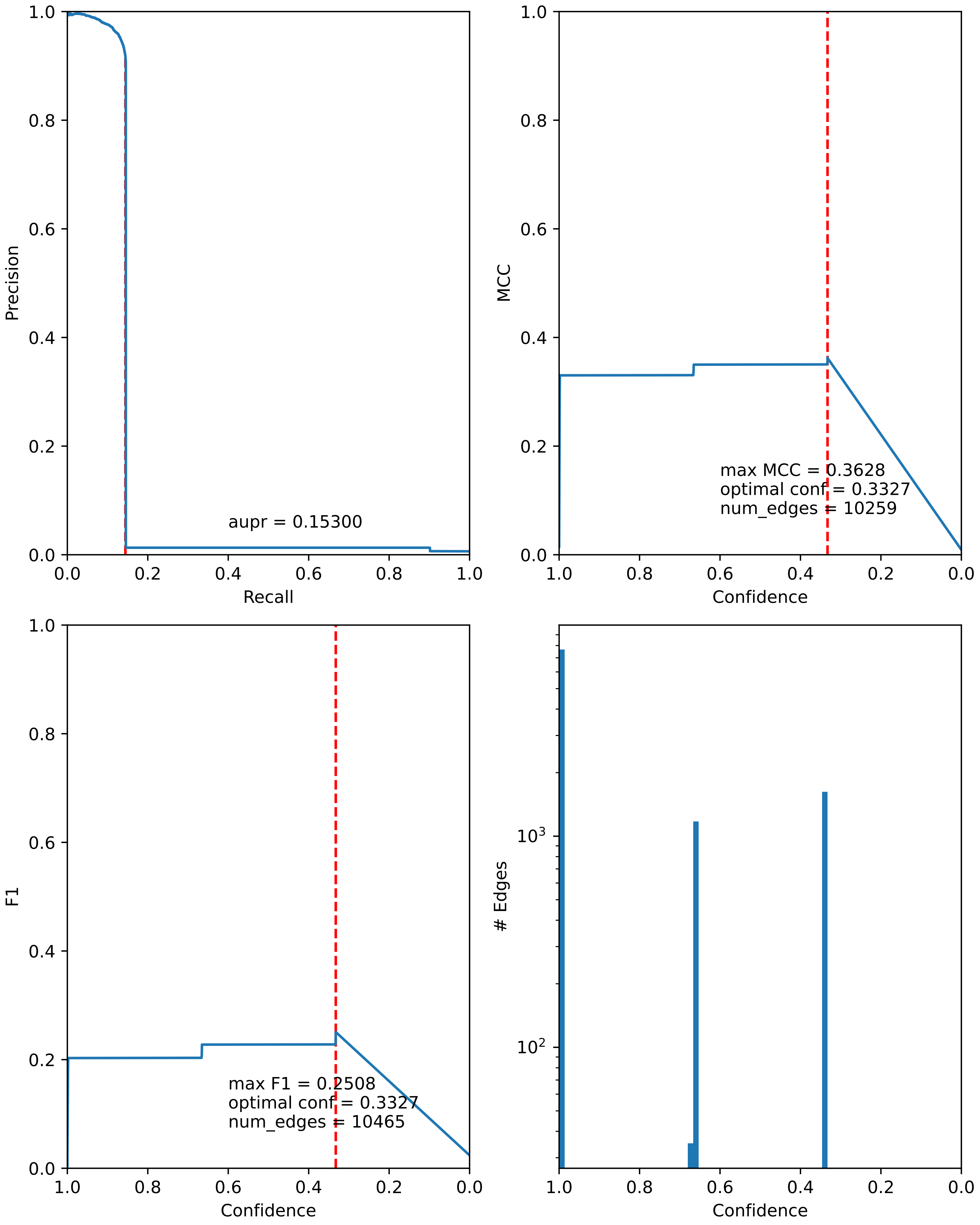

### combined_metrics.pdf

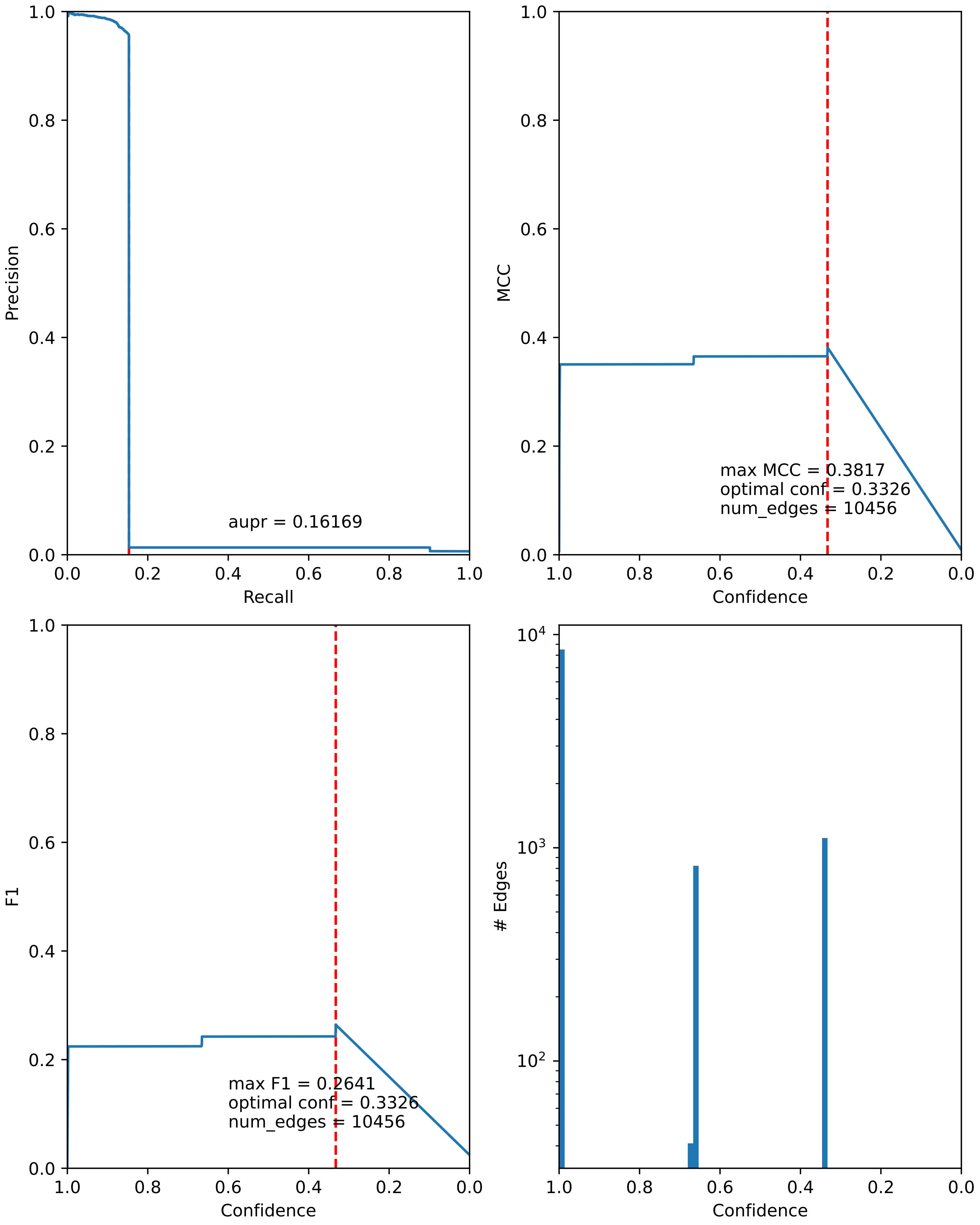

### combined_metrics.pdf

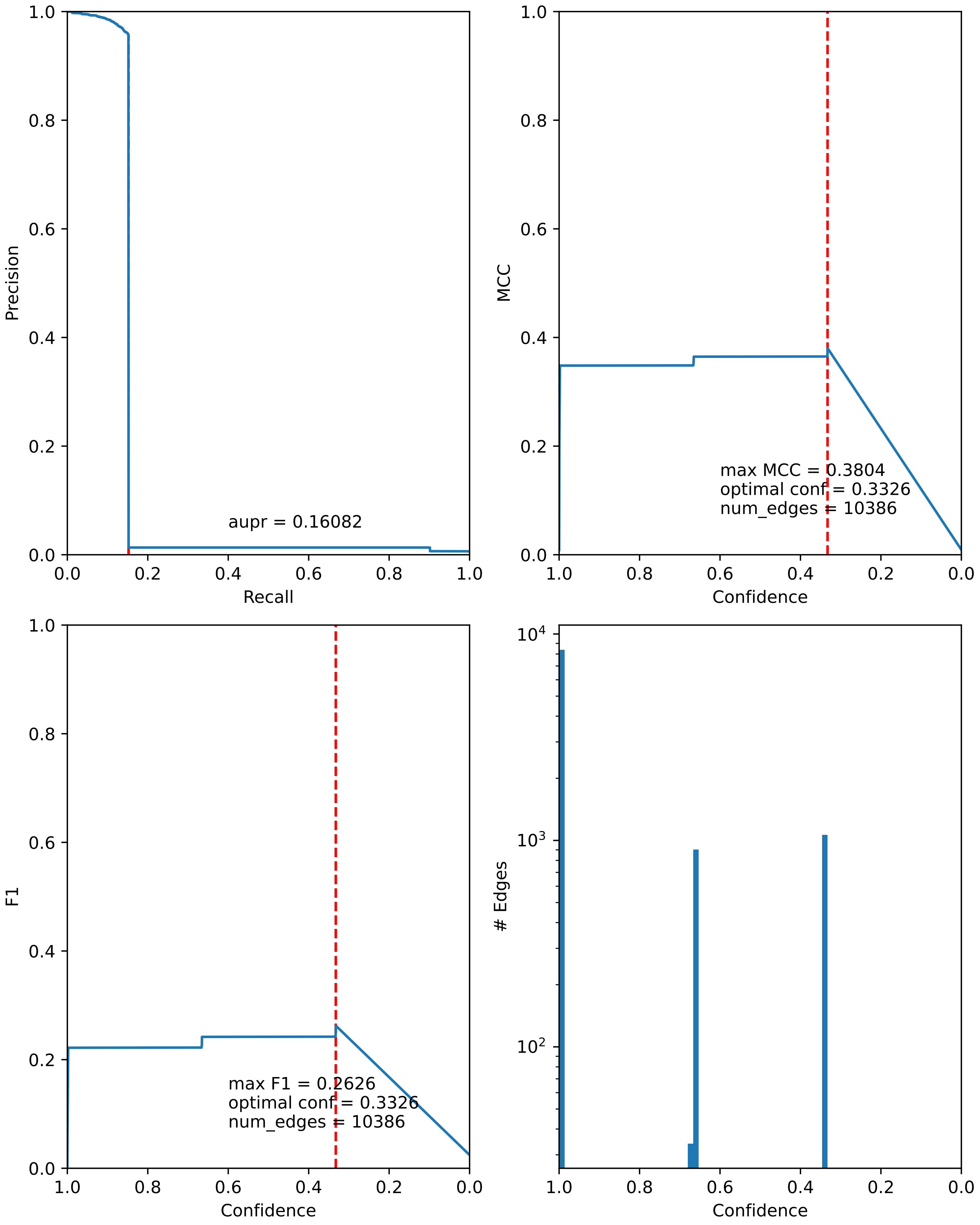

### combined_metrics.pdf

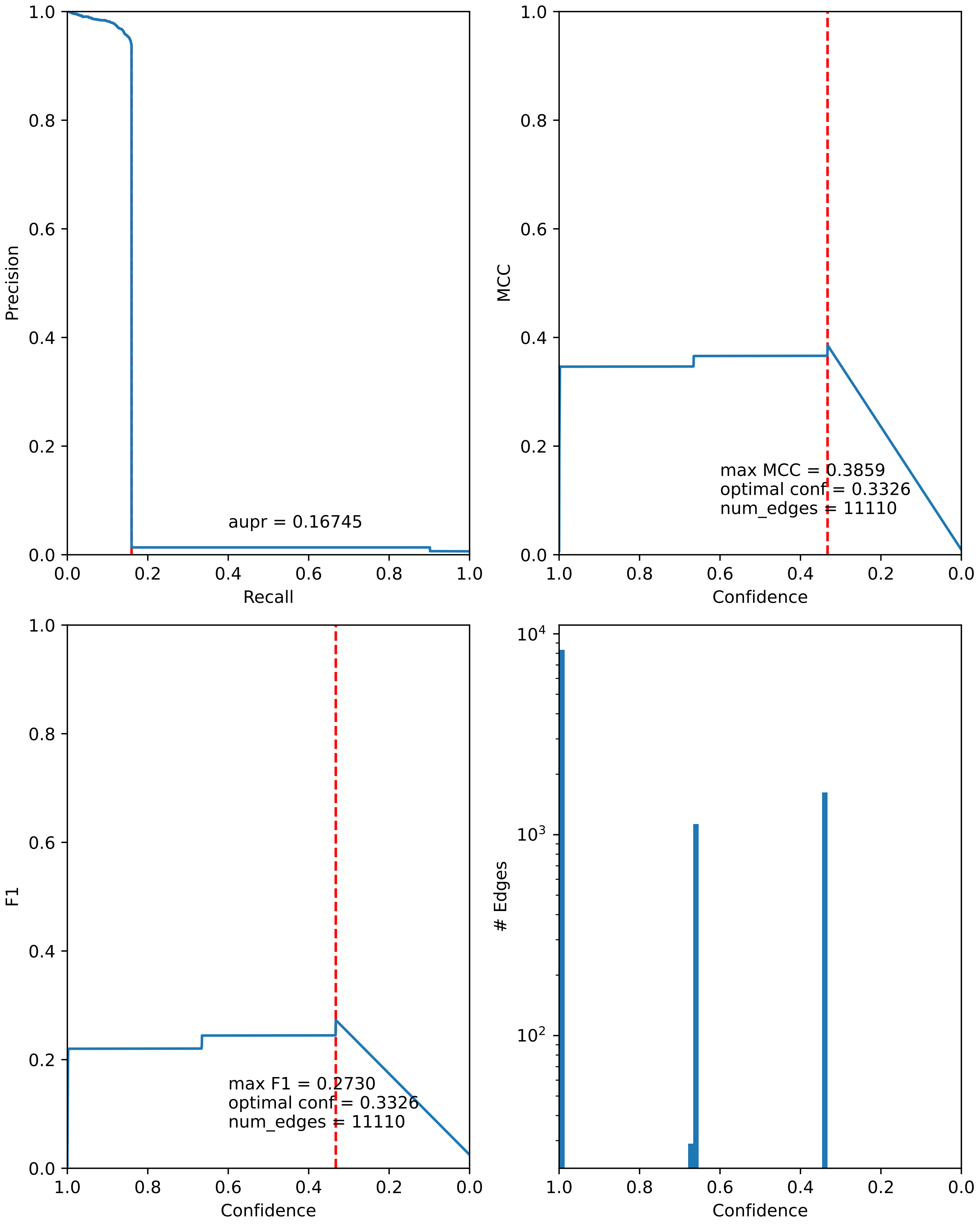

### combined_metrics.pdf

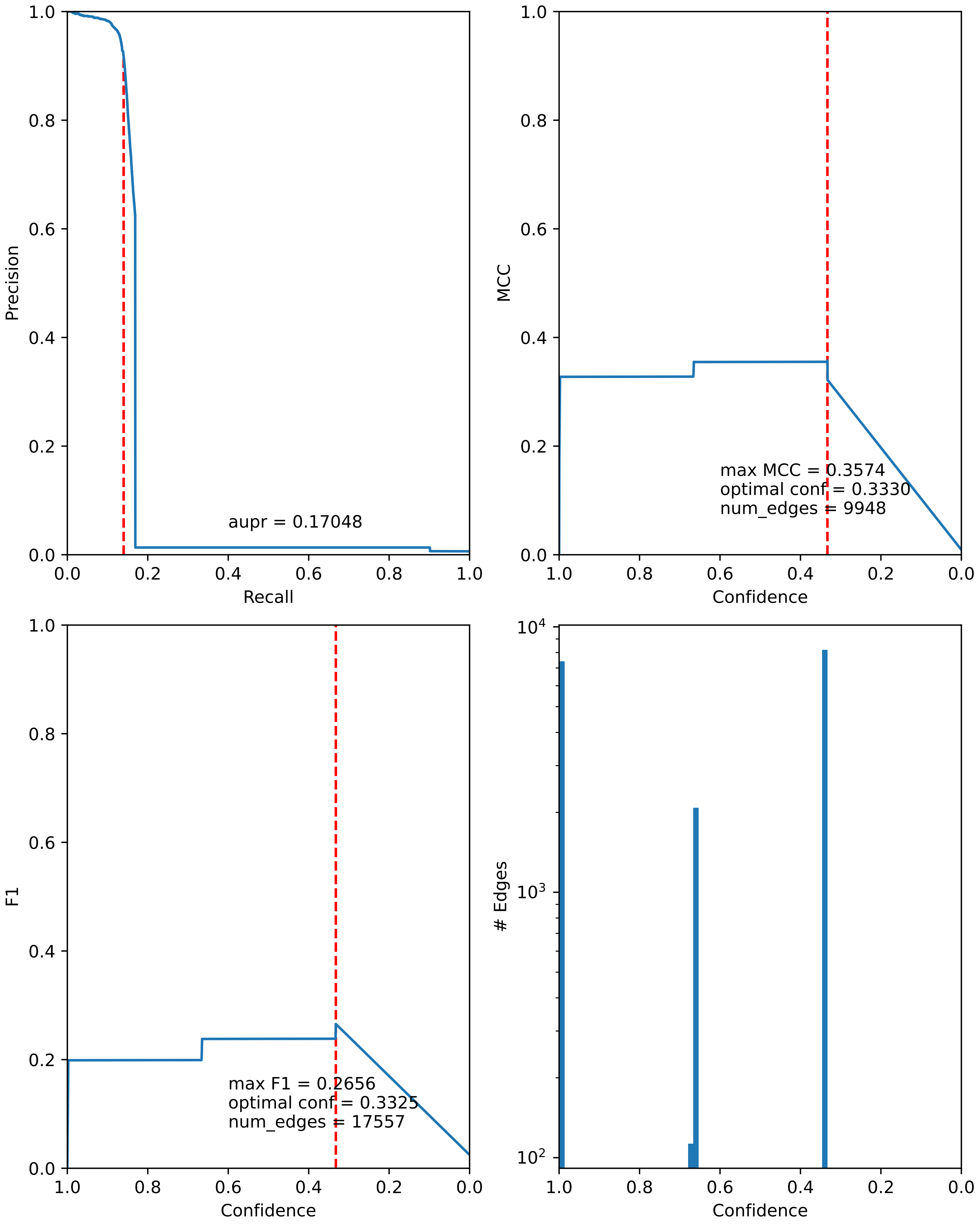

### combined_metrics.pdf

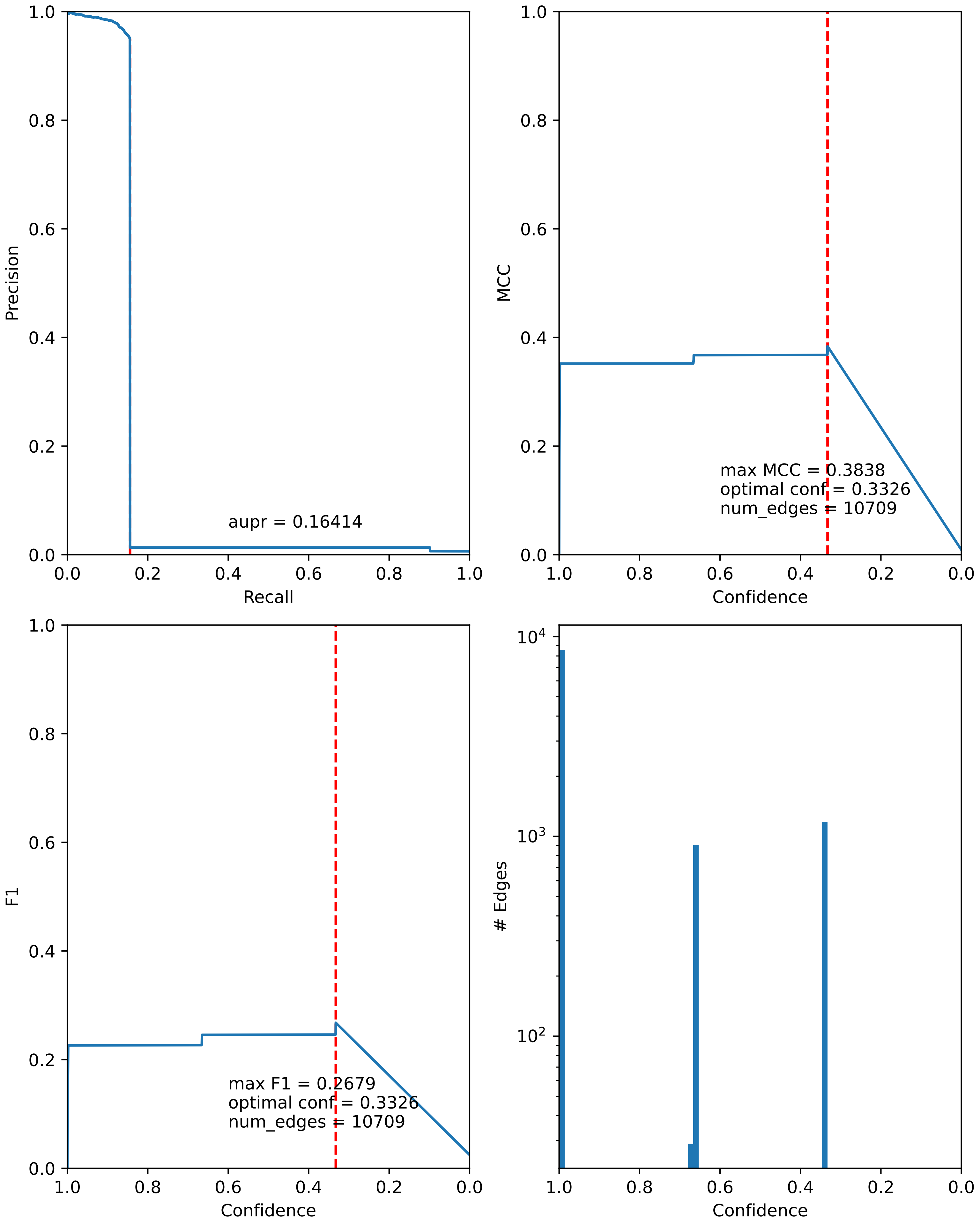

### combined_metrics.pdf

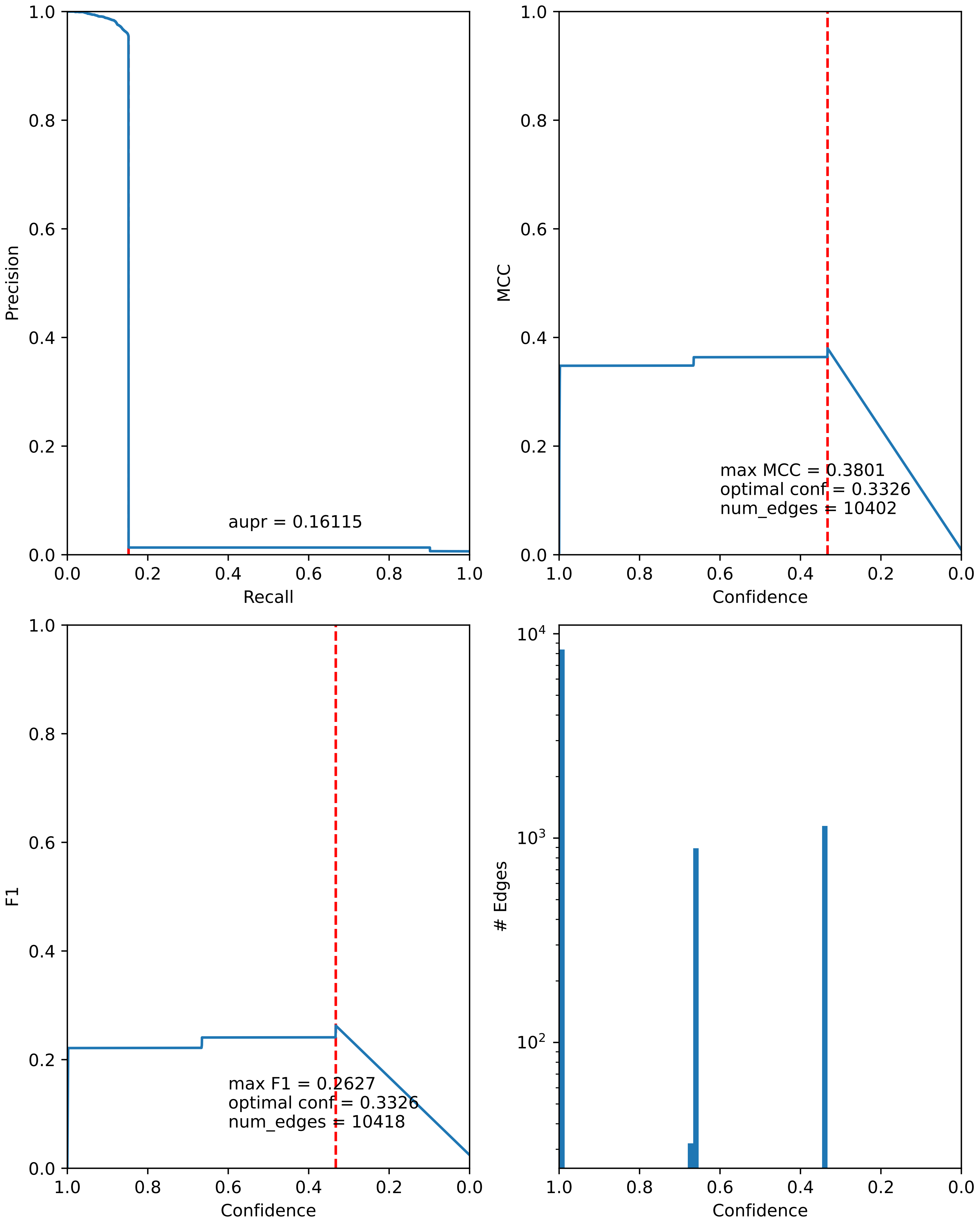

### combined_metrics.pdf

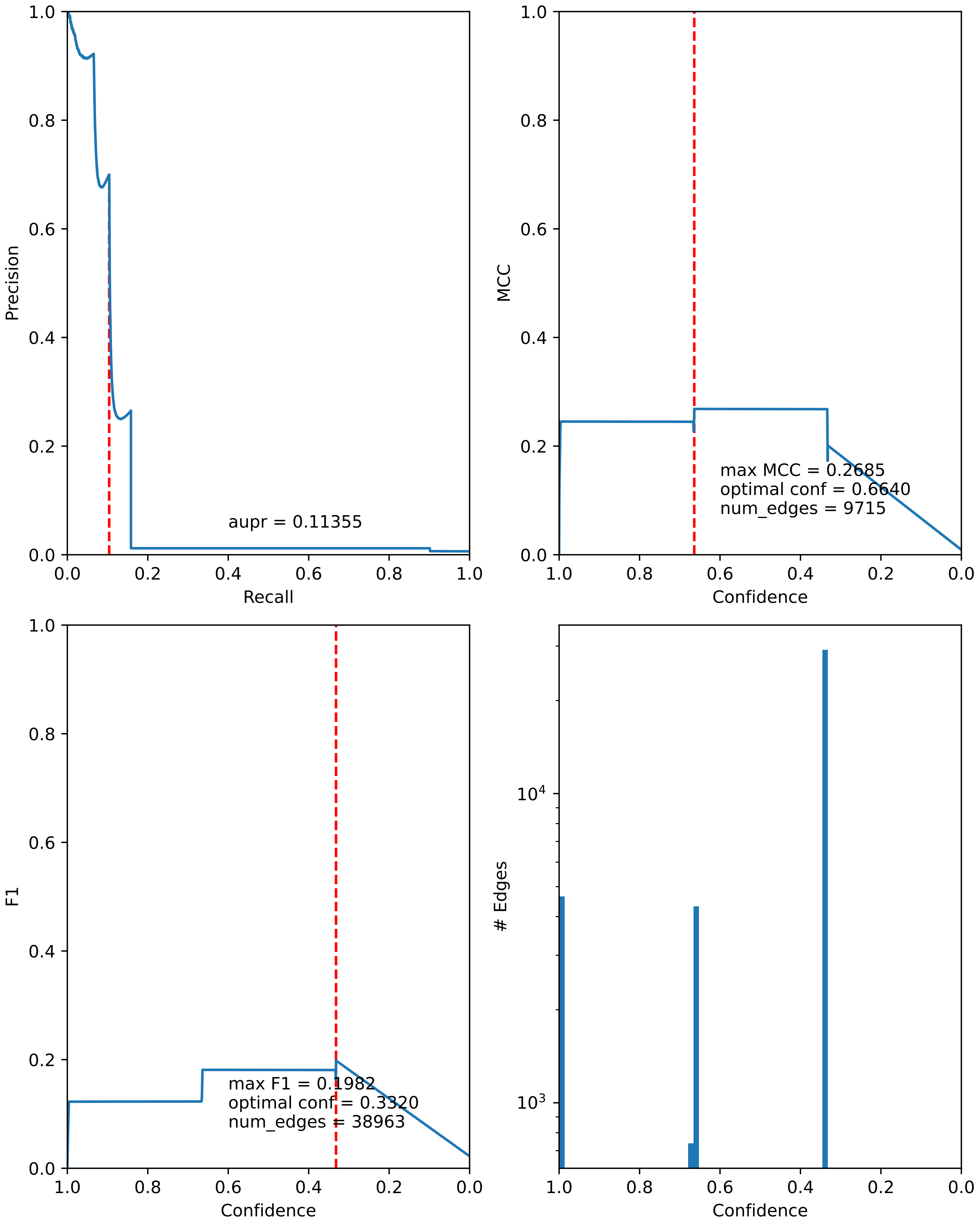

### combined_metrics.pdf

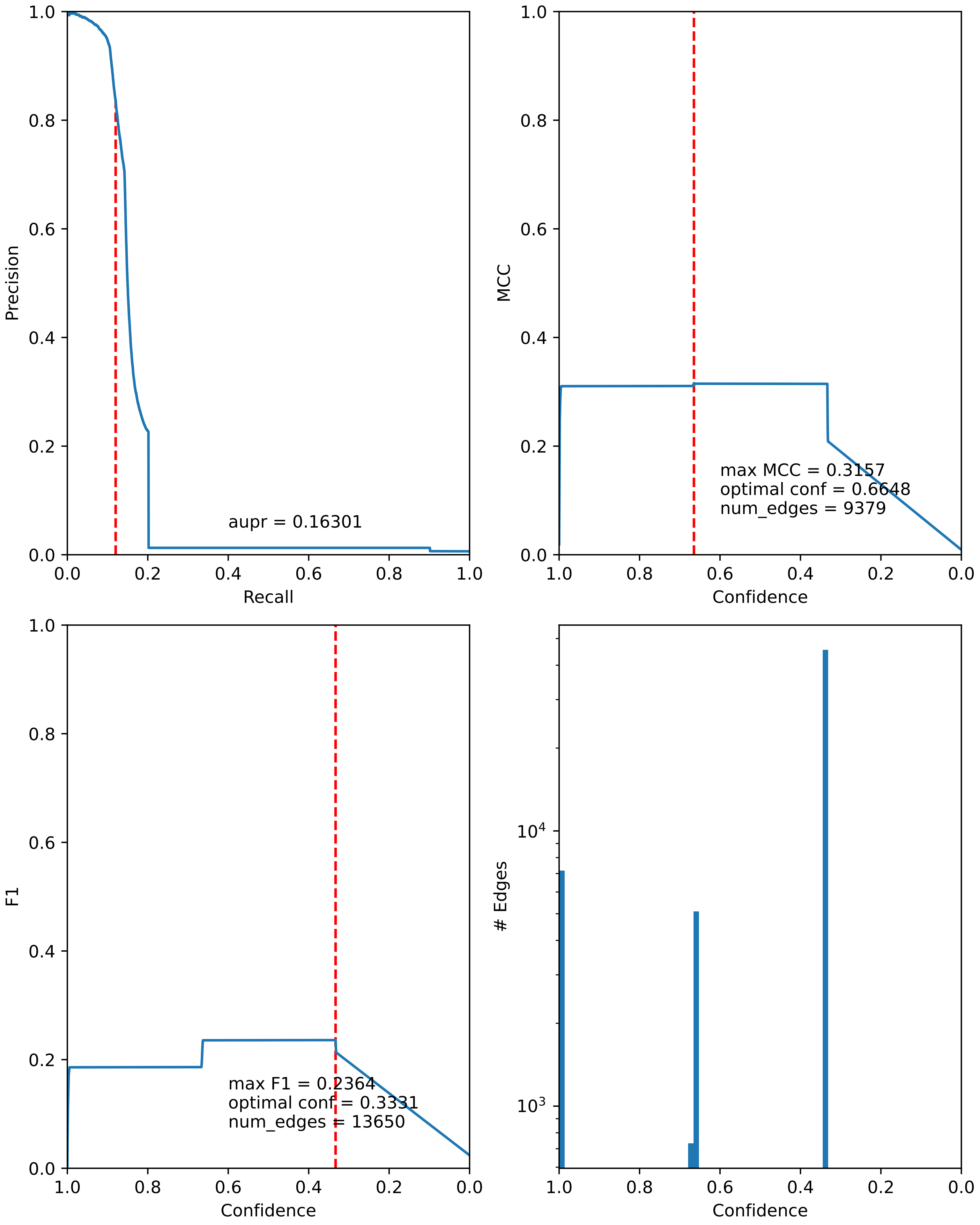

### combined_metrics.pdf

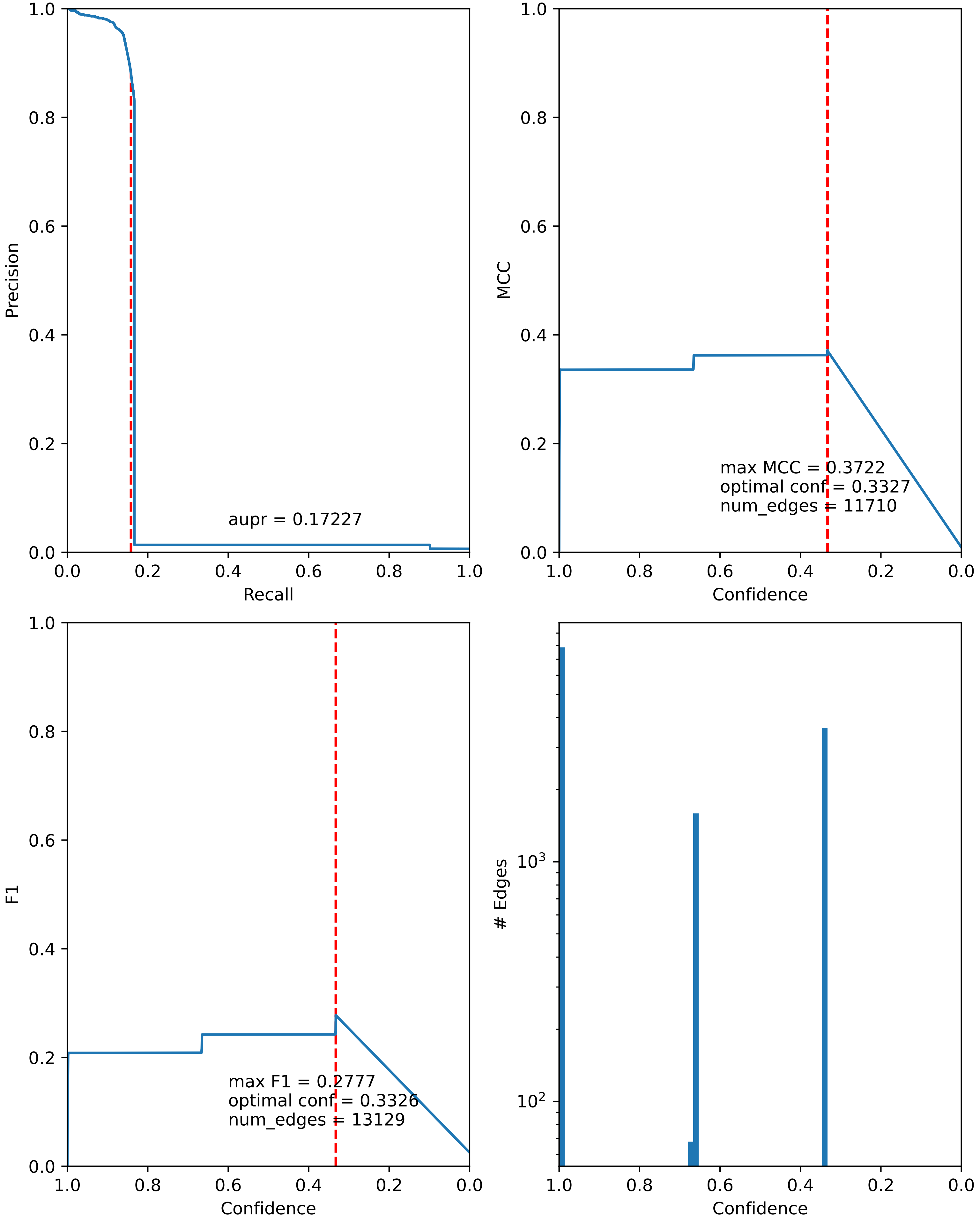

### combined_metrics.pdf

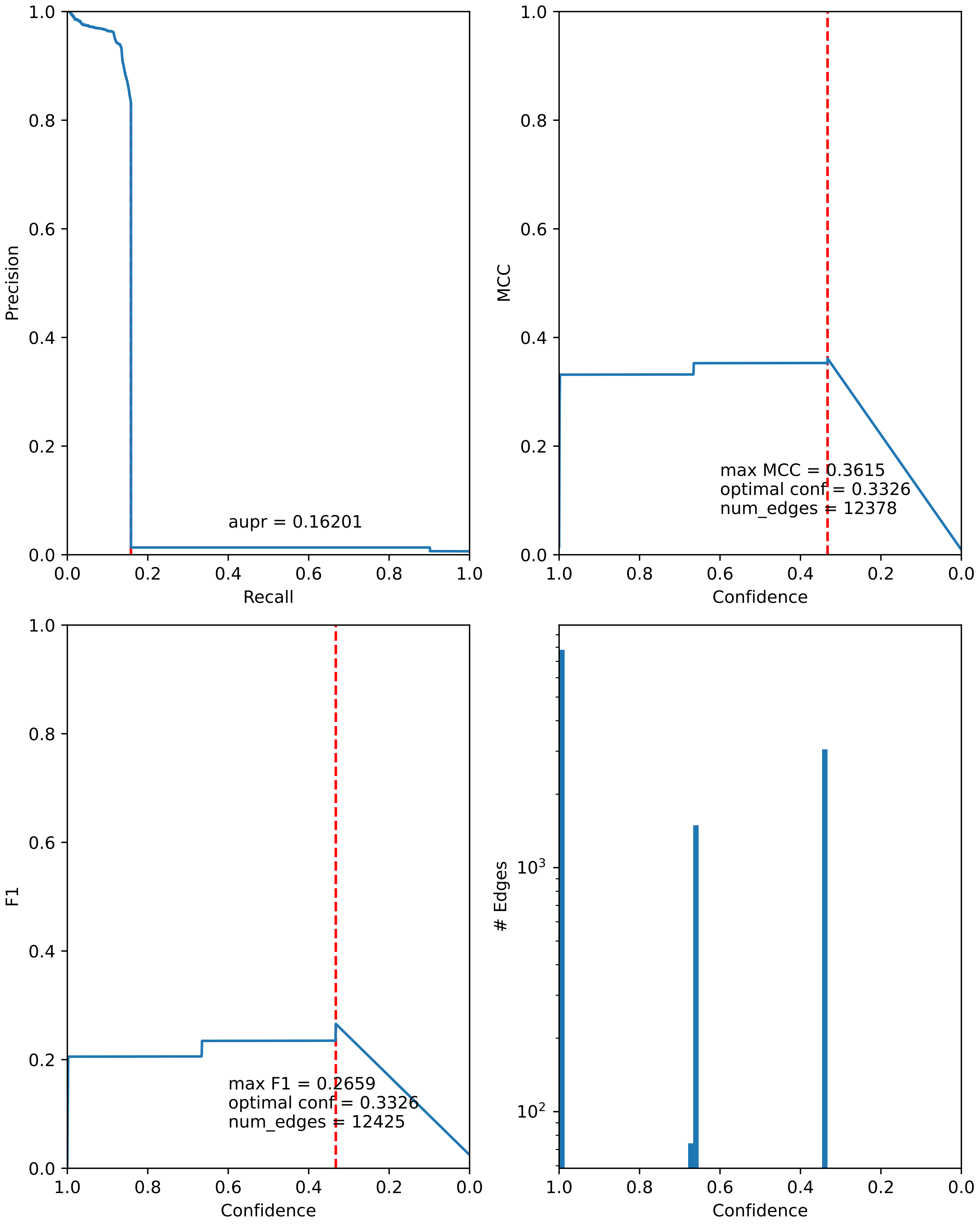

### combined_metrics.pdf

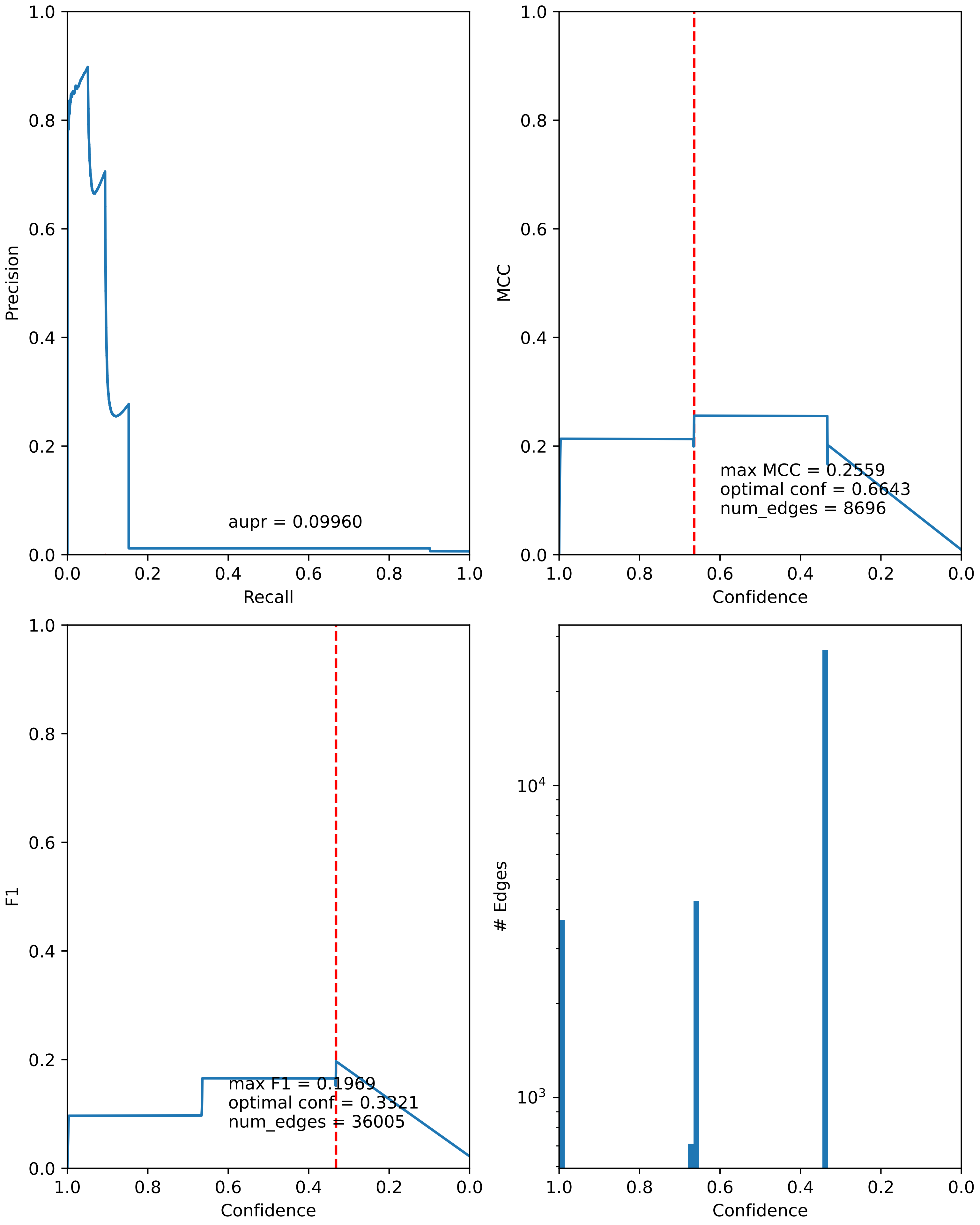

### combined_metrics.pdf

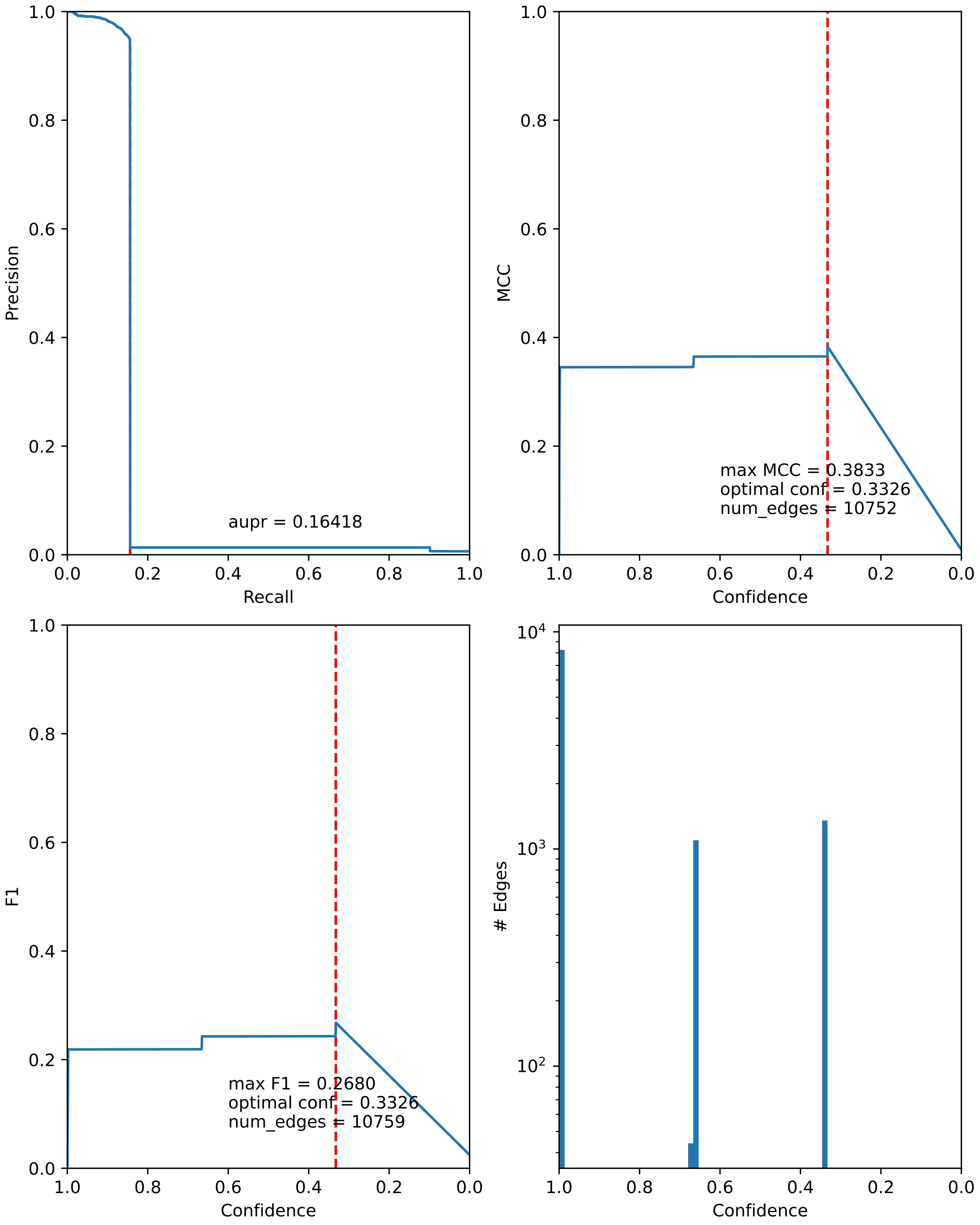

### combined_metrics.pdf

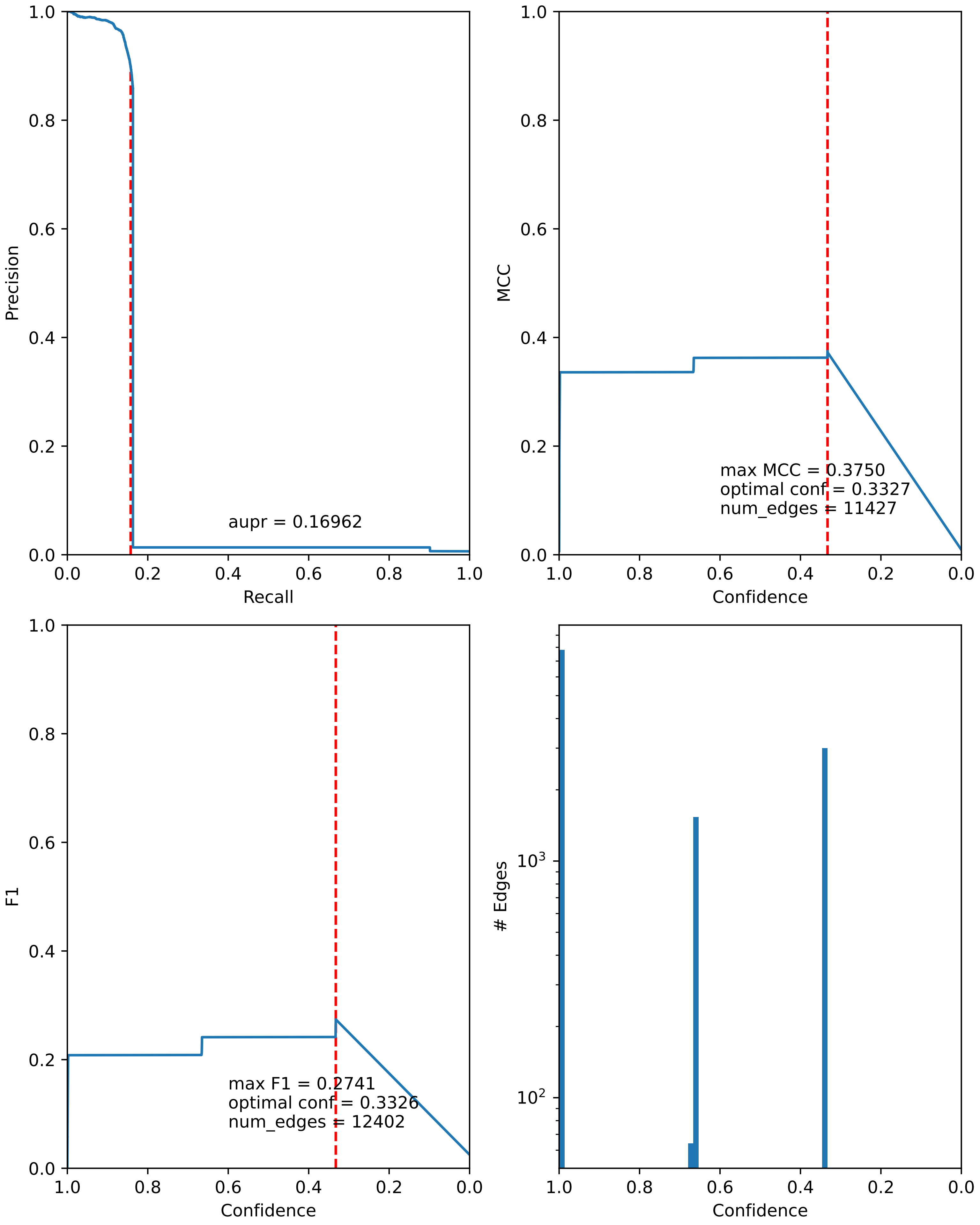

### combined_metrics.pdf

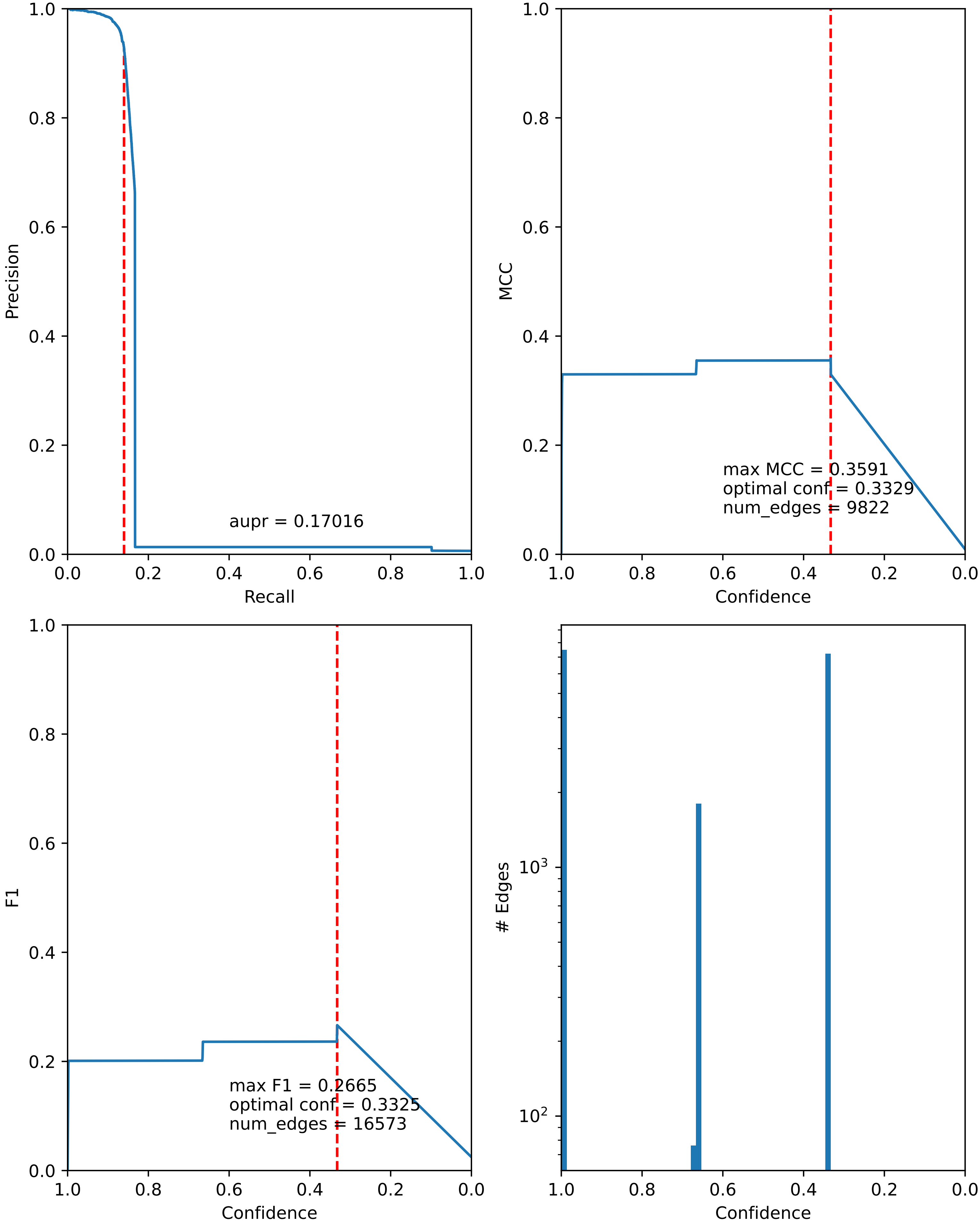

### combined_metrics.pdf

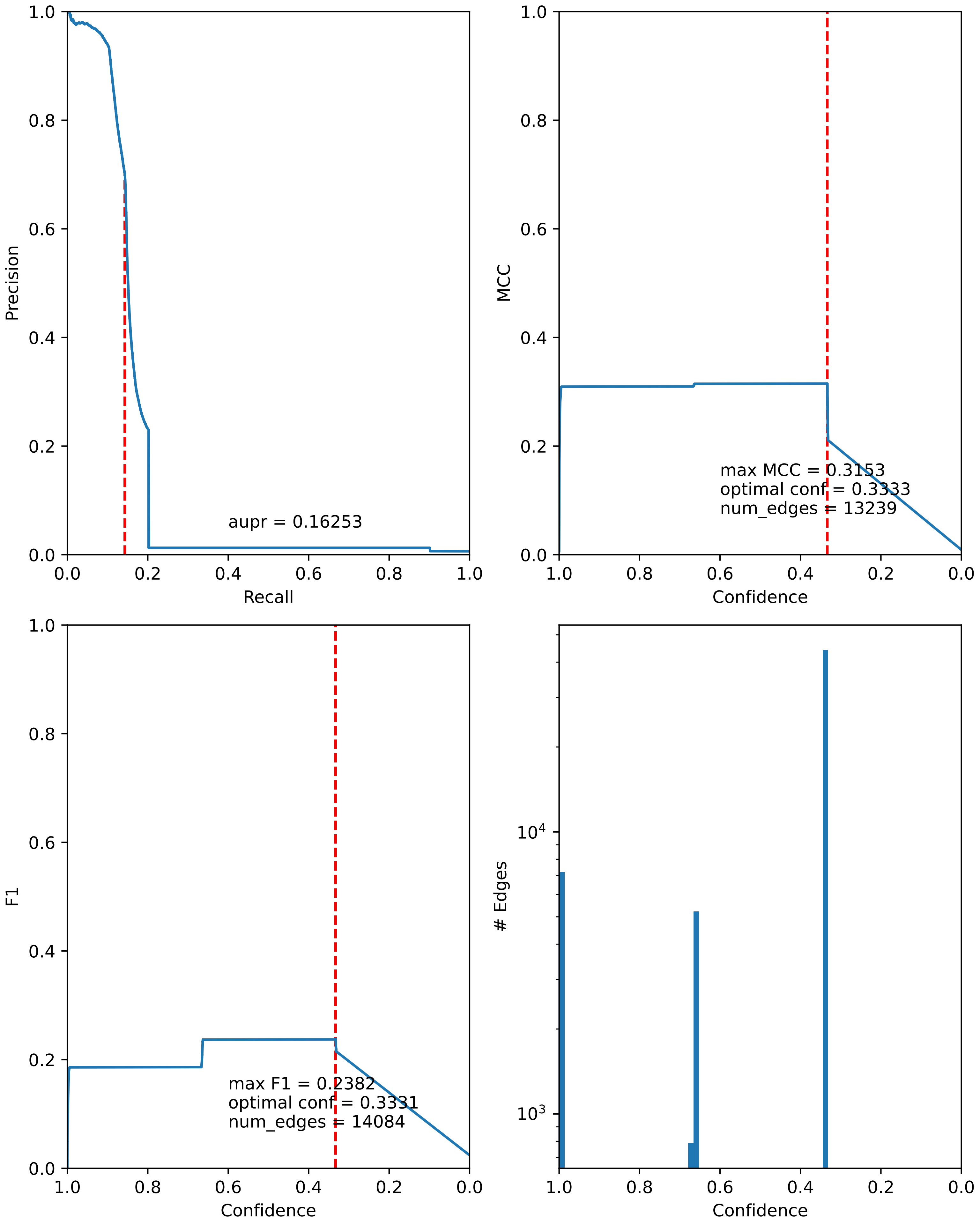

### combined_metrics.pdf

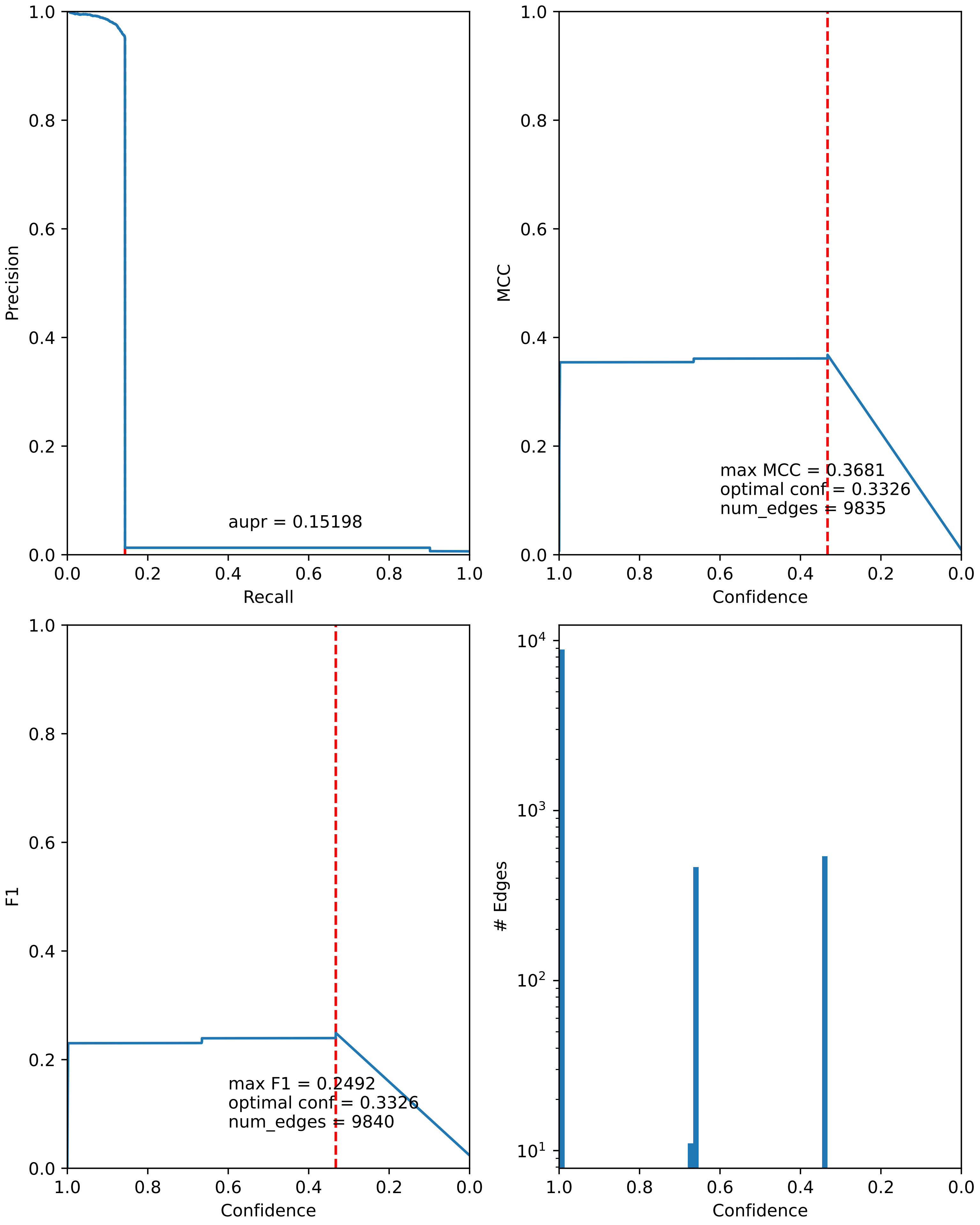

### combined_metrics.pdf

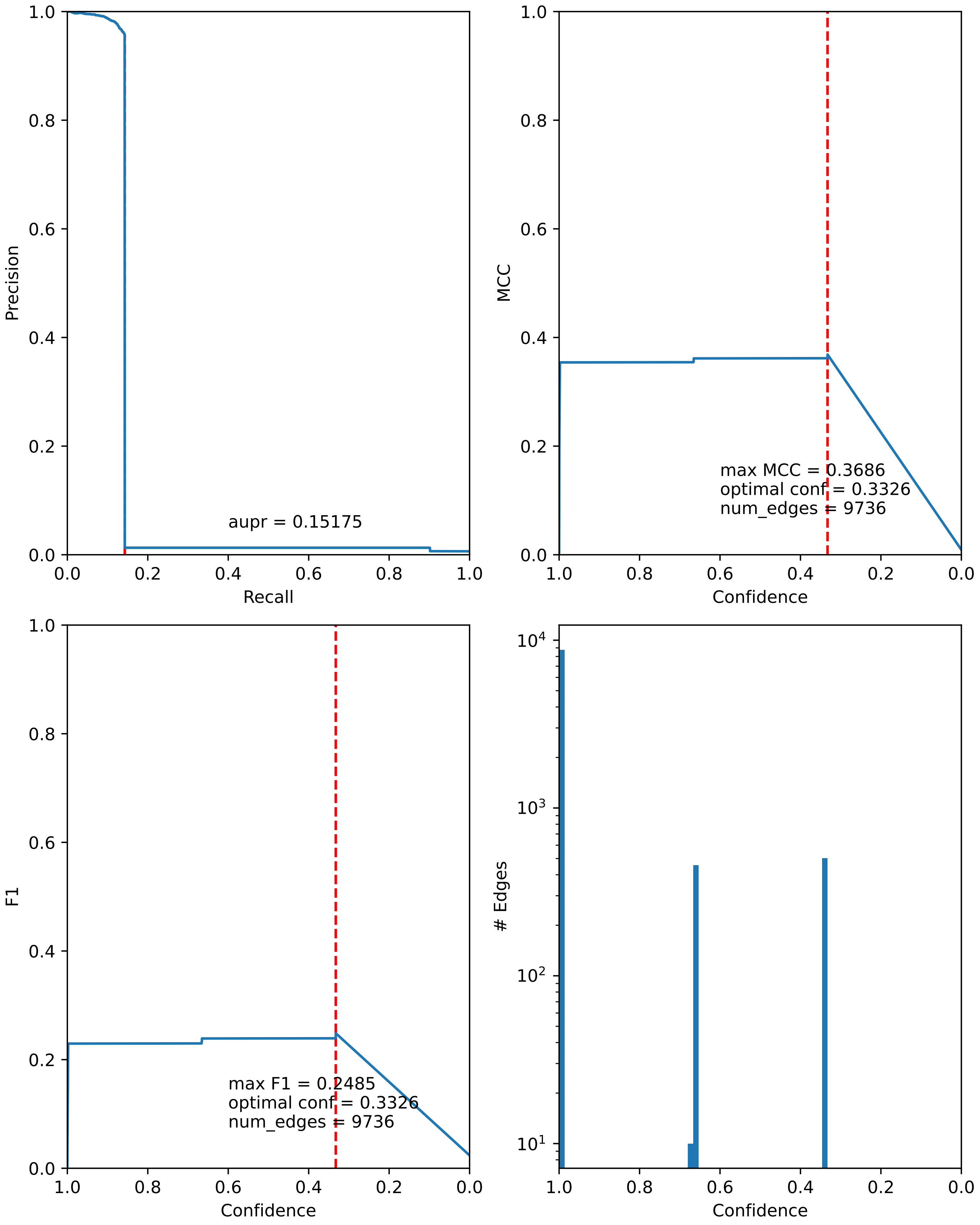

### combined_metrics.pdf

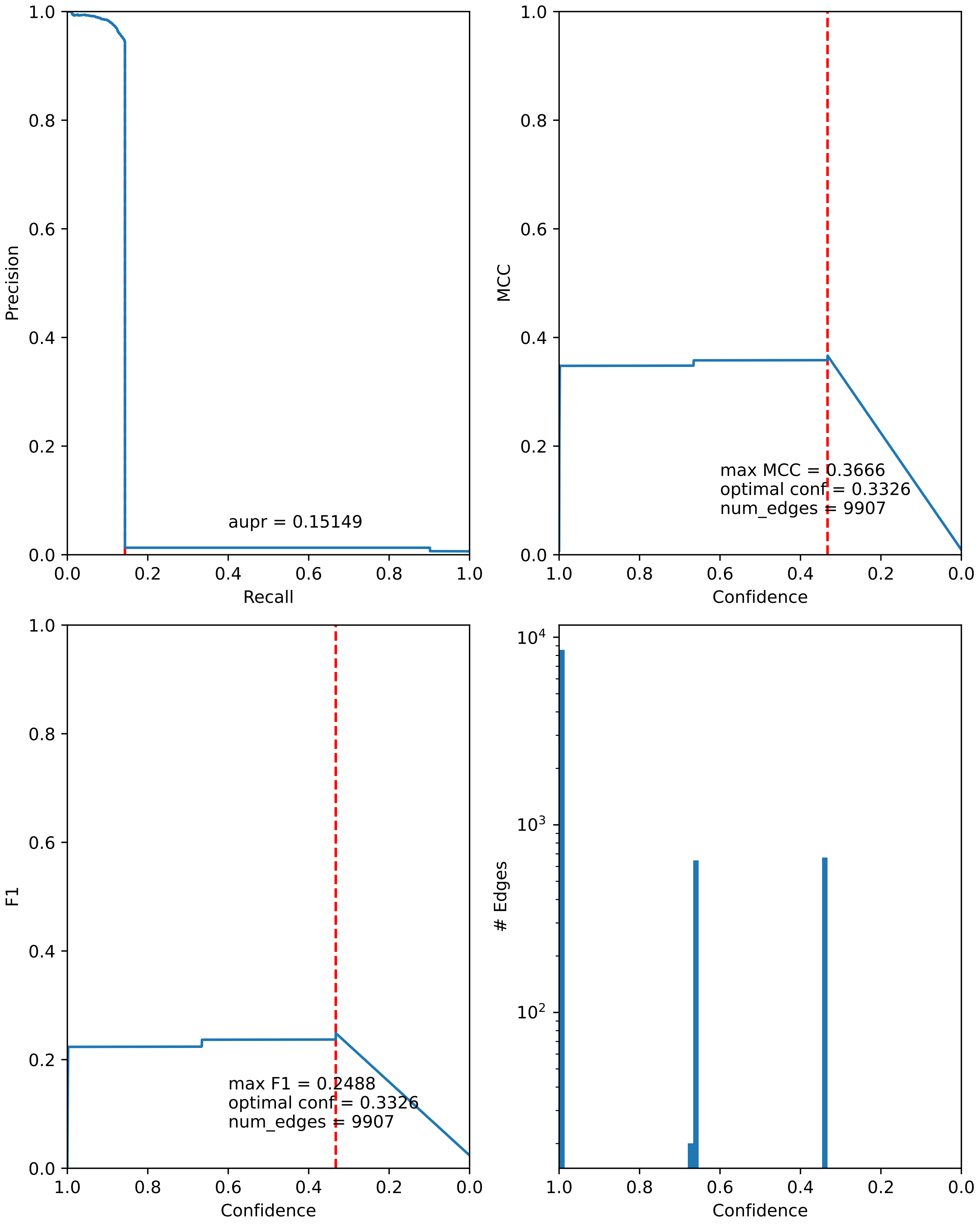

### combined_metrics.pdf

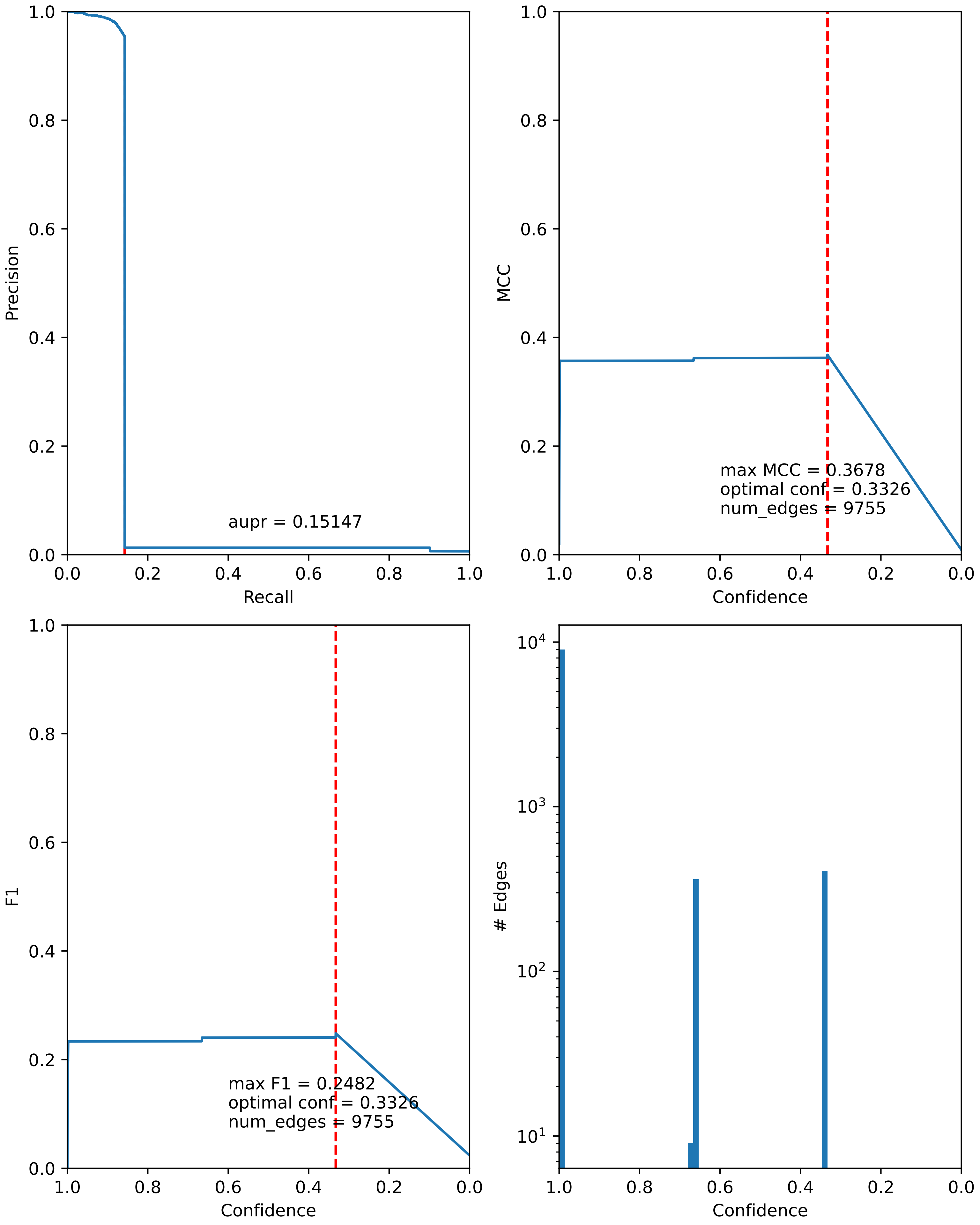

### combined_metrics.pdf

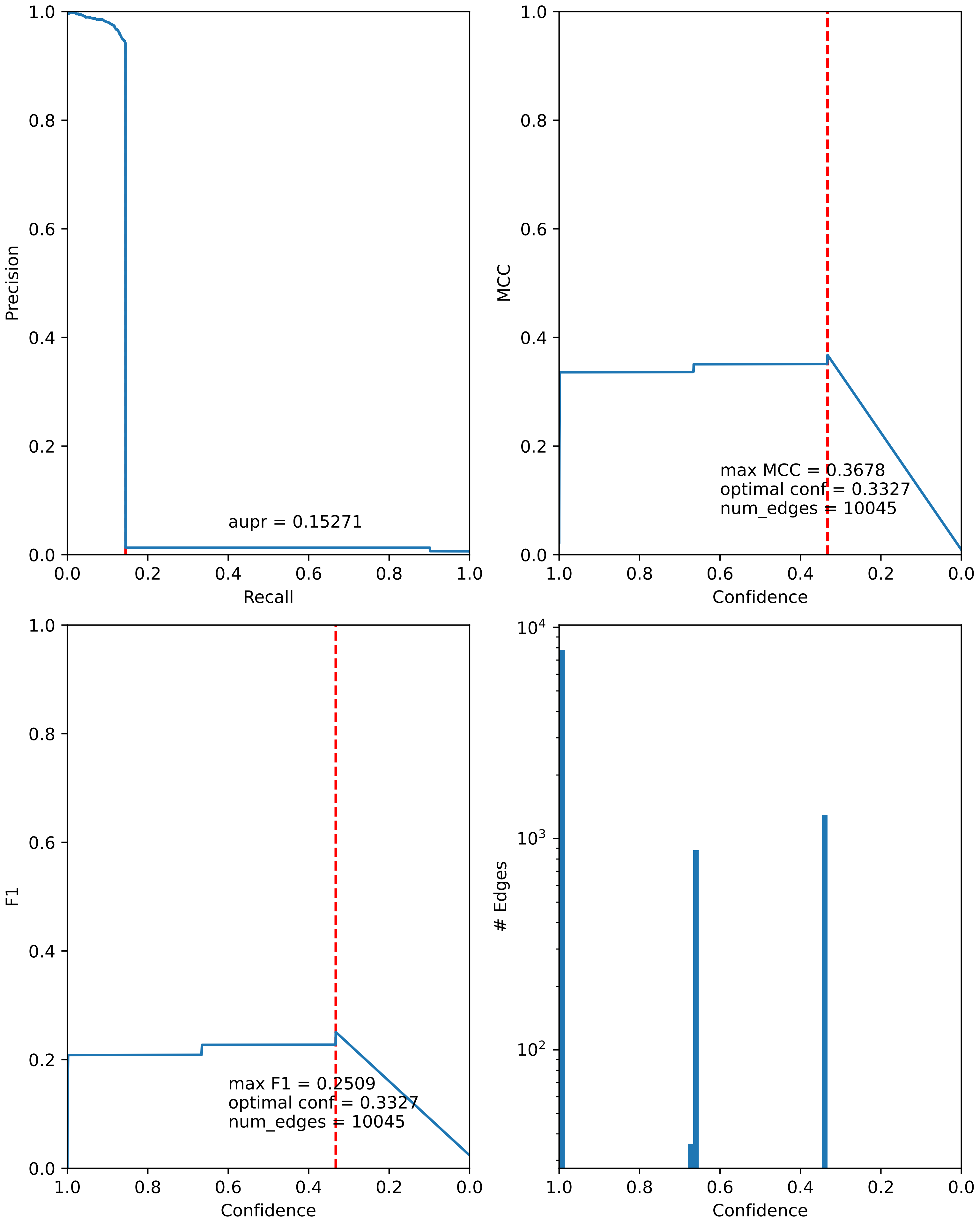

### combined_metrics.pdf

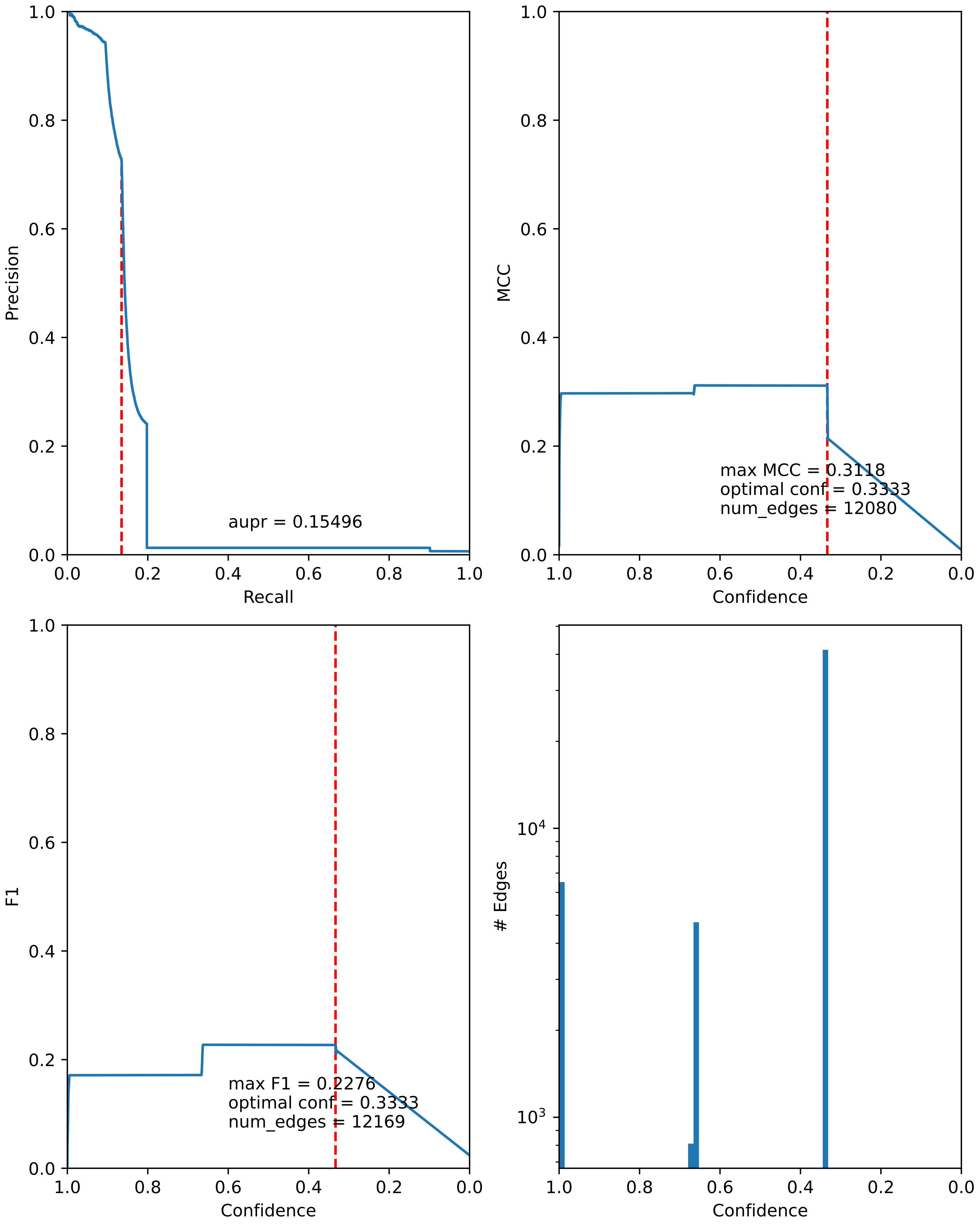

### combined_metrics.pdf

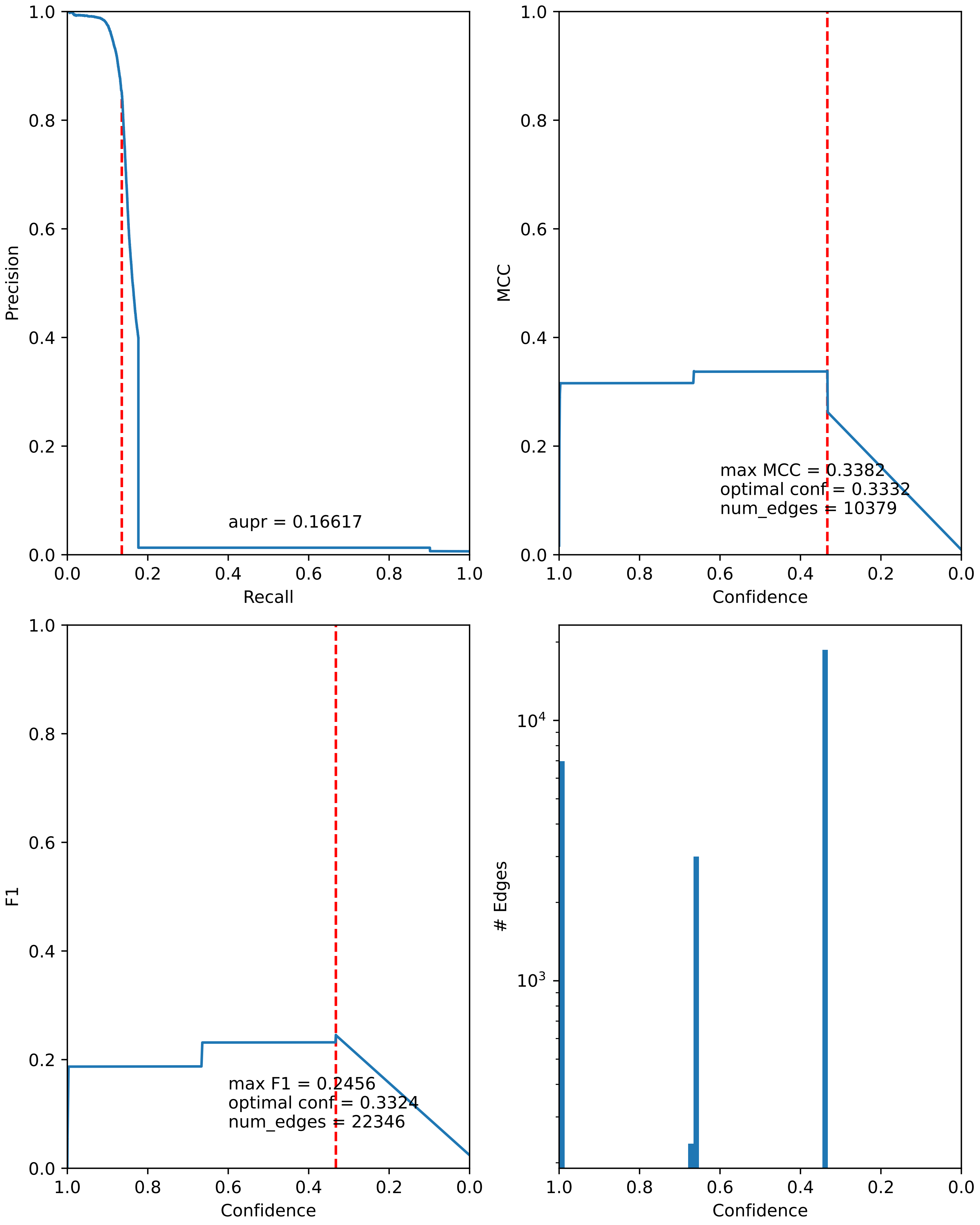

### combined_metrics.pdf

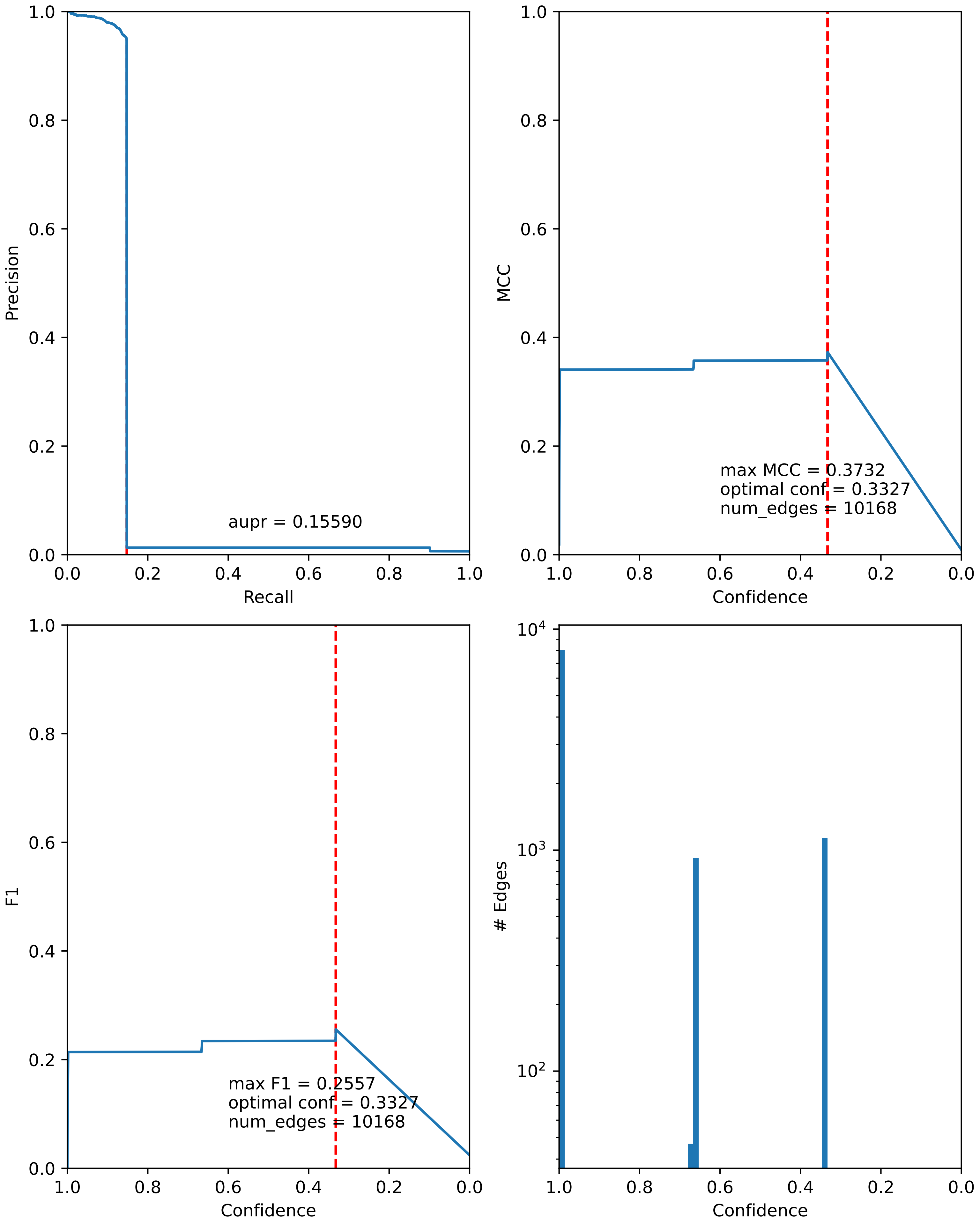

### combined_metrics.pdf

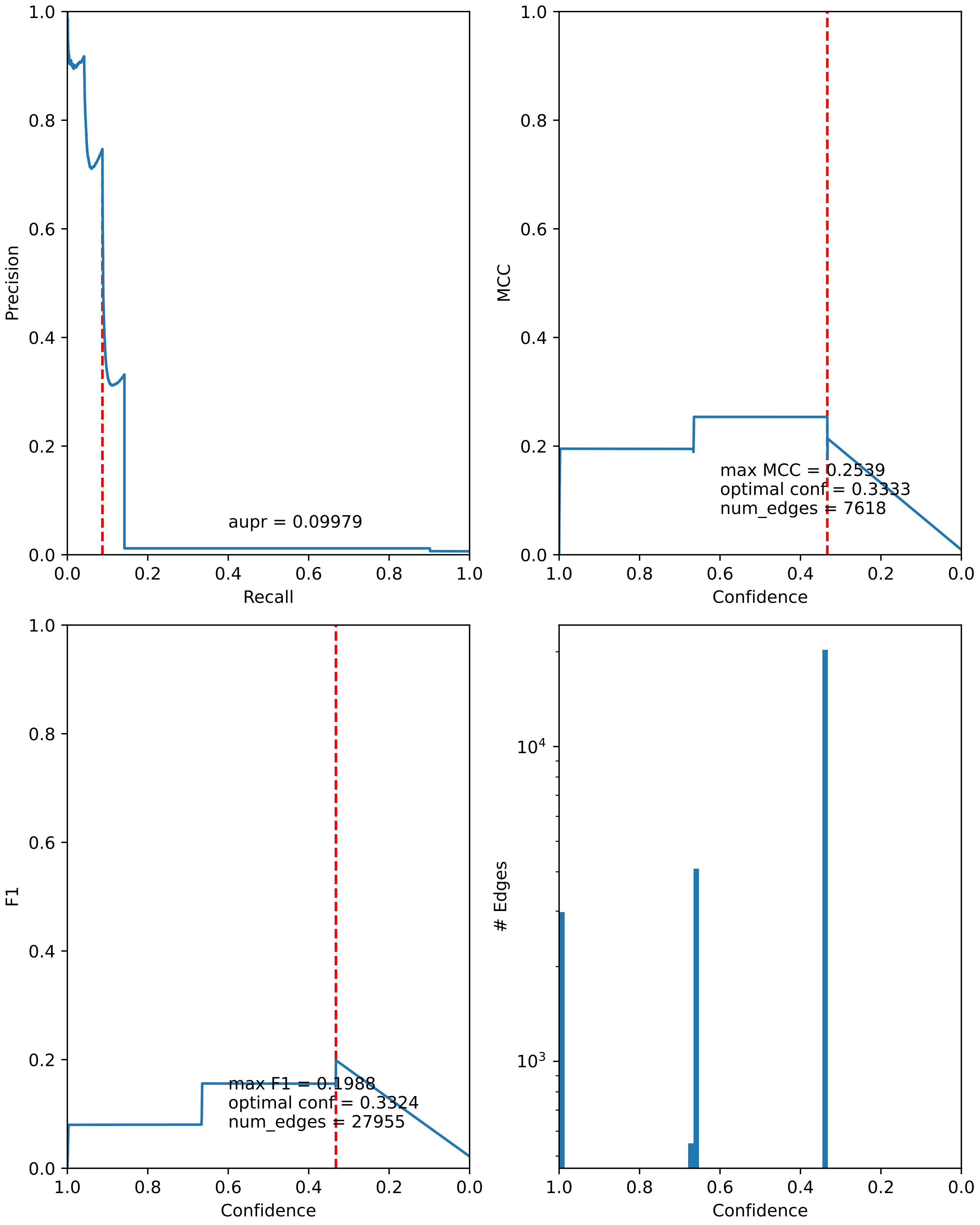

### combined_metrics.pdf

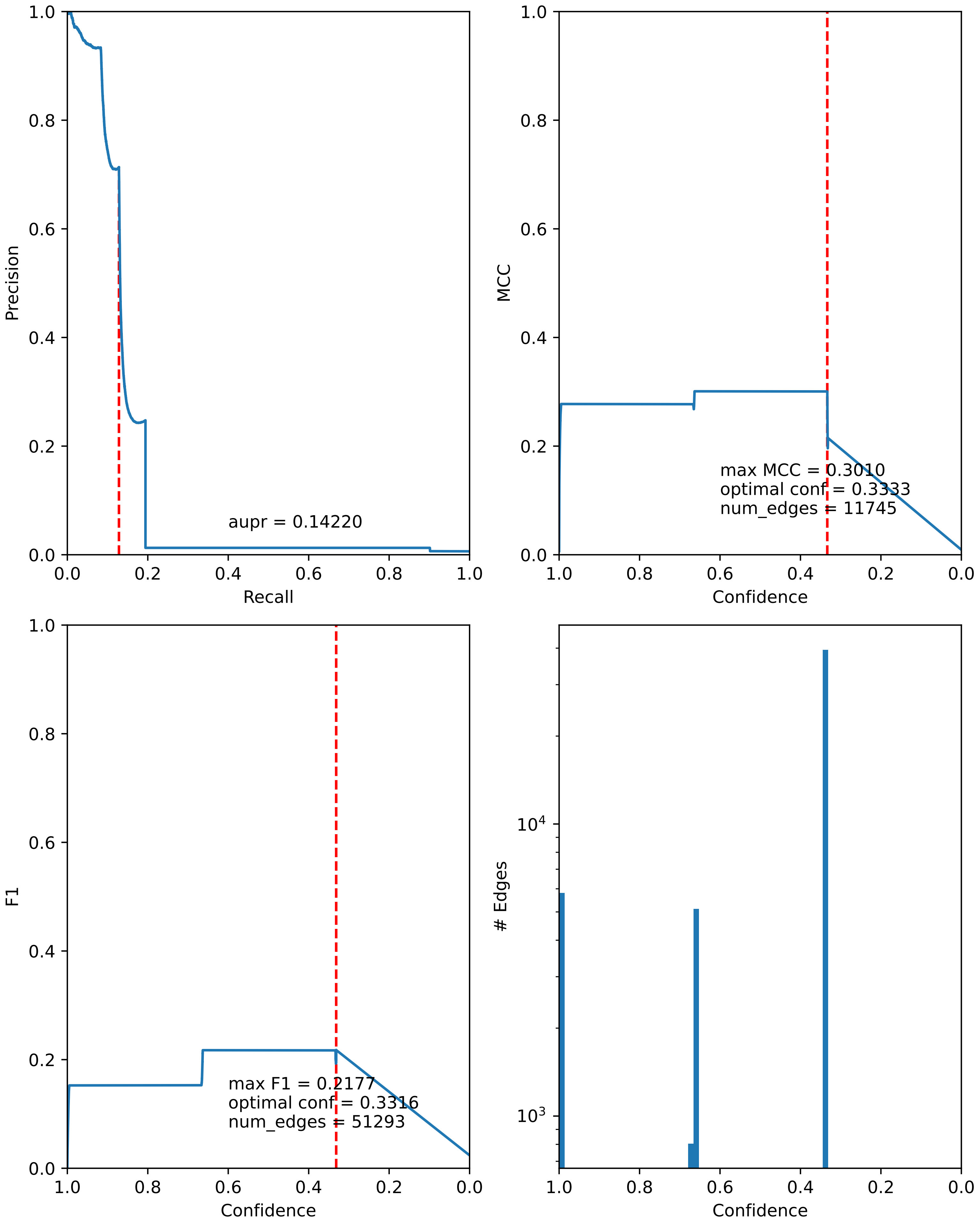
